# Genome wide variation in the Angolan Namib desert reveals unique Pre-Bantu ancestry

**DOI:** 10.1101/2023.02.16.528838

**Authors:** Sandra Oliveira, Anne-Maria Fehn, Beatriz Amorim, Mark Stoneking, Jorge Rocha

## Abstract

The populations of the Angolan Namib Desert have been largely neglected in previous surveys of the genomic landscape of southern Africa. Although at present the Namib is culturally dominated by Southwest Bantu-speaking cattle-herders, the region exhibits an extraordinary ethnographic diversity which includes an array of semi-nomadic peoples whose subsistence strategies fall outside the traditional division between foraging and food production and can thus be referred to as “peripatetic”. Among these small-scale populations are the last speakers of the Kwadi branch of the Khoe-Kwadi language family associated with the introduction of pastoralism into southern Africa (Kwepe), as well as a range of groups whose origins remain enigmatic (Kwisi, Twa and Tjimba). Using genome-wide data from 208 individuals belonging to nine ethnically diverse groups from the Angolan Namib and adjacent areas (Kwepe, Kwisi, Twa, Tjimba, !Xun, Kuvale, Himba, Nyaneka, Ovimbundu) in combination with published data from other regions of Africa, we reconstruct in detail the histories of contact emerging from pre-historic migrations to southern Africa and show that peripatetic groups from southwestern Angola stand out for exhibiting elevated levels of an unique, regionally-specific and highly divergent Pre-Bantu ancestry. These findings highlight the importance of the Namib for understanding the deep genetic structure of Africa.

## Introduction

It is generally accepted that the pre-colonial genetic diversity of southern Africa resulted from the sequential layering of at least three different sets of peoples with contrasting genetic, linguistic and livelihood profiles: i) an early occupation by the ancestors of foragers now speaking languages of the Kx’a and Tuu families (1–3); ii) a more recent dispersal (~2 kya) of Late Stone Age pastoralists from East Africa, speaking languages of the Khoe-Kwadi family (4, 5); and iii) a subsequent arrival (~1.2 kya) of Iron Age groups tracing their origins to West-Central Africa, who speak Bantu languages and rely to various degrees on agricultural and pastoral lifeways (6–8).

Previous studies have shown that autochthonous southern African foragers are among the most deeply divergent human populations and can be broadly subdivided into three major genetic subgroups associated with the northern, central and southern Kalahari (2, 9–12). Linguistically, the northern Kalahari is home to speakers of the Ju subgroup of Kx’a, while the central and southern Kalahari are mainly linked to the Taa and !Ui branches of Tuu, respectively (SI Appendix, Fig. S1A-B) (13). Beyond the Kalahari Basin, ancestry related to southern African foragers has also been detected in contemporary populations from Zambia and eastern Africa (2, 14), as well as in ancient DNA recovered from individuals who lived in Malawi between 8.1-2.5 kya (15, 16). These findings indicate that foraging populations carrying a genetic ancestry related to modern Kx’a and Tuu speakers were more widespread in the past, before being absorbed or extinguished by incoming food-producers.

The expansion of Bantu-speaking peoples, in particular, is known to have impacted areas showing traces of previous forager occupation and might have played a major role in narrowing the distribution of foraging populations (1, 6, 8) (SI Appendix, Fig. S1D). Nevertheless, in spite of well-known cases of Bantu-forager admixture, most Bantu groups retain their cultural identity and are genetically closer to other Bantu speakers than they are to neighboring non-Bantu populations (8, 17).

In contrast with the Bantu expansion, available data from groups speaking languages from the Khoe-branch of Khoe-Kwadi suggest that the pastoral dispersal from East Africa associated with this language family was strongly shaped by complications arising from areal contact, admixture and diffusion (2, 3, 10, 11, 18). In spite of sharing various amounts of a genetic ancestry related to eastern Africa (3, 15, 19), Khoe groups include both pastoralist and foraging communities who have significant genetic contributions from autochthonous southern African and Bantu-speaking populations (2, 11). Due to this fragmented distribution of genetic and cultural make-ups, the population landscape of southern Africa can only be fully understood by adopting a bottom-up approach that takes into account the complexity of the multiple contact scenarios involving Khoe-Kwadi speakers in different regions. Until now, the best studied areas are the Central Kalahari – home of Kalahari Khoe speakers – and, albeit to a lesser extent, the southern and western Kalahari Basin fringe historically occupied by Khoekhoe speakers (SI Appendix, Fig. S1C). In contrast, little is known about southwestern Angola. Here, the isolated Kwadi branch was spoken until the mid-20th century by a group known as the Kwepe who dwelt in the Angolan Namib desert, close to the Kuroka River mouth (20).

Although both the Kwadi language and speech community were thought to be extinct, we were able to locate a group self-identifying as Kwepe who still live along the Kuroka River, in areas close to their originally reported location (21–23) (SI Appendix, Fig. S2). Despite the Kwepe’s recent shift to the Bantu language Kuvale (24), we identified the descendants of the Kwadi speakers recorded in 1965 by the linguist Ernst Westphal (25) and found two women who still remembered a considerable amount of the now extinct language’s lexicon and grammar. The present-day Kwepe are small stock herders who are surrounded by an array of dispossessed groups also inhabiting the Kuroka River Basin, known as Kwisi, Twa and Tjimba (SI Appendix, Fig. S2). These three communities have been considered to descend from a distinct layer of pre-Bantu foragers, who occupied southern Africa along with the ancestors of Kx’a and Tuu speakers, and whose original language and culture have been lost (26, 27). Regardless of their origins, the Kwepe and the other peoples from the Kuroka River Basin form a cluster of marginalized populations, sharing the languages and cultural habits of the Bantu-speaking Kuvale and Himba, who constitute part of the Herero pastoral tradition of southwest Africa and represent the socially dominant force in the Angolan Namib (26, 28). Due to their strong cultural and socio-economic dependence on their dominant pastoral neighbors, the Kwepe, Kwisi, Twa and Tjimba are best described as peripatetic peoples (29).

Motivated by the highly diverse population landscape of the Angolan Namib, we hypothesize that extant populations from this area may preserve part of the ancestry of the Kwadi, as well as genetic traces of vanished forager populations inhabiting southwestern Africa prior to the Bantu expansion. Here, we generated genome-wide data from 208 individuals belonging to nine ethnic groups from the Angolan Namib and surrounding areas, subsisting on foraging, peripatetic, pastoral and agropastoral lifeways (SI Appendix, Text 1 and Table S1). To contextualize our findings within the wider area of southern Africa and beyond, we further combined the data from Angola with other Africans previously genotyped on the same array (Fig. 1A; SI Appendix, Figs. S2-3 and Table S2). Our results show that the descendants of the formerly Kwadi-speaking Kwepe and the other peripatetic groups from the Angolan Namib preserve a unique pre-Bantu genetic ancestry, highlighting the importance of southwestern Angola as a key area for understanding the peopling of southern Africa

**Figure 1.**
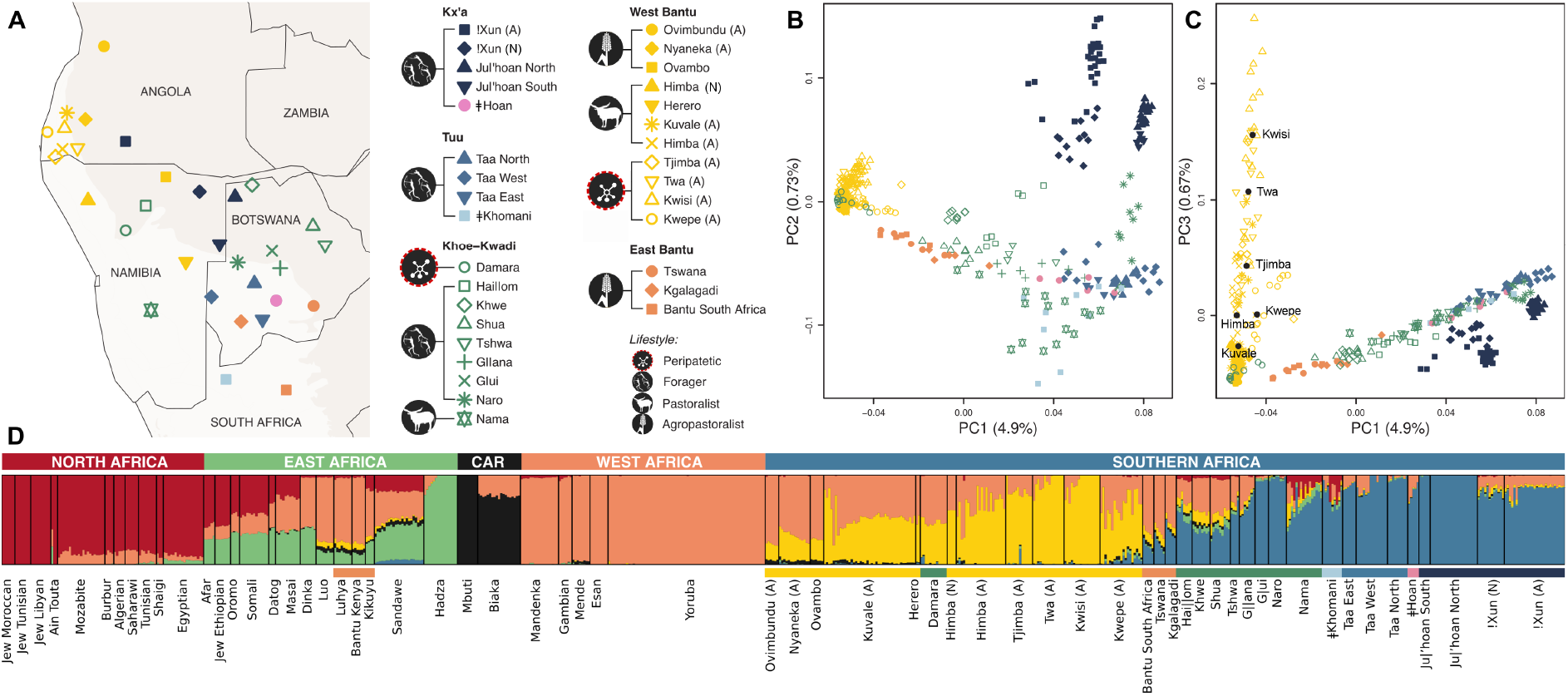
Population structure in southern Africa. A) Map showing approximate sample locations. The lightest background color shows the desert and xeric shrublands biome (Olsen et al., 2001). All samples from Angola are newly generated and marked with “(A)” in the legend, while samples from the same ethnic group collected in Namibia are marked with “(N)”. B-C) Principal Component Analysis of the populations shown in panel A. Plots of PC1 vs. PC2 (B) and PC1 vs. PC3 (C). The centroids for each group from the Angolan Namib are shown by black dots. D) ADMIXTURE analysis for k = 6 (k with the lowest cross-validation error) of an extended dataset of 63 African populations (SI Appendix, Fig. S3, and Tables S1-2). The horizontal color bar below the ADMIXTURE barplot indicates the language group. CAR=Central African Rainforest.

## Results

In a principal component analysis (PCA) (30), the Angolan !Xun foragers are most similar to southern African groups speaking languages from the Ju branch of Kx’a, while all other sampled Angolan groups are closer to West Africans and to Bantu-speaking populations (Fig. 1B; SI Appendix, Fig. S3). Among those groups, the populations from the Angolan Namib display a notable substructure, forming a gradient of genetic differentiation best captured by PC3, which stretches from the Kuvale and Himba cattle herders to the peripatetic Kwisi and Twa (Fig. 1C; SI Appendix, Fig. S4). Additionally, the differentiation displayed in PC3 is correlated (r=0.87; P<0.001) with increasing amounts of an ancestry component revealed at k=6 by the unsupervised population clustering approach implemented in ADMIXTURE (31) (yellow in Fig. 1D; SI Appendix, Figs. S5-8). This component overlaps with a previously identified “NW-Savannah” ancestry that was found to be especially common in northwestern Namibia among the Southwest Bantu speaking Himba, Herero and Ovambo (10, 11). It is also predominant in the Khoekhoe-speaking Damara who are genetically very similar to the Herero and Himba (2, 23), and probably adopted their language from Nama pastoralists with whom they had historically established a peripatetic-like association (SI Appendix, Text 1) (29). The varying proportions of this ancestry in southwestern Angola and northwestern Namibia seem to be broadly associated with subsistence strategy and socio-economic status, being on average lowest in the Nyaneka, Ovambo and Ovimbundu agropastoralists (18-27%) and highest among the peripatetic Tjimba, Twa, and Kwisi (79-93%). An analysis of Identity-By-Descent (IBD) sharing (32) among these populations further indicates that between-group sharing of IBD segments is highest among the peripatetic communities (SI Appendix, Fig. S9).

As the southwestern Angolan pastoral scene is dominated by a highly hierarchized, caste-like matriclanic organization (SI Appendix, Text 1) (23), it is conceivable that the observed genetic structure was caused by drift and inbreeding associated with the marginalization of peripatetic communities. Alternatively, it is also possible that it reflects different levels of admixture with an unsampled or no longer existing population.

To investigate the role of genetic drift, we assessed the levels of genetic diversity in several populations from southwestern Angola and northwestern Namibia by computing for each group the total length of IBD (cM) segments shared between individuals (32) and the total length (Mb) of runs of homozygosity (RoHs) per individual (33). We found that in southwestern Angola, the lowest and highest IBD and RoH lengths are found among agropastoralists and peripatetics, respectively (Fig. 2A; SI Appendix, Fig. S10A). Moreover, the peripatetic groups display much higher mean lengths of shared IBD segments above 10 cM than other southern African populations (Fig. 2B). Estimates of effective population size (Ne) across time leveraging information from shared IBD segments (34) further reveal strong bottlenecks in peripatetic groups starting ~20 generations ago (SI Appendix, Fig. S11). Together, these results suggest that drift played a major role in the genetic differentiation of the Angolan communities with a lower socio-economic status.

**Figure 2.**
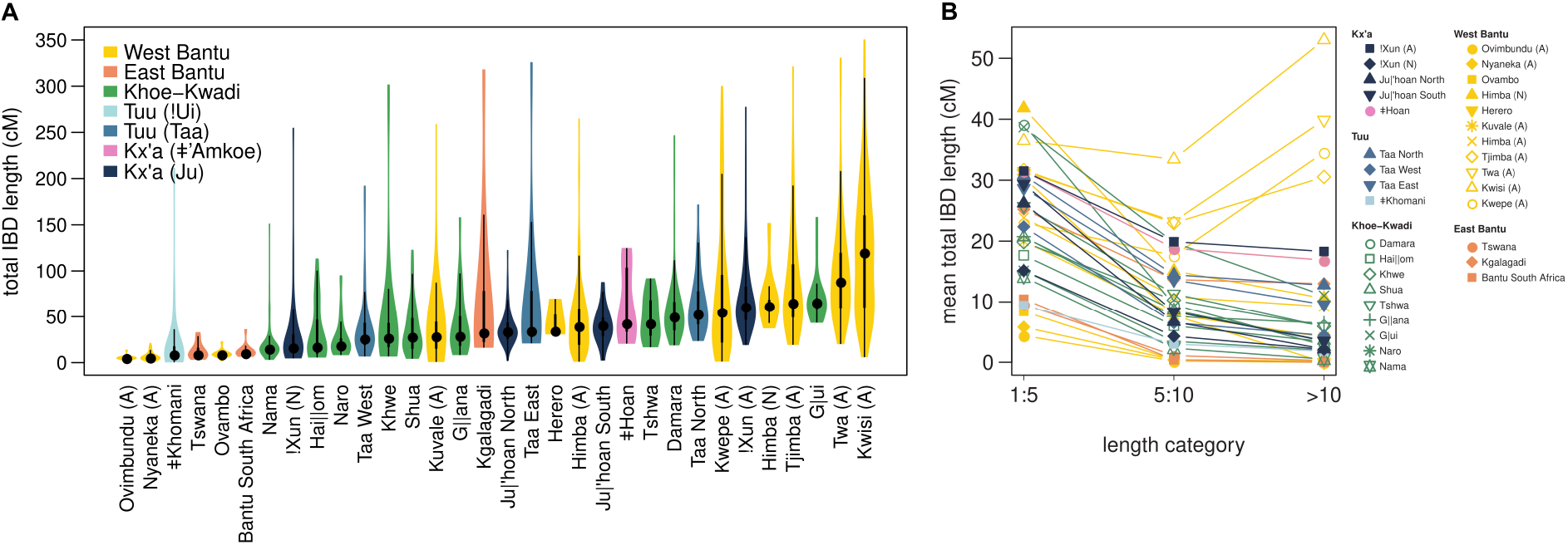
IBD sharing in southern Africa. A) Violin plots showing the distribution of total IBD length (cM) shared withing groups. The thick and thin black lines represent the interquartile range and the upper and lower adjacent values, respectively; the central dot shows the median. B) Mean total IBD length for different length categories.

We next used CHROMOPAINTER under settings that minimize the effects of genetic drift by not allowing pastoralists and peripatetics from southwestern Angola and northwestern Namibia to copy haplotypes from each other (Fig. 3; SI Appendix, Fig. S12) (35, 36). Under these conditions, the Kwepe, Twa and Kwisi display very similar copying profiles that are clearly differentiated from the profiles of their pastoralist (Fig. 3C-D) and agropastoralist (SI Appendix, Fig. S12C-D) neighbors. The profile of the Tjimba is close but not identical to the other peripatetics, while the Damara fully align with the Bantu-speaking pastoralists (Fig. 3C-D; SI Appendix, Fig. S11C-D). Comparisons of profile differences between each group and representative pastoralist (Kuvale) and agropastoralist (Ovimbundu) populations further show that peripatetic groups have decreased amounts of Bantu and West African-related ancestries, while copying elevated numbers of haplotypes from groups carrying southern African forager ancestry and, to a lesser extent, the Mbuti rain forest hunter-gatherers and several East African groups (Fig. 3A-B; SI Appendix, Fig. S12A-B). This finding highlights the impact of differential admixture with pre-Bantu populations and suggests that drift and inbreeding were not the only factors influencing their genetic differentiation.

**Figure 3.**
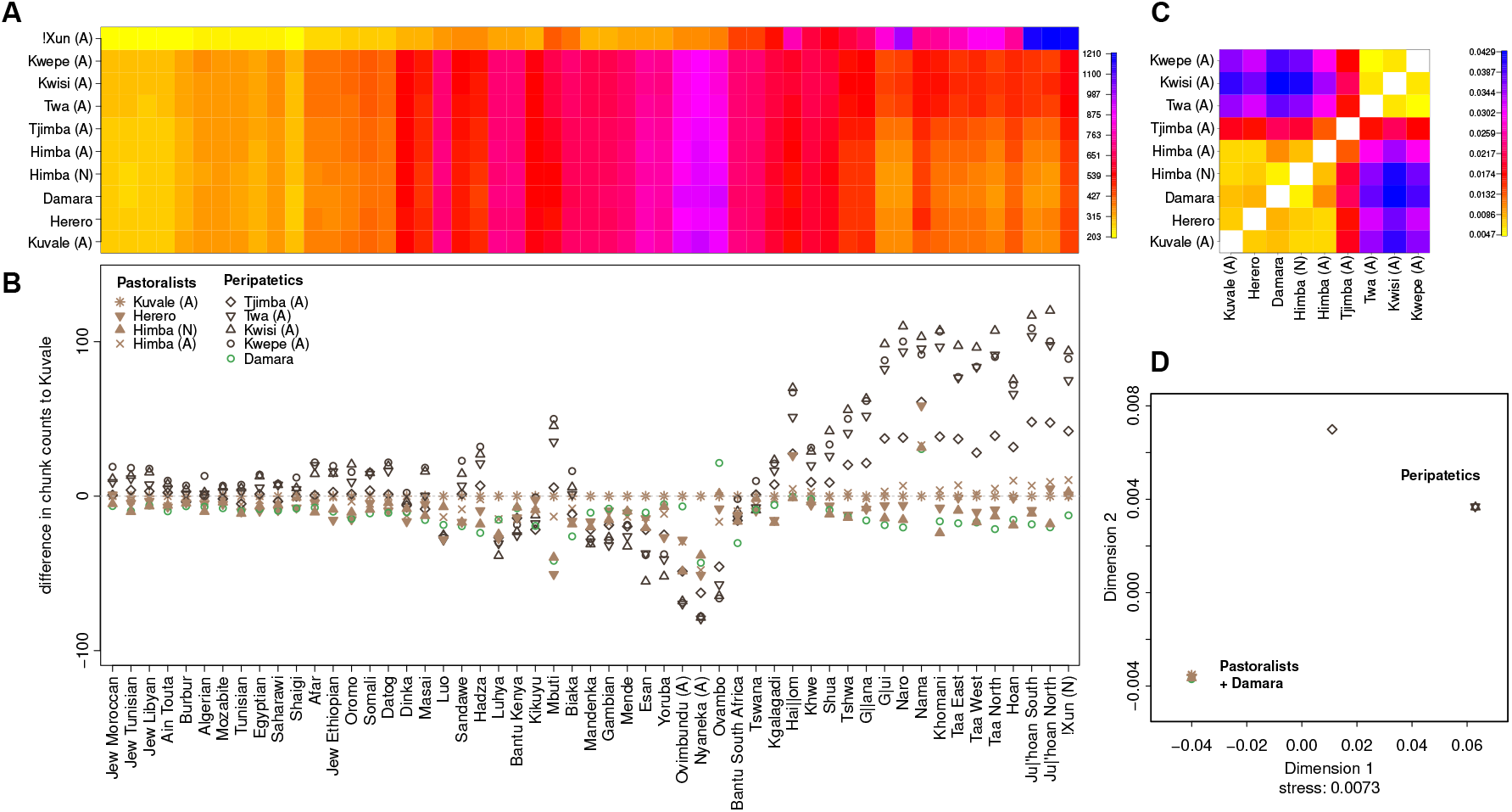
Chromopainter profiles for southwestern Angolan and northwestern Namibian groups. Bantu-speaking agropastoralists (Ovimbundu, Nyaneka, and Ovambo) are included as donors (columns), together with more distantly related African groups. A) Chromopainter coancestry matrix. The color gradient indicates the average number of DNA chunks a group copies from the donor populations. B) Differences between the number of DNA chunks copied by each group and the Kuvale – used here as a baseline. C) Average distance (TVDxy) between copying profiles. D) Multidimensional scaling calculated on the TVDxy distances.

To further assess the role of pre-Bantu admixture while including ancient individuals as potential sources (15), we used qpAdm (37). By testing the fit of different mixture models, we confirm that the peripatetic peoples (Kwepe, Kwisi, Tjimba and Twa) diverge from their neighbors by displaying higher amounts (10-14%) of southern African forager ancestry (here represented by an ancient South Africa 2000BP genome), and a detectable contribution (4-5%) from East Africa, best matched by an ancient genome retrieved from a pastoral context in Tanzania (Luxmanda 3100BP) (Fig. 4; SI Appendix, Text 2, Fig. S13, and Tables S3-4).

**Figure 4.**
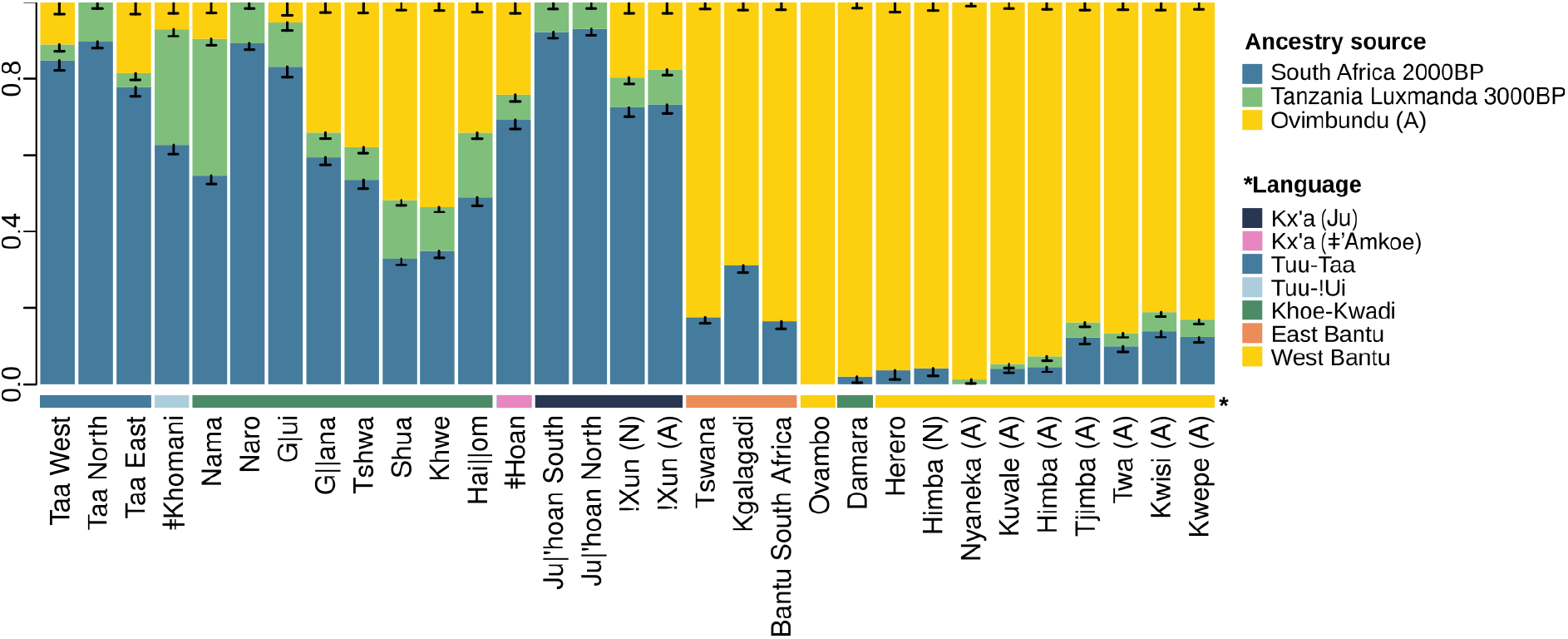
Ancestry proportions estimated with qpAdm in southern African populations. Standard errors (black bars) were calculated with a weighted block jackknife.

Our inference of the relative order of mixing of the Bantu-, autochthonous southern African- and East African-related ancestries in southwestern Angola and northwestern Namibian groups, based on ancestry covariances (38), indicates that the first admixture event involved southern African- and East African-related ancestries (SI Appendix, Table S5), in accordance with the archaeological data suggesting that pastoralism was introduced from eastern into southern Africa prior to the Bantu expansion (39). While we could not obtain reliable estimates for this admixture event (SI Appendix, Text 3), we detected signals of admixture between Bantu and the previously admixed pre-Bantu ancestries dating to ~600-1100 years ago by using Wavelets, GLOBETTROTER, and ALDER (SI Appendix, Fig. S14, and Tables S5-7).

Like the Angolan Namib, other areas where languages of the Khoe-Kwadi family are spoken today were strongly shaped by contact and display highly variable amounts of southern African-, East African- and Bantu-related ancestries (Fig. 4). To reconstruct the contact histories of the Khoe-Kwadi following their arrival to southern Africa, we carried out local ancestry decomposition (40) and analyzed the population relationships within each of the three major settlement layers of southern Africa (SI Appendix, Fig. S15-S18).

Based on PCA projections, we found that the East African ancestry identified in the genomes of Khoe-Kwadi speakers and other southern Africans is related with pastoralist groups clustering around the ancient Tanzania Luxmanda individual (3100BP) (SI Appendix, Fig. S16). Some Nama and ǂKhomani individuals are additionally related to East African groups with high amounts of Eurasian ancestry, likely due to admixture with Europeans during colonial times.

Ancestry-specific PCA (SI Appendix, Figs. S17-18) and clustering analysis based on average pairwise differences (SI Appendix, Fig. S19) further show that southern African- and Bantu-related ancestries of Khoe-Kwadi groups are highly heterogeneous and mirror the genetic composition of their neighbors. This pattern becomes especially clear when the local ancestry information is combined with IBD inferences to obtain southern African- and Bantu-specific IBD sharing (Fig. 5; SI Appendix, Fig. S20). For example, the Khwe from the northern Kalahari Basin fringe have southern African- and Bantu-related ancestries that are similar to those of !Xun foragers and Southwest Bantu agropastoral groups living in the same area (Fig. 5; SI Appendix, Fig. S20). To their south, Khoe-speakers from the Central Kalahari share southern African-related ancestry with the neighboring Taa and ǂHoan, and Bantu-related ancestry with local East Bantu speakers (Fig. 5; SI Appendix, Fig. S20). The southern African- and Bantu-related ancestries of the Khoekhoe-speaking Nama reflect their migration history along the Atlantic coast of southern Africa. While their southern African-related ancestry resembles that of the ǂKhomani, who inhabit the southernmost areas of the Kalahari, their Bantu ancestry is similar to that of Southwest Bantu-speaking groups from northwestern Namibia (Fig. 5; SI Appendix, Fig. S20). As the Nama are known to be a branch of Khoekhoe-speaking groups who migrated northwards from South Africa (41), it is likely that they first acquired their southern African-related ancestry in the South, and admixed with Bantu populations only later after reaching Namibia.

**Figure 5.**
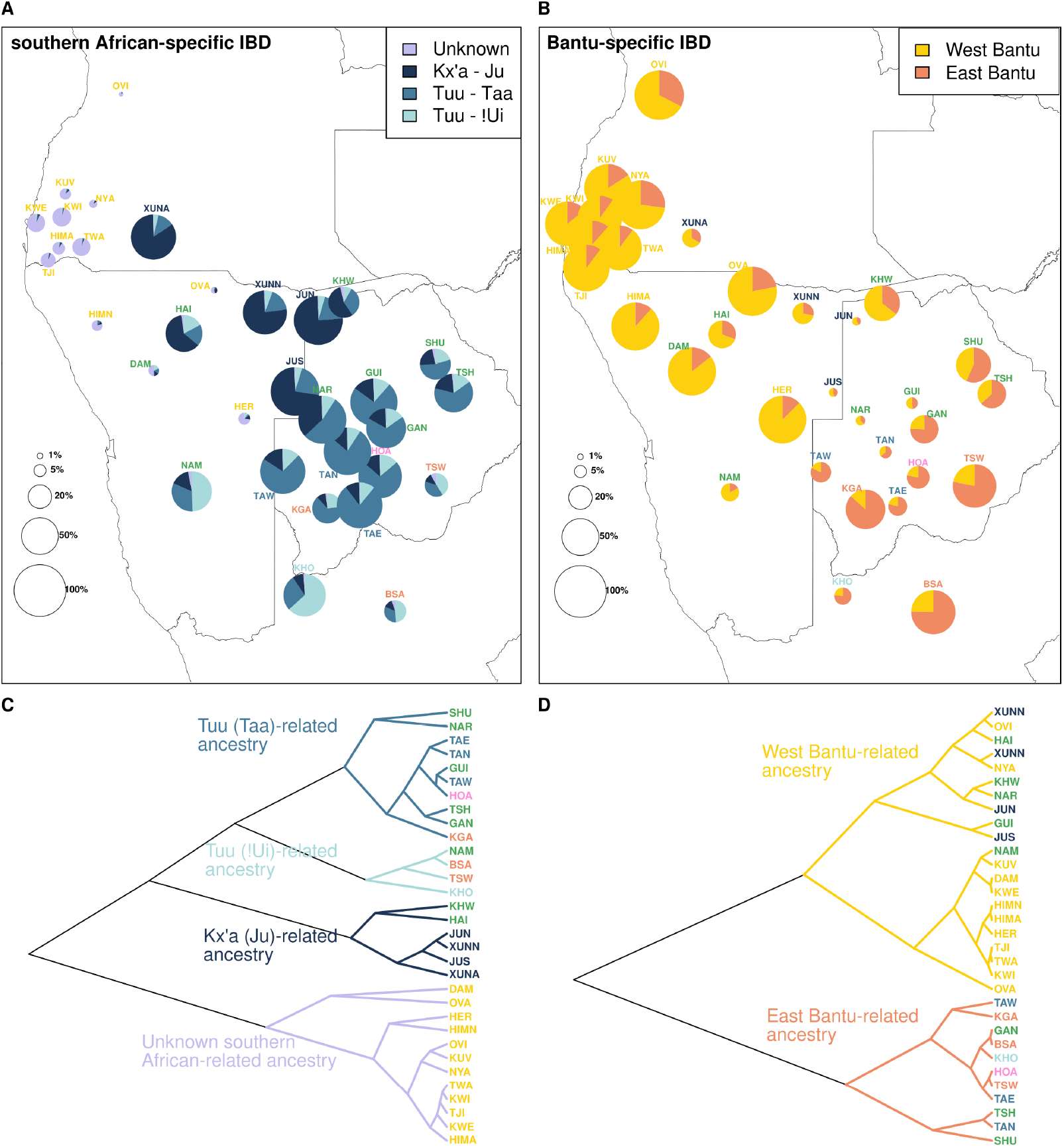
Summary of the ancestry-specific IBD sharing. The pie-charts represent the proportion of autochthonous southern African-related (i.e., neither Bantu-nor East African –related) (A) and Bantu-related (B) IBD shared on average between each target group and the pools of populations identified in the figure legend. The category “Unknown” represents southern African-related IBD shared among the peripatetic groups from the Angolan Namib. The percentage of southern African (A) and Bantu (B) ancestry within each target group is represented by the total area of the pie-chart. (C) and (D) show an UPGMA clustering of all target groups according to their shared southern African- and Bantu-related IBD, respectively. Detailed results for every population pair are shown in Fig. S20. Population abbreviations according to Tables S1-2.

In the Angolan Namib, the formerly Kwadi-speaking Kwepe and the other peripatetic groups all share Bantu-related ancestry with the southwestern Bantu pastoralists that surround them (Fig. 5; SI Appendix, Fig. S19-20). However, their southern African-related ancestry does not match any of the major ancestry components that have previously been described in southern Africa (Fig. 5; SI Appendix, Fig. S19-S20): despite sharing large amounts of southern African-related IBD segments among themselves, the peripatetics stand out for their lack of IBD sharing with present-day southern African forager groups (Fig. 5; SI Appendix, Fig. S20), suggesting that their southern African-related ancestry resulted from admixture with a deeply-divergent unsampled group. The same ancestry is also found in other groups from southwestern Africa, including the Damara from Namibia, but the detected frequencies are much lower than in the Angolan peripatetics (Fig. 5; SI Appendix, Fig. S20). The uniqueness of this new genetic component (henceforth called KS-Namib) is also supported by a PCA analysis undertaken with the EMU approach (42). This approach, which allows for the detection of population structure even with high levels of missing data, shows that KS-Namib can be readily separated from all known major African ancestries, including the southern African-related component identified in ancient (8,100-2,500 BP) hunter-gatherers from Malawi (Fig. 6 C, D).

**Figure 6.**
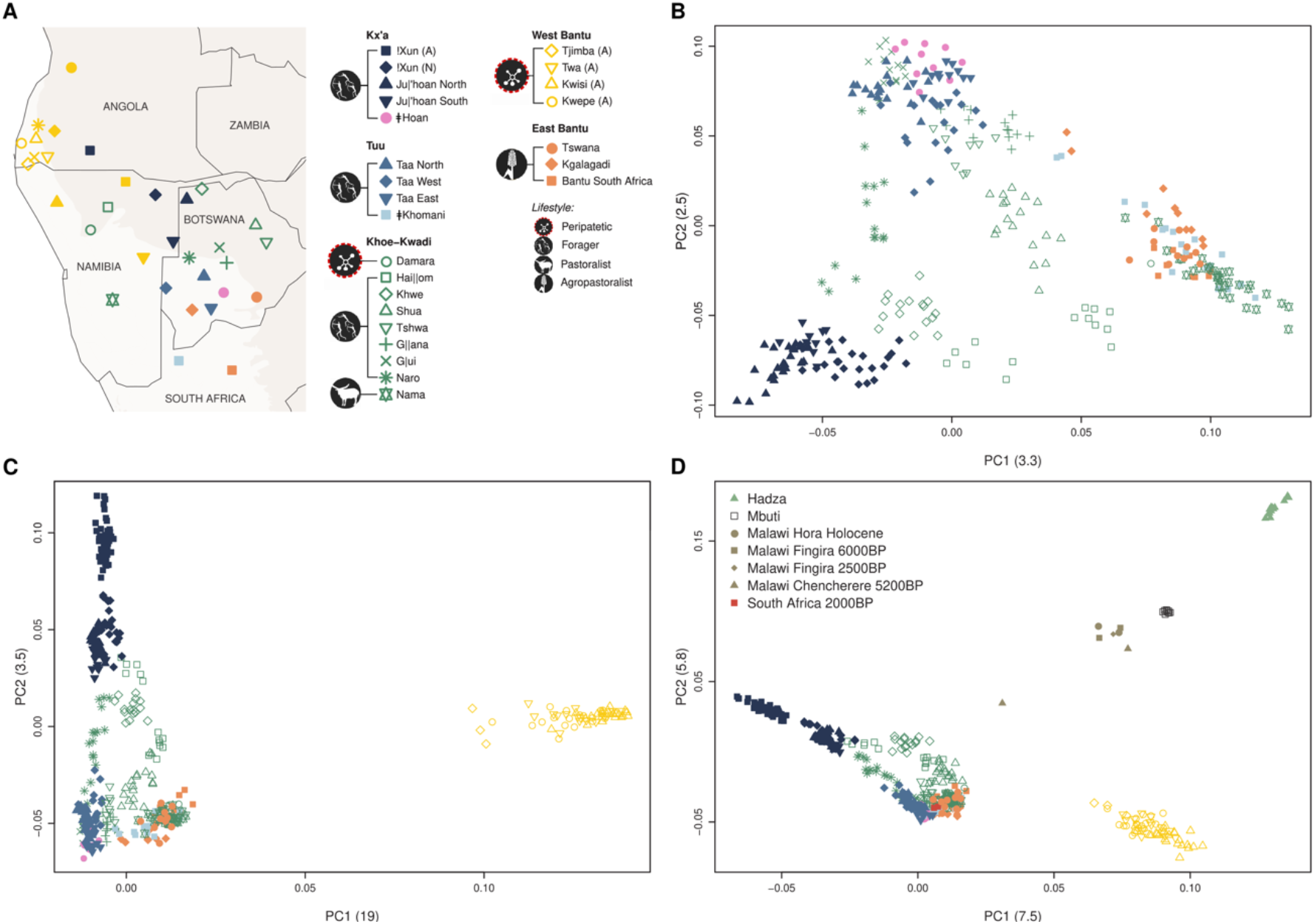
PCA of southern African-related ancestry. A) Map showing approximate sample locations. The lightest background color shows the desert and xeric shrublands biome (Olsen et al., 2001). B-D) PCA build with previously published southern African individuals that display <10% of missing data after masking the non-southern African ancestries (B), additionally including Angolan individuals (C), and other relevant modern or ancient groups (D).

Taken together, these results suggest that the Angolan Namib Desert and its surrounding areas preserve the legacy of an extinct, deeply divergent human group with no close matches in extant populations from within and outside southern Africa.

## Discussion

The Angolan Namib Desert provides an invaluable framework to examine the history and consequences of contact and admixture between different migratory waves into the wider region of southern Africa. In spite of being culturally dominated by southwestern Bantu-speaking cattle herders, the area is remarkable for the presence of several impoverished groups with a peripatetic way of life, who have attracted a considerable degree of ethnographic interest (20, 26, 41, 43, 44). Our results highlight the heterogeneity of the population landscape of the Angolan Namib by showing that, despite their high amounts of Bantu ancestry (~80%), all sampled peripatetic groups display elevated levels of an eastern African ancestry and a previously undocumented southern African-related component (KS-Namib) (Figs. 4-6). The co-occurrence of the two pre-Bantu ancestries among all peripatetic groups hints at a complex contact and admixture history. Since the eastern African component has also been detected in Khoe-speaking groups of the Kalahari Basin (Fig. 4), it is likely that it was introduced to southwestern Angola by the ancestors of the present day Kwepe as part of the Kwadi branch of the Khoe-Kwadi pastoral dispersal. By contrast, KS-Namib has a more restricted distribution, being especially common in the Angolan Namib and appearing in residual amounts in other groups of southwestern Africa, including the Damara from Namibia who are historically linked to a peripatetic way of life (Fig. 5). This distribution suggests that KS-Namib was more likely associated with a resident foraging population of southwest Africa than with migrants from elsewhere.

While at present no hunter-gatherers resembling Kx’a- and Tuu-speaking groups exist in the Angolan Namib, an early account by 16th century traveler Duarte Pacheco-Pereira states that the areas around the Kuroka River mouth were then inhabited by nomadic groups who lived on fishing and built houses from whale ribs which they covered with seaweed (45). This description is reminiscent of coastal foragers often referred to as “Strandlopers” who once lived near the seaside in Namibia and South Africa, but went extinct during the 19th century (46). Although historical records note them to be Khoekhoe-speaking, their culture differed from the herding groups further inland, and their origins may ultimately trace to resident hunter-gatherer groups, as suggested by the long history of maritime foraging in southern Africa (47). An ancient, pre-pastoral origin of southern African marine foragers was recently supported by genome-wide data from a 2,241-1,965 kya old skeleton excavated at St. Helena Bay on the coast of South Africa (15, 48). However, the genetic profile of this individual is close to contemporary Tuu-speaking hunter-gatherers from inland areas of southern Africa and not to the KS-Namib component (Fig. 6). Hence, it is likely that only studies of ancient DNA from the southwestern coast of Angola will be able clarify the ancestral relationships between KS-Namib and extinct forager populations. While the archaeological record for the Angolan Namib is sparse, it is possible that prehistoric human remains associated with this ancestry may be recovered around shell midden deposits and vestiges of coastal settlements that have been reported in areas close to the Kuroka River mouth (49, 50).

An ancient occupation of the Namib coast is also supported by oral stories describing an encounter between the Kwadi-speaking Kwepe and resident peoples who had no fire and ate raw fish on the beach (26, 49, 51). Some anthropologists have equated these resident fishermen with the ancestors of the Kwisi and Twa, due to their present socio-economic marginalization and historically-documented association with hunting and gathering, which contrasts with the Kwepe’s higher reliance on small-scale pastoralism (26, 49, 51).

However, while our results show that present-day Twa and Kwisi can be genetically separated from the Kwepe (SI Appendix, Fig. S4 and S6), the three groups are virtually indistinguishable once the effects of genetic drift are attenuated (Fig. 3, SI Appendix, Fig. S12). This pattern suggests that all extant peripatetic groups are equally related to different ancestries, thus challenging any attempts to establish continuity between specific modern populations and ancient foragers, beyond ethnographic considerations. The microcosmos of the Angolan Namib can therefore be considered a highly stratified polyethnic system (52) where groups with different genetic and ethnolinguistic backgrounds admixed, but maintained sharp divisions based on their socio-economic status.

This contact profile of southwestern Angola is remarkably similar to other areas of southern Africa where Khoe-Kwadi-speaking migrants encountered resident populations with different linguistic and genetic legacies. A defining characteristic of all these areas is an admixture history which starts with the fusion between an eastern African ancestry and different resident southern African forager components, later followed by various degrees of admixture with Bantu speakers from the western and eastern streams of the Bantu migrations (18) Considering the available genetic and archeological data (39, 53), we hypothesize that after an initial break-up of the proto-population in the northwestern Kalahari, the Khoe-Kwadi diverged into different groups migrating into specific contact areas (Figs. 4-5; SI Appendix, Fig. S1 and Fig. S20): while Khoekhoe speakers moved south, encountering !Ui-speaking groups, Kalahari Khoe-speakers took an eastward route where they encountered Ju speakers in the northern, and Taa and ǂHoan speakers in the central Kalahari. Finally, the Kwadi branch migrated towards southwestern Angola, arriving in areas inhabited by a now extinct foraging group associated with the KS-Namib component. More recently, most Khoe-Kwadi-speaking peoples were further impacted by East and West Bantu-speaking populations, adding to their diverse genetic make-ups (Fig. 5).

Together, our results show that contact areas associated with the confluence of different migratory waves can harbor the ancestry of vanished groups predating the arrival of food-production in Africa. While the full diversity and geographical extension of these early foragers may ultimately be revealed by ancient DNA, detailed studies of highly admixed small-scale communities can still provide unique opportunities to probe the deep genetic structure of the continent.

## Materials and Methods

### Sample information

This study includes a total of 208 samples from nine ethnic groups of southwestern Angola. The sample size, linguistic affiliation, subsistence pattern, and sampling locations for each group are summarized in Table S1. Samples from the Nyaneka and Ovimbundu were collected as described in (54), and samples from the remaining populations (!Xun, Kwepe, Kwisi, Twa, Tjimba, Himba, Kuvale) collected as described in (21). Written informed consent was obtained from all participants. This study was developed in the framework of a collaboration between the Portuguese-Angolan TwinLab established between CIBIO/InBIO and ISCED/Huíla Angola, has the ethical clearance of ISCED and the CIBIO/InBIO-University of Porto boards, and the support and permission of the Provincial Governments of the Namibe and Kunene.

### Genotyping and quality control

All sampled individuals were genotyped on the Affymetrix Axiom Genome-Wide Human Origins Array (55). The newly generated data were analyzed together with data from 54 African populations (550 individuals) previously genotyped on the same array (Table S2) (2, 55, 56), after filtering out SNPs with a missing rate higher than 10%, SNPs with deviations from Hardy-Weinberg equilibrium (i.e., p-value < 0.001 in more than two populations), and SNPs from non-autosomal markers. These filters yielded a final set of 607,761 SNPs. No individuals had missing rates above 10%. We excluded from the analyses 37 individuals from Angola and 25 individuals from elsewhere due to cryptic relatedness. Specifically, we removed one individual from each pair that exhibited a proportion of identity by descent (IBD) higher than 0.2, computed in PLINK v1.9 as P(IBD=2) + 0.5*P(IBD=1) (33). We additionally generated a dataset pruned for linkage disequilibrium with PLINK v1.9 (33), removing SNPs with r2 > 0.4 in 200 kb windows, shifted at 25 SNP intervals. The pruned dataset includes a total of 350,719 SNPs. These datasets were additionally merged with ancient samples from (15, 57, 58).

### Phasing

All present-day samples analyzed were jointly phased with Beagle 4.1 (59) using the HapMap genetic map available from the Beagle website (https://faculty.washington.edu/browning/beagle/beagle.html).

### Population structure analyses

#### a. Genotype-based

PCA was computed using the smartpca software from EIGENSOFT 6.0.1 (30), without exclusion of outliers (‘numoutlieriter: 0’). ADMIXTURE v1.3 (31) was used to estimate ancestry proportions originating from k ancestral populations, with k ranging from 2 to 16, and applying a cross-validation procedure, for a total of 15 independent runs. Additionally, we used DyStruct v.1.1.0 (60) to identify shared ancestry with relevant ancient individuals from East and Southern Africa, taking into account the individual’s archaeological age and assuming a generation time of 29 years (61). The DyStruct analysis included only a subset of the present-day groups used in the ADMIXTURE analysis and the Angolan groups were additionally downsampled to lower the impact of large sample sizes in clustering analysis. We performed 6 independent runs, using 2 to 6 ancestral populations. The ADMIXTURE and DyStruct results were plotted with pong (62).

These analyses were carried out using the LD-pruned dataset.

#### b. Haplotype-based

We used refinedIBD v.17Jan20.102 (32) to identify IBD blocks shared between individuals and homozygous-by-descent (HBD) blocks shared within each individual. Throughout the text we refer to the combined IBD and HBD blocks simply as IBD. Blocks within a 0.6 centiMorgan (cM) gap were merged using the software merge-ibd-segments v.17Jan20.102 (https://faculty.washington.edu/browning/refined-ibd), allowing one inconsistent genotype between the gap and block regions. The IBD blocks were then partitioned into three length categories (1–5 cM, 5–10 cM, and over 10 cM) to investigate IBD sharing across different time periods (63, 64). The average total length shared within and between populations was summarized for each category using networks build with the R package ggraph v2.0.5, and the distribution of total lengths shared within populations was inspected with the R package vioplot v.0.3.7.

We estimated changes in the effective population size (Ne) though time based on IBD blocks of at least 2 cM shared within populations, using the software IBDNe v.23Apr20.ae9 (34). The Ne is shown for generations 4 to 50, which corresponds to the time period for which IBD segments are informative when using SNP array data (34).

ROH were identified in PLINK v1.9 (33) using the LD-pruned dataset and default parameters: sliding-windows of 50 SNPs, a ROH minimum length of 1 Mb, five missing genotypes and one heterozygote allowed per window, and a scanning window hit rate of 0.05 required for a SNP to be eligible for the ROH.

We used CHROMOPAINTER v2 (35) to infer a “painting” or copy profile for individuals from southwestern Angola and northwestern Namibia, based on two different sets of donor and recipient populations. These were chosen to test if all groups share a common ancestor before any recent isolation. In the first set, pastoralists and peripatetics from southwestern Angola and northwestern Namibia (recipients) were only allowed to copy haplotypes from Bantu-speaking agropastoralists (Nyaneka, Ovimbundu, Ovambo) or more distantly related African groups, therefore minimizing differences in copy profiles caused by recent genetic drift (Fig. 3). In the second set, agropastoralists were included as recipients and excluded from the donors, hence all individuals from southwestern Angola and northwestern Namibia could only copy haplotypes from distantly-related African groups (SI Appendix, Fig. S12). To account for the impact of uneven donor sample sizes in the resulting copy vectors, each analysis was run three times with a random sample of five individuals per donor population. We used all of the available individuals for populations with less than five individuals.

We initially estimated the mutation emission and switch rate parameters using 10 iterations of the Expectation-Maximisation (EM) algorithm and a subset of chromosomes (1, 5, 10, 15 and 22). The inferred parameters were averaged by chromosome (taking into account their number of SNPs), and then by individuals. We fixed these parameters and performed a new CHROMOPAINTER run for all chromosomes.

The copy profiles generated for each recipient individual under each analysis were displayed in coancestry matrices that represent the average copy profiles of three runs. The difference between the average copy profiles of any pair of populations x and y, under each set, was quantified using the total variation distance (TVDxy) (36, 65). A non-metric multidimensional scaling (MDS) was carried out in R, using the function isoMDS.

### Ancestry modeling with qpAdm

We used qpAdm v.650 (55) to test one-wave, two-wave, and three-wave admixture models for each southern African population and to estimate their respective admixture proportions. The tests were conducted using the same reference and source populations as in (15) - chosen to capture major strands of ancestry in Sub-Saharan Africa, as well as using a modified set of reference and source populations that is best suited for populations of Southern Africa (SI Appendix, Text 2 and Fig. S13). We applied a “rotating” strategy (15, 66) for each of the two sets of populations, in which a defined number of sources (one, two, or three) was selected iteratively from a source pool, while all the other populations in the set were used as references. We rejected models if their P-values were lower than 0.05, if there were negative admixture proportions, or if the standard errors were larger than the corresponding admixture proportion. As more than one model was often accepted per population (SI Appendix, Table S3 and S4) we display the results according to different criteria (SI Appendix, Text 2 and Fig. S13).

### Local ancestry inference

Local ancestry inference was carried out using RFMix v2 (40) (https://github.com/slowkoni/rfmix) for all southern African individuals. Equal size samples (n=13) of Yorubans, Somalis, and southern African individuals (8 Ju|’hoan North, 5 Taa West) that occupy the rightmost position in PC1 (Fig. 1 B-C) were used as training sources to capture ancestry related to the Bantu expansion, the eastern African pastoral migration and the indigenous southern African hunter-gatherer stratum, respectively. We ran RFMix with three iterations and the option “reanalyze-reference” to account for any admixture in the references, used a minimum of five reference haplotypes per tree node, and assumed 25 generations since the admixture event.

### Admixture timing

The relative order of mixing of different ancestries in admixed populations from southwestern Angola and northwestern Namibia was inferred using the Admixture History Graphs (AHG) approach (67). In an admixed population with two ancestry components (A and B) that later receives a third component (C) via admixture, the ancestry proportions of A and B will covary with C, but the ratio of A and B throughout the population will be independent from C. The AHG approach involves estimating the covariance of the frequencies of A/B with C, A/C with B, and B/C with A, across all individuals in the population, and identifying the configuration that produced the smallest absolute value of the covariance estimate. The individual ancestry proportions used in this test were those inferred by RFMix.

The dating of the admixture events was obtained via the wavelet-transform analysis (38), which uses the width of ancestry blocks identified by RFMix to determine the time since admixture by comparing the results to simulations (38, 67). In this analysis, we assume the order of events as inferred by the AHG approach.

We additionally estimated admixture events with GLOBETROTTER (68). First, to minimize any noise in the admixture inference caused by outlier individuals we performed a fineSTRUCTURE analysis (35), which hierarchically divides individuals into genetically homogeneous groups. By comparing those groups with the ethnic label of each individual we identified a total of 32 outlier individuals (3 Kwepe, 3 Kwisi, 5 Twa, 3 Tjimba, 4 Himba, 8 Kuvale, and 5 !Xun), which we excluded from the admixture analyses. Then, we performed another CHOMOPAINTER run in which both recipient and surrogate individuals were “painted” by the same set of donors as in Fig. 3. Finally, we ran GLOBETROTTER, using 10 painting profiles per individual and the coancestry matrix for the total length of haplotype sharing obtained with CHROMOPAINTER. All donor populations were included as possible surrogates (i.e. sources of admixture). Briefly, GLOBETROTTER dates a maximum of two admixture events based on the decay of linkage disequilibrium (LD) versus genetic distance among the segments copied from pairs of surrogate populations, and infers the sources of admixture as a linear combination of the DNA of the sampled groups. As recommended by the authors (68), we performed this analysis with and without a “NULL” individual to evaluate the consistency of the estimates. We ran 50 bootstrap iterations to infer confidence intervals for the date estimates.

For comparison, we also estimated the time of admixture between Bantu and Southern African ancestries based on the exponential decay of Linkage Disequilibrium (LD) using ALDER v1.03 (69) with default settings and using as references the Yoruba and Ju|’hoan North. All admixture dates are presented assuming a generation time of 29 years (61).

### Ancestry-specific analyses

For the ancestry-specific analyses, all ancestries except the one of interest were masked (treated as missing) in each haploid genome. Additionally, we masked all positions within each haploid genome for which the marginal probability returned by RFMix was smaller than 1, thus excluding any parts of the genome where the ancestry assignment was ambiguous. We first confirmed the validity of the local ancestry inference and masking procedures by computing a PCA on the training sources used in RFMix (Yoruba, Somali, and the least admixed Ju|’hoan North and Taa West individuals) and projecting on it all target southern African haploid genomes after masking (SI Appendix, Fig. S15). The PCA was carried out with smartpca (55) using the lsqproject option. Note that to keep standard file format requirements while allowing for unequal missingness in the two haploid genomes composing an individual, we treat each haploid genome as an individual in the PCA projections, therefore displaying twice the number of diploid individuals. Since all genomes with less than 95% missing data for a given ancestry overlap with the expected source population, we used this cut-off for the ancestry-specific PCA.

The PCA targeting East African-specific ancestry was built with unmasked genomes from present-day East African populations, excluding Bantu-speaking groups and the Luo (who display a Bantu-related profile) and filtering out SNPs with a missing rate higher than 10% (SI Appendix, Fig. S15). Southern African individuals with less than 95% missing data after masking the non-East African ancestry were then projected, together with previously published ancient genomes from East Africa (SI Appendix, Fig. S16). The PCA of the autochthonous southern African ancestry was built with a subset of southern African individuals displaying <25% of missing data after masking the non-southern African ancestries, and the additional removal of SNPs with a missing rate higher than 15%. The remaining southern African individuals having between 25% and 95% missing data after masking were projected (SI Appendix, Fig. S17). Similarly, the PCA of Bantu-specific ancestry was constructed using southern African individuals, as well as Bantu-speaking groups from East Africa, having <25% of missing data after masking the non-Bantu ancestries, with the additional removal of SNPs with a missing rate higher than 15%. The remaining individuals having between 25% and 95% missing data after masking the non-Bantu ancestries were projected (SI Appendix, Fig. S18).

PCA was additionally carried out with an alternative method (EMU-EM-PCA for Ultra-low Coverage Sequencing Data), designed to handle high missingness in genetic datasets (Fig. 6) (42). This analysis was computed on two eigenvectors, using southern African individuals that have <10% of missing data after masking.

We calculated ancestry-specific pairwise differences and constructed heatmaps with built-in dendrograms using R (https://www.R-project.org/). Specifically, we used the function hclust, the hierarchical clustering method complete linkage, and a correlation matrix to compute the distance between both rows and columns of the pairwise distance matrix, represented in R by as.dist(1-cor(x)) (SI Appendix, Fig. S19).

Finally, we combined the information on IBD sharing between southern African individuals with their local ancestry profiles to obtain ancestry-specific IBD segments. For this analysis, we used the raw IBD blocks before the merging step, since that step would lead to loss of information about the haplotype of origin. While an IBD block might be a mosaic of more than one ancestry, the ancestry along two haplotypes that are identical-by-descent should in theory match. Yet, in practice mismatches can be found if the RFMix inference is not perfect. Thus, we excluded from this analysis IBD blocks whose ancestry along both haplotypes is inconsistent for more than 25% of the corresponding IBD length (in cM). For each of the remaining IBD blocks shared between two (haploid) individuals we record the length of the block (in cM) associated with each specific ancestry, and then compute the sum of all lengths per ancestry. The average sum of IBD lengths (cM) from a specific ancestry, as well as the average sum of IBD lengths excluded due to inconsistent ancestry assignments are displayed for each pair of populations using stacked barplots (SI Appendix, Fig. S20). This procedure, made available in https://github.com/sroliveiraa/asIBD, was applied separately for IBD blocks belonging to different length categories (1–5 cM, 5–10 cM, and over 10 cM).

We additionally summarize the results for length category 1–5 cM by computing the average IBD sharing between each group and pools of populations representing the genetic diversity of southern African and Bantu-related ancestries. These comprise West Bantu and East Bantu-related ancestries, as well as southern African ancestries represented by Kx’a – Ju, Tuu – Taa and Tuu -!Ui-speaking populations. We further include in the comparisons an “unknown” component to account for the southern African-related ancestry shared with peripatetic groups from the Angolan Namib (Fig. 5A-B). A UPGMA cladogram representing the Eucledian distance between proportions of IBD sharing between each population and these major groups was built with the function upgma of the R package poppr (Fig. 6C-D).

## Acknowledgments

We thank all sample donors for their participation in this study, the governments of Namibe and Kunene Provinces in Angola for supporting our work, João Guerra, Raimundo Dungulo, and Serafim Nemésio for assistance in the preparation of fieldwork, António Mbeape, José Domingos, and Okongo Toko for assistance with sample collection, Irina Pugach for help running wavelets, and Magda Gayà-Vidal for valuable discussions on a previous version of this paper. The research presented in this paper was carried out as part of the Portuguese-Angolan TwinLab established between BIOPOLIS-CIBIO and ISCED/Huíla, Lubango. Financial support was provided by FEDER funds through the Operational Programme for Competitiveness Factors—COMPETE, by National Funds through FCT—Foundation for Science and Technology under PTDC/BIA-EVF/2907/2012, FCOMP-01-0124-FEDER-028341, PTDC/BIA-GEN/29273/2017, by the project NORTE-01-0246-FEDER-000063, supported by Norte Portugal Regional Operational Programme (NORTE2020), under the PORTUGAL 2020 Partnership Agreement, through the European Regional Development Fund (ERDF), and by the Max Planck Society. S.O. was supported by the FCT grant SFRH/BD/85776/2012, A.M.F by the FCT contract CEECIND/02765/2017, and B.A. by the FCT grant 2021.07988.BD. This paper is dedicated to the memory of our late colleague Prof. Samuel Aço.

