## Supplementary Material for "Genome wide variation in the Angolan Namib desert reveals unique Pre-Bantu ancestry"

Sandra Oliveira

Anne-Maria Fehn

Beatriz Amorim

Mark Stoneking

Jorge Rocha

**This PDF file includes:**

Supporting texts

1. Presentation of the studied populations
2. *qpAdm* analyses
3. Admixture dating analyses

SI References

Figures S1 to S20

**Supporting Information Text**

**SI Text 1. Presentation of the studied populations**

The population landscape of southwestern Angola is characterized by a diverse set of ethnolinguistic groups who represent the major population layers of southern Africa.

The !Xun or Sekele (Fig. S2), sampled in the province of Kunene, form the northernmost extension of peoples speaking languages from the Ju branch of Kx'a (Fig. S1A) and are linguistically and culturally related to !Xun and Ju|'hoan groups living in northern Namibia and Botswana (1, 2). They mostly subsist on hunting small game and collecting wild food, but also perform temporary work for their dominant Bantu neighbors who speak the Ambo dialect Kwanyama.

The Kuvale and Himba are two Bantu-speaking groups from the broad Herero ethnolinguistic division (3, 4), which forms part of the "Cimbebasia" subgroup of southwest Bantu (5). The Kuvale and Himba inhabit the northern and southern semi-arid areas of the Angolan Namib Desert (Fig. S2), respectively, and maintain a strong pastoral tradition which is shared by other Himba and Herero communities from Namibia. They are economically and socially dominant over the ethnic minorities of the Namib desert, as attested by the dissemination of their languages, attire, and other cultural features (6, 7). They are surrounded by other southwest Bantu-speaking groups following an agropastoral subsistence pattern, such as the Nyaneka who live on the Chela plateau to the east of the Namib desert and speak a language of the Nyaneka-Nkhumbi cluster (4), as well as the Ovimbundu who form the largest ethnolinguistic group of Angola and inhabit a vast area to the north of Namibe province, stretching from the Bié Plateau to the Atlantic coastline (Fig. S2) (5). All Bantu groups studied in this paper can firmly be placed within the western stream of the Bantu migrations, based on the historical record and the genealogical position of their languages (Fig. S1D) (5, 8).

The Kwepe are small-stock herders inhabiting areas in and around the Kuroka River mouth (Fig. S2). Before shifting to their present dialect of Kuvale during the first half of the 20th century (Fehn, 2019), their original language Kwadi was recorded by António de Almeida (9) and Ernst Westphal (10–12), who encountered few speakers living near the Kuroka River. During our fieldwork (2013-2014), we not only encountered the descendants of Westphal's informants, but also found two elderly women who had learned the language as children and remembered a considerable amount of lexical items and grammatical phrases. Kwadi has been linked to the Khoe languages spoken further south in the Kalahari Basin (13) and is therefore associated with the dispersal of Khoe-Kwadi languages and pastoralism from eastern Africa (Fig. S1C) (14).

The Kwisi and Twa are two Kuvale-speaking groups with low socio-economic status, who have been suggested to make part of a distinct layer of non-Khoisan hunter-gatherers inhabiting the areas in and around the Namib desert prior to the Bantu expansion (Fig. S2) (6, 9, 15). Unlike the Kwepe, who are clearly associated with a pre-Bantu layer through their historically documented use of the Kwadi language, the Twa and Kwisi have never been recorded as speaking any language other than their current Bantu varieties, making it impossible to speculate about possible affiliations to other ethnolinguistic groups of southern Africa. Both groups are said to have followed a foraging tradition which may also have included the exploration of maritime resources, but now adhere to mixed subsistence strategies ranging from small-stock herding to blacksmithing and sorcery (6). Despite their secluded way of life, we were able to contact and study two Kwisi communities from the regions of Oluheke and Virei in the northern part of the Namib desert, as well as a Twa group living to their south at Onkokoa.

A further small-scale group, the Himba-speaking Tjimba, are sometimes considered to be descended from pre-Bantu foragers (16), but have more commonly been interpreted as impoverished Himba (17). The Tjimba community studied in this paper dwells around the resident Himba population of Iona National Park in the southern part of the Namib desert (Fig. S2).

Regardless of their origins, the Kwepe, Kwisi, Twa and Tjimba communities are united by their high mobility and economic dependence on their more powerful pastoral neighbors, thus following a lifestyle best described as peripatetic (18). They thus resemble the Damara from Namibia who, in historical times, were found orbiting around the Khoekoe-speaking Nama pastoralists whose language they adopted, despite being genetically indistinguishable from the Herero and Himba (3, 19).

With the exception of the !Xun, all groups from southwestern Angola follow a strict matrilineal descent system in which clan labels are inherited from the mother, and livestock is passed to the mother's brother's son, rather than to one's own offspring (17). Our studies on mtDNA in the Himba, Kuvale, Kwepe, Kwisi, Twa and Tjimba have demonstrated that matriclans have a high genealogical consistency in all groups from the Angolan Namib, including the peripatetics (3, 20).

### SI Text 2. *qpAdm* analyses

As previously shown (21), using an adequate set of reference and source populations is critical to accurately estimate admixture proportions in *qpAdm*. While the underlying phylogeny relating all populations does not need to be known, the populations used as references cannot share drift with the target populations more recently than the sources.
When testing different admixture models for ancient and present-day African groups, Skoglund and colleagues (22) selected nine populations as possible sources: Mbuti, Dinka, Mende, South Africa 2000BP, Tanzania Luxmanda 3100BP, Ethiopia 4500BP, Levant Neolithic (PPNB), Anatolia Neolithic, and Iran Neolithic. In each test, one to three populations were chosen iteratively from this source pool, while the remaining were defined as "outgroups" or "right populations", together with the additional groups: Denisova, Loschbour, Ust Ishim, Georgian, Iranian, Greek, Punjabi, Orcadian, Ami, and Mixe. While this diverse set of populations (hereafter referred to as the Skoglund set) might be among the best possible choices to test admixture models across the whole African continent, it might not be the most suitable at a regional level.

Several studies have shown that southern Africa was settled by three main layers of people: foragers speaking Kx'a and Tuu languages, Eastern African pastoralists, and Bantu-speaking farmers with origins in West-Central Africa (12, 14, 19, 23–27). Accordingly, when using the pool of sources from the Skoglund set, the best models of admixture for southern African populations often include as best proxies South Africa 2000BP, Tanzania Luxmanda 3100BP, and the Mende (each representing one of the layers). Yet, for some southern African populations, the best model includes South Africa 2000BP and the Dinka – an Eastern African population that in ADMIXTURE analyses has a mixed profile of East- and West-African related ancestry (Fig. S6 and Fig. S13A-B). These results can be partially explained by a preference to report only the model with the minimum number of sources if the p-value of that model already reached a predefined threshold (usually 0.05 or 0.01) (Fig. S13A-B) and might also be affected by a violation of the model requirements. If the West African ancestry of the Dinka is at least partially the result of admixture with East Bantu groups, then the Dinka is not only serving as a proxy for an Eastern African lineage in the model but will also share drift with the test groups more recently than the Mende, who represent the West African ancestry (22) but are not actually a Bantu-speaking group. To avoid a model violation due to the inclusion of both Mende and Dinka, we tested admixture models using the following modifications: i) addition of Ovimbundu, a Bantu-speaking population whose profile better represents the West African ancestry that spread into southern Africa, to the sources, ii) use of the more distantly related Mende only as a reference population, iii) and removal of the Dinka from all models. We display the results of the best models according to different criteria, and for comparison also show the corresponding results using the Skoglund set (Fig. S13, Table S3 and S4).

Based on the modified set of populations, the southern African groups that were previously described as a mixture of Mende and Dinka are now composed by all or a subset of three sources:

South Africa 2000BP, Tanzania Luxmanda 3100BP, and Ovimbundu (Fig. S13E-H). Likewise, the modifications had an impact on the results for some Angolan groups. When the Skoglund set is used (Fig. S13A-D) the marginalized groups from Angola are described as a mixture of three components, represented by South Africa 2000BP, Tanzania Luxmanda 3100BP, and the Mende. However, other populations from Angola are composed of West African ancestry (Mende) and ancestry from Central African Rainforest Hunter-gatherers (Mbuti) but no ancestry related to southern African foragers (South Africa 2000BP). These results portray a sharp dichotomy between the nature of introgression in marginalized and non-marginalized neighboring groups and do not align with the remaining genetic analyses (Fig. S12). Instead, they are likely the product of model violations and unsuitable source-reference pools. The use of our modified set of populations (Fig. S13E-H) shows that the Angolan groups are best described by varying amounts of the same three ancestry components that are found elsewhere in southern Africa. More complex models with additional sources (>3) were not tested and thus cannot be excluded.

### SI Text 3. Admixture dating analyses

Both Wavelets and GLOBETROTTER can estimate up to two admixture events. In Wavelets, the estimation of the most recent event is done by treating all blocks originating from previous events as part of a single population (source 1). In southwestern Angolan and northwestern Namibian groups, these blocks correspond to those identified by RFMix as being of East and Southern African origin. Source 2 would then be composed of all blocks identified as Bantu by RFMix. On average, the proportion of source 1 in southwestern Angolan and northwestern Namibian groups ranges from 13% and 26% (Table S5) - a percentage for which admixture times could be successfully recovered based on simulations (28). To estimate the earlier event, the blocks of the most recent ancestry are masked and the signal of admixture is analyzed using the remaining part of the genome (28). However, since each of the ancestries corresponding to the earlier event in southwestern Angolan and northwestern Namibian groups comprise only a small fraction of the genome (below what has been previously tested) it is likely that the estimates are unreliable. The admixture dates (Table S5) range from 3200 years in the Kwisi to 6000 years in the Damara - a time period that largely predates the arrival of the East African-related ancestry in southern Africa based on ancient DNA studies. Additionally, the admixture dates are highest (and further away from reasonable values) for the groups that have the lowest amount of pre-Bantu ancestries. Therefore, even though we report all of our estimates, our discussion focuses exclusively on the most recent event.

When estimating admixture in the Bantu-speaking groups from southwestern Angola and northwestern Namibia using GLOBETROTTER, only one admixture signal was recovered (Table S6). The single sources that best match the ancestries involved in this admixture event are the Ovambo (a Bantu-speaking population) and the Nama or the ǂKhomani – two southern African populations that display the highest ratio of East to Southern African ancestry (Fig. 4). Since all of the present-day populations from southern Africa display some amount of East African introgression, none of the surrogate populations used as potential sources in the GLOBETROTTER analysis represent exclusively the autochthonous southern African ancestry but are already the product of an admixture event. Thus, given the lack of suitable unadmixed surrogates and the minor proportion of East and Southern African ancestry in the tested groups, it is not surprising that GLOBETROTTER fails to detect the older admixture event. The admixture dates inferred for the most recent event are consistent across all tested methods (Fig. S14).

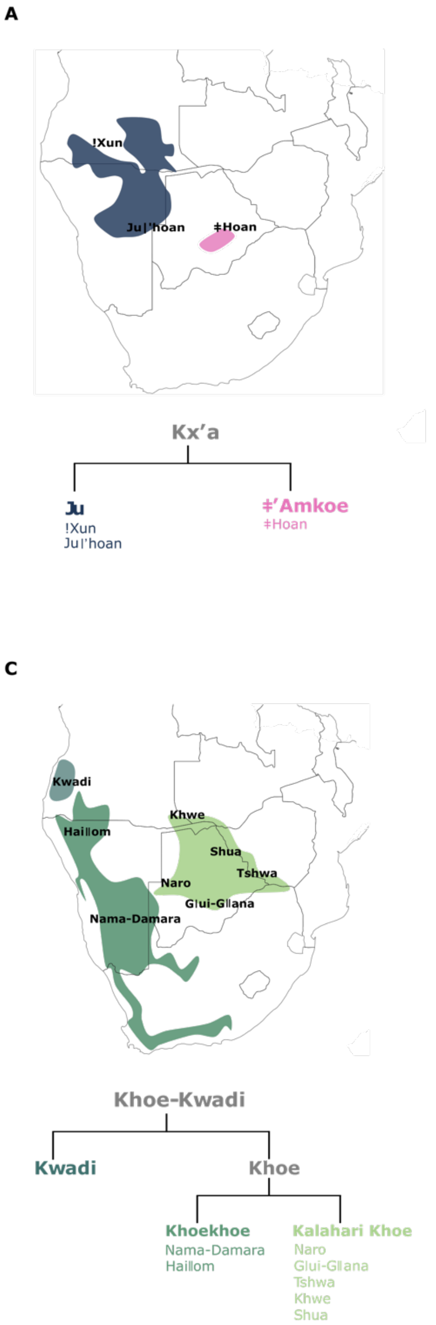


**Figure S1. Subclassification and historically attested distribution of the four pre-colonial language families of southern Africa.** A) Kx'a (29); B) Tuu (30, 31); C) Khoe-Kwadi (31, 32); D) Bantu (8, 25). The approximate locations of languages spoken by southern African populations analyzed in this study are indicated in the maps.

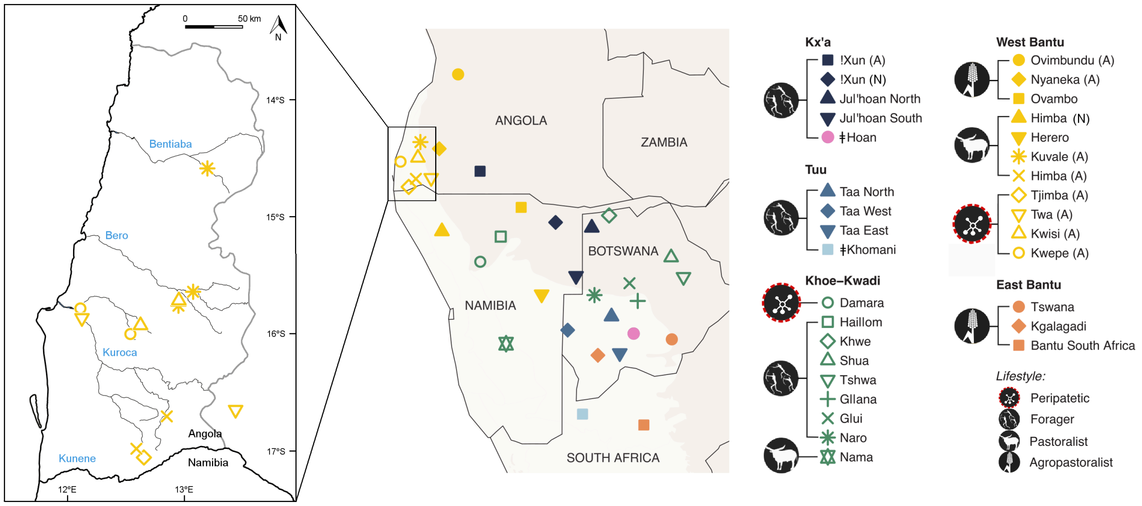


**Figure S2. Map of southern Africa showing the location of the newly studied Angolan groups and other southern African groups included in the reference dataset.** The map displays the approximate sample locations. In the inset map, the Angolan Namib province is delimited by a gray contour, country borders are shown in black, and the names of the main intermittent rivers are indicated in blue. All samples from Angola are marked with “(A)” in the legend, while samples from the same ethnic group collected in Namibia are marked with “(N)”.

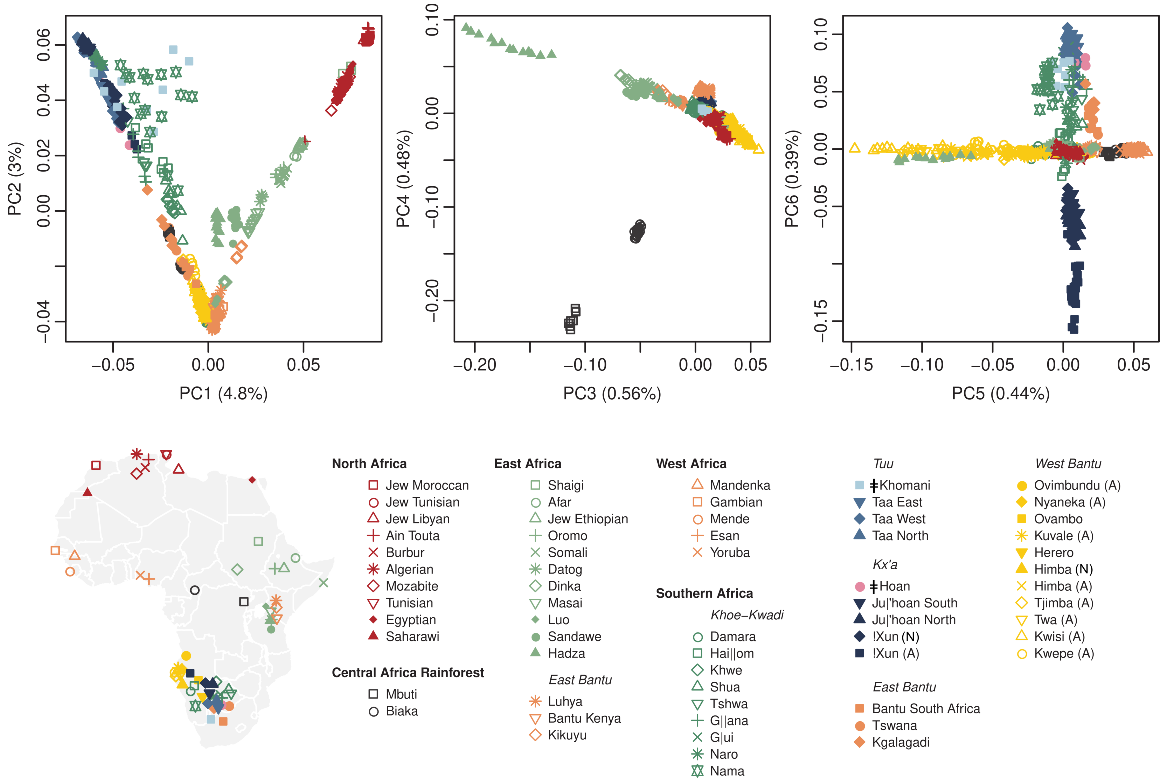


**Figure S3. PCA of African populations showing the first six PCs.** The map displays the approximate sample locations. In the legend, geographic labels are shown in bold and relevant linguistic divisions are shown in italic; all samples from Angola are marked with "(A)", while samples from the same ethnic group collected in Namibia are marked with "(N)".

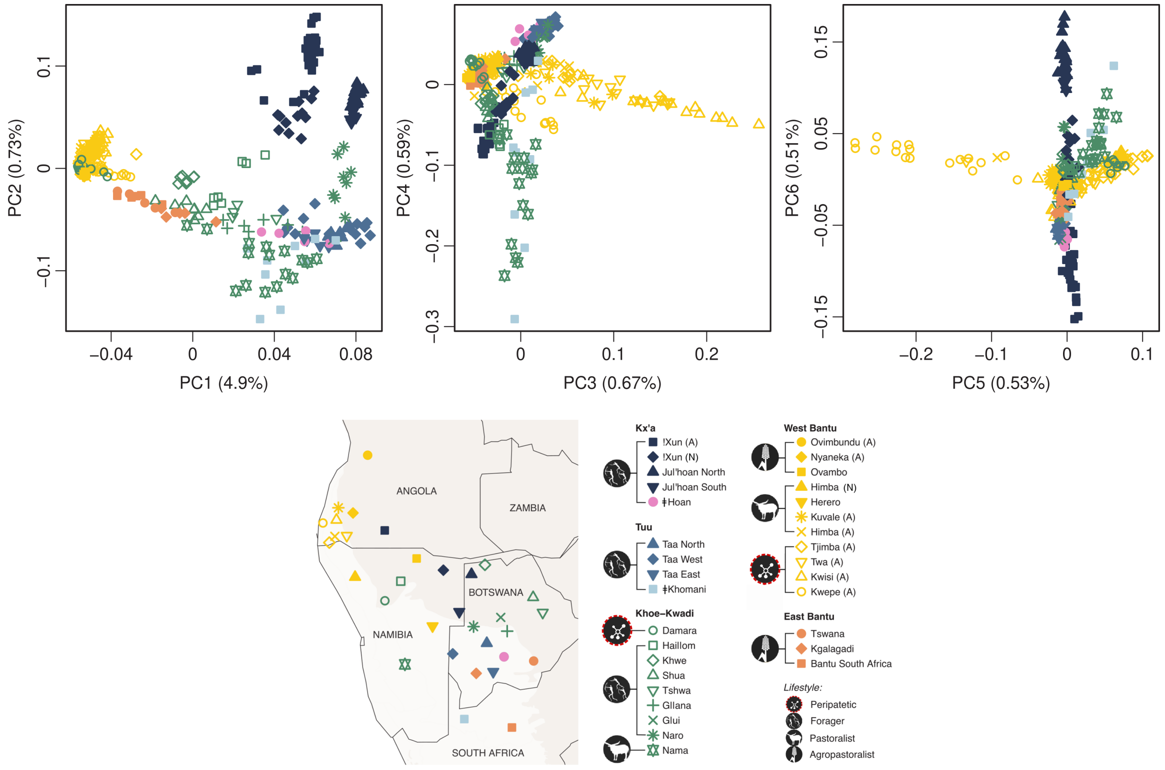


**Figure S4. PCA of southern African populations showing the first six PCs.** The map displays the approximate sample locations. The lightest background color shows the desert and xeric shrublands biome (Olsen et al., 2001). In the legend, all samples from Angola are marked with "(A)", while samples from the same ethnic group collected in Namibia are marked with "(N)".

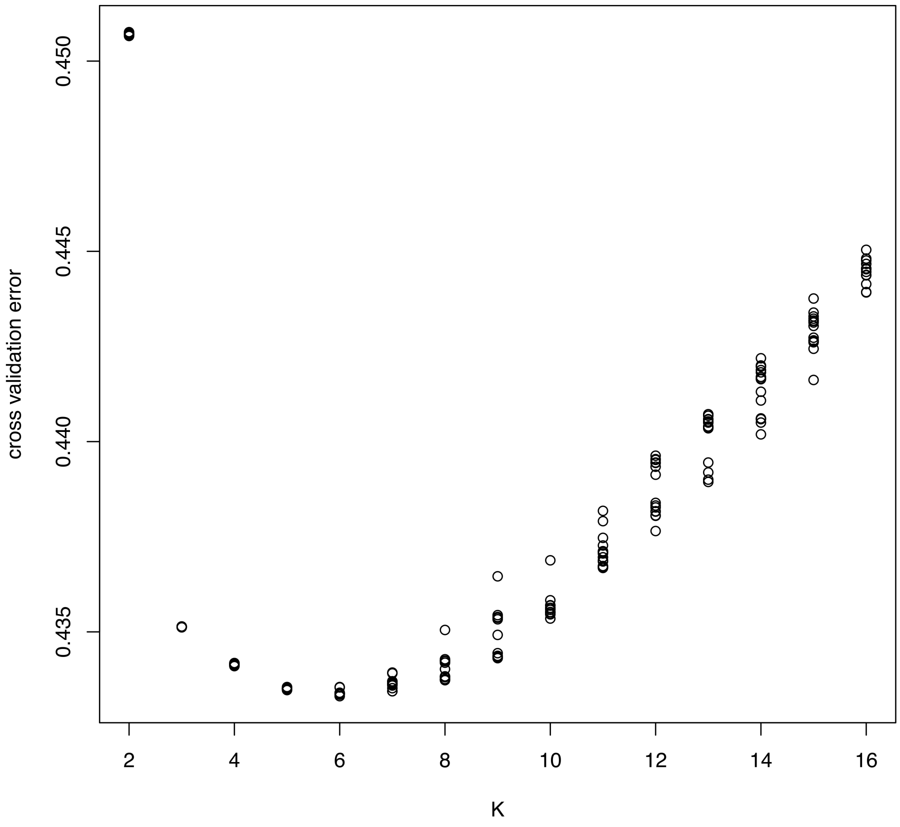


**Figure S5. Cross-validation error for 15 ADMIXTURE runs under each model (K = 2 to K = 16).**

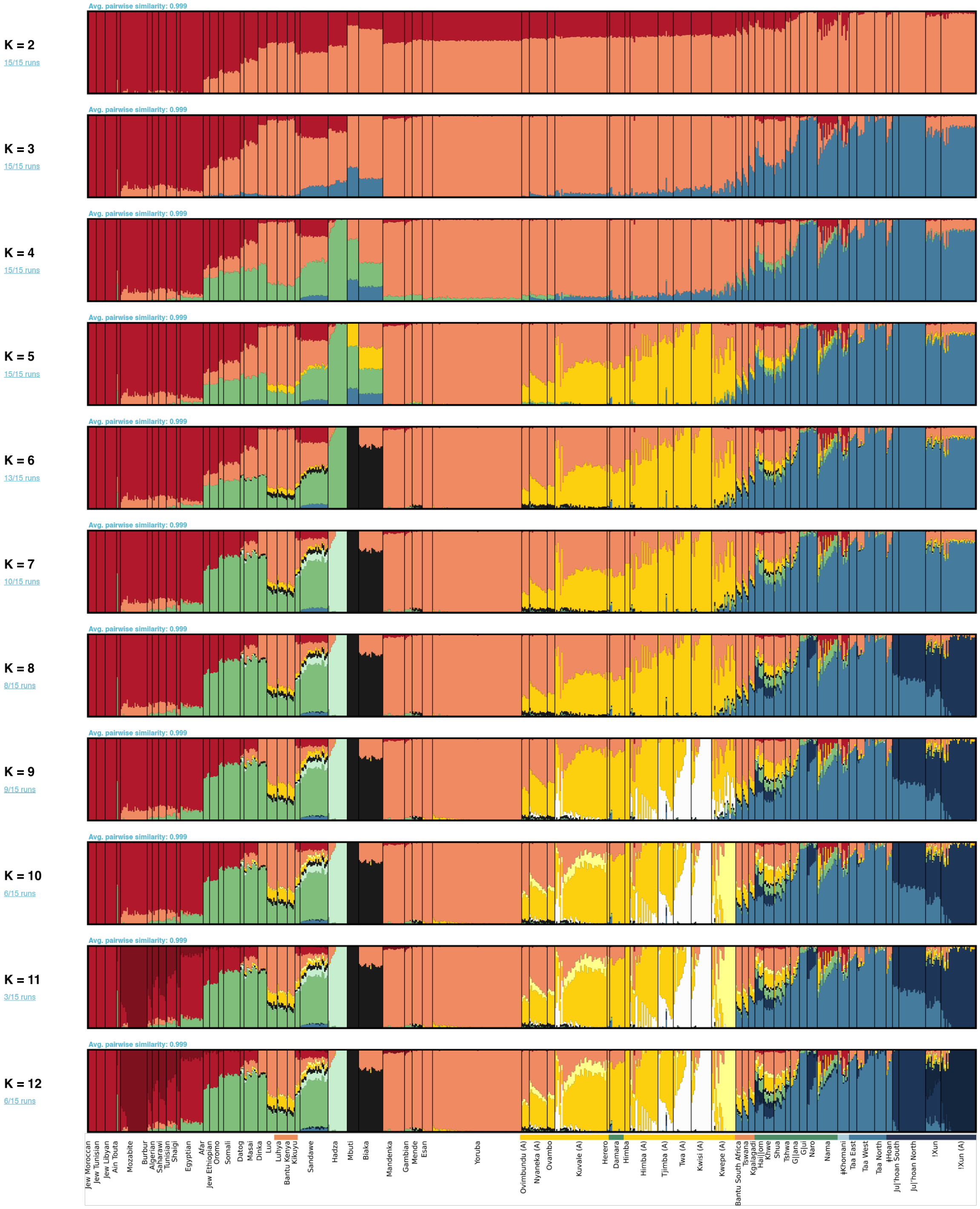


**Figure S6. ADMIXTURE results for K = 2 to K = 12.**

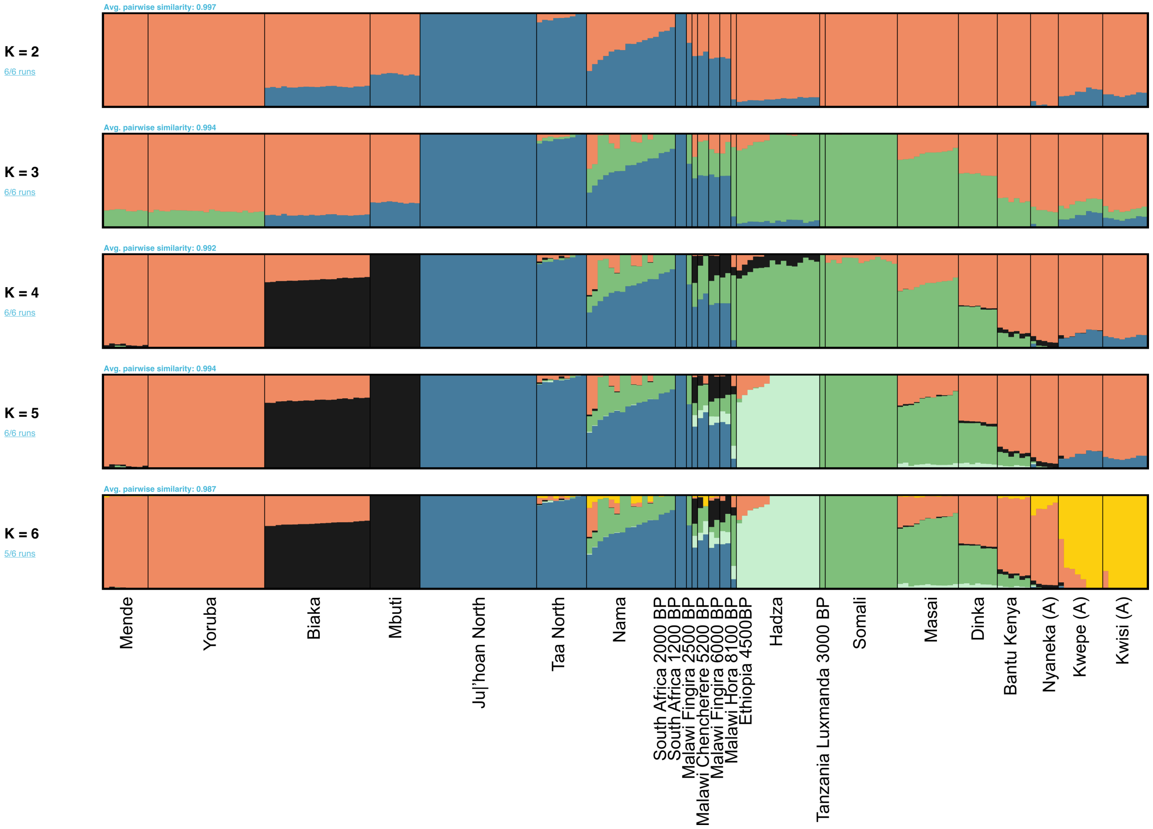


**Figure S7. DyStruct results for K = 2 to K = 6.** This analysis includes relevant ancient individuals from East and Southern Africa and a subset of the present-day groups used in the ADMIXTURE analysis. The Angolan groups included here were downsampled to lower the impact of large sample sizes in the clustering analysis.

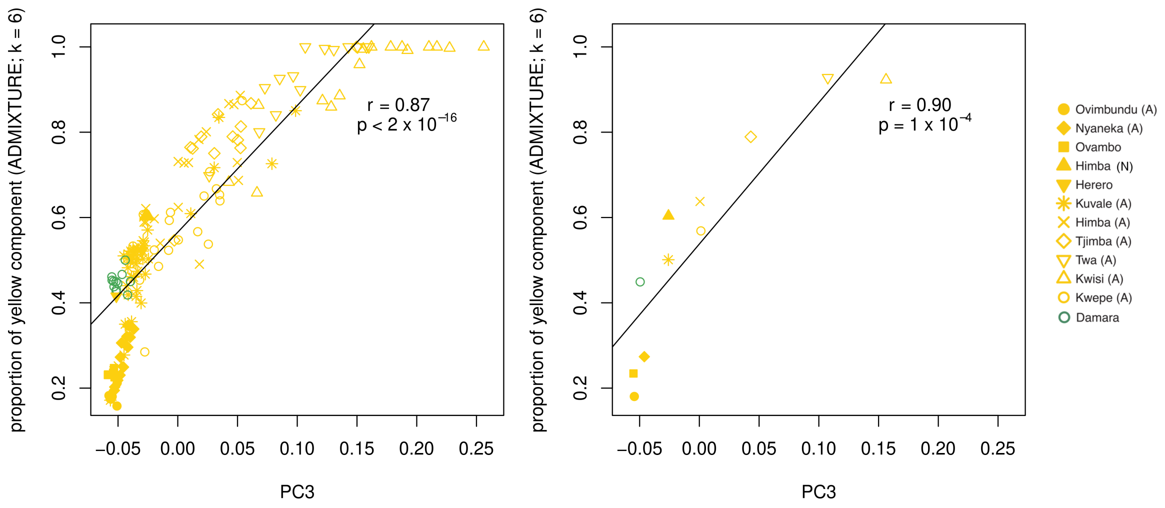


**Figure S8. ADMIXTURE results vs PC3 scores in southwestern Bantu-speaking groups and the Damara.** The proportion of yellow component at K=6 and the PC3 scores are shown for each individual and averaged by population on the left and right panels, respectively. The black line shows the linear regression. r – Pearson correlation coefficient; p – P value.

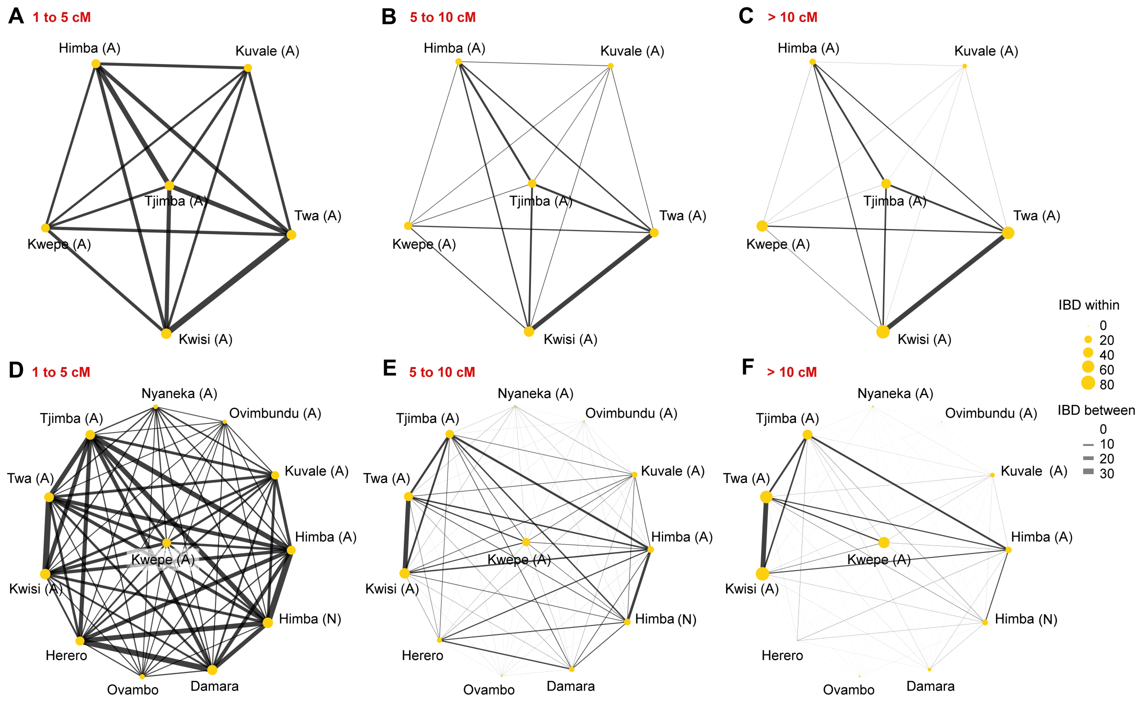


**Figure S9. Identity-by-descent (IBD) sharing among groups from the Angolan Namib Desert (A-C) and among all southwestern Bantu-speaking groups plus the Damara (D-F).** The circle size and line width are proportional to the mean total IBD length (cM) shared by two individuals from the same group (IBD within) or from different groups (IBD between) within each length category. The 1-5, 5-10, and >10 cM length categories roughly correspond to time intervals of 1,500-2,500 years ago, 500-1,500 years ago, and 0-500 years ago, respectively (33).

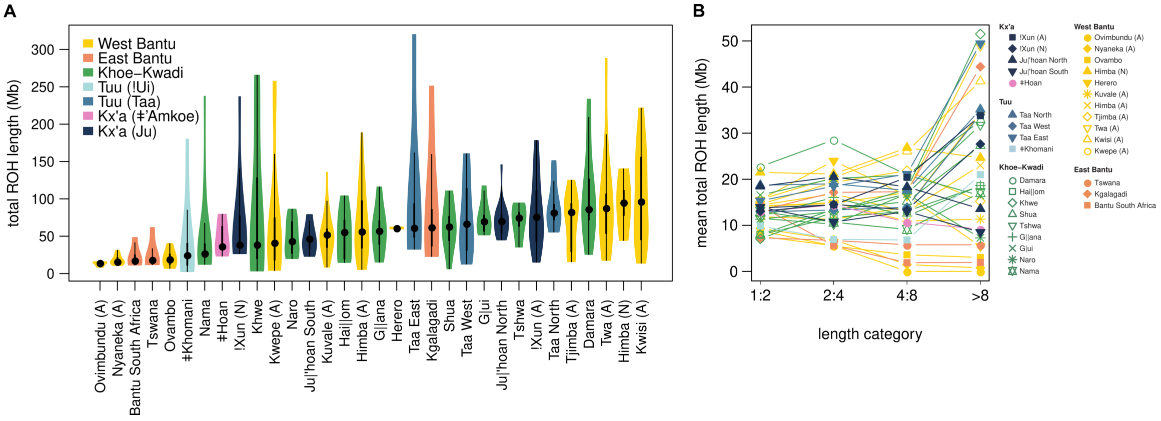


**Figure S10. Runs of homozygosity (ROH) in southern Africa.** A) Violin plots showing the distribution of total ROH length (Mb) per population. The thick and thin black lines represent the interquartile range and the upper and lower adjacent values, respectively; the central dot shows the median. B) Mean total ROH length per population for different length categories.

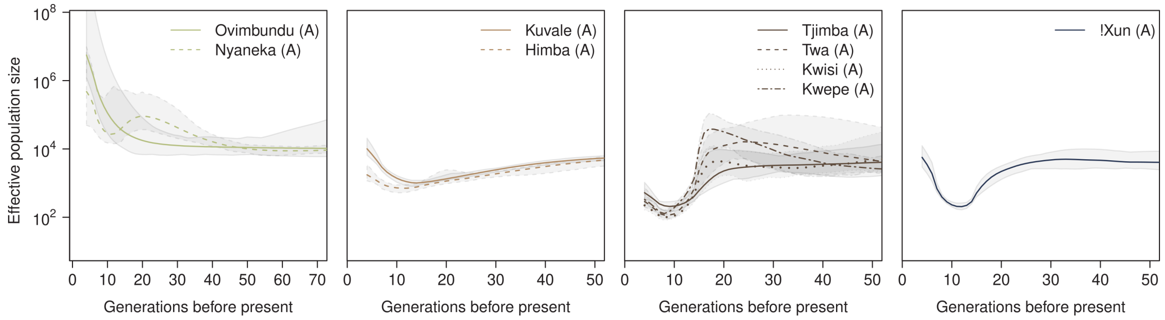


**Figure S11. Effective population size (*Ne*) across time.** *Ne* estimates based on IBD sharing within groups from Angola. The plot is truncated at 4 generations before present due to the higher uncertainty associated with Ne estimates for very recent times.

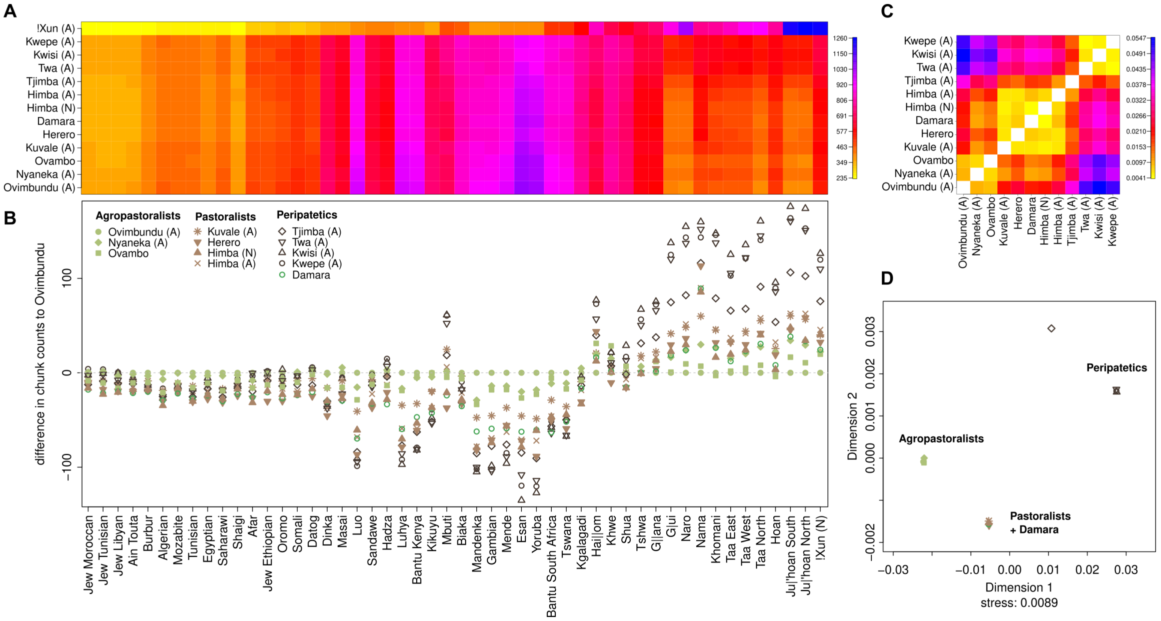

**Figure S12. Chromopainter profiles for southwestern Angolan and northwestern Namibian groups, when only distantly related African groups are used as donors.** A) Chromopainter coancestry matrix. The color gradient indicates the average number of DNA chunks a group copies from the donor populations. B) Differences between the number of DNA chunks copied by each group and the Ovimbundu - used here as a baseline. C) Average distance (TVDxy) between the copying profiles. D) Multidimensional scaling calculated on the TVDxy distances.

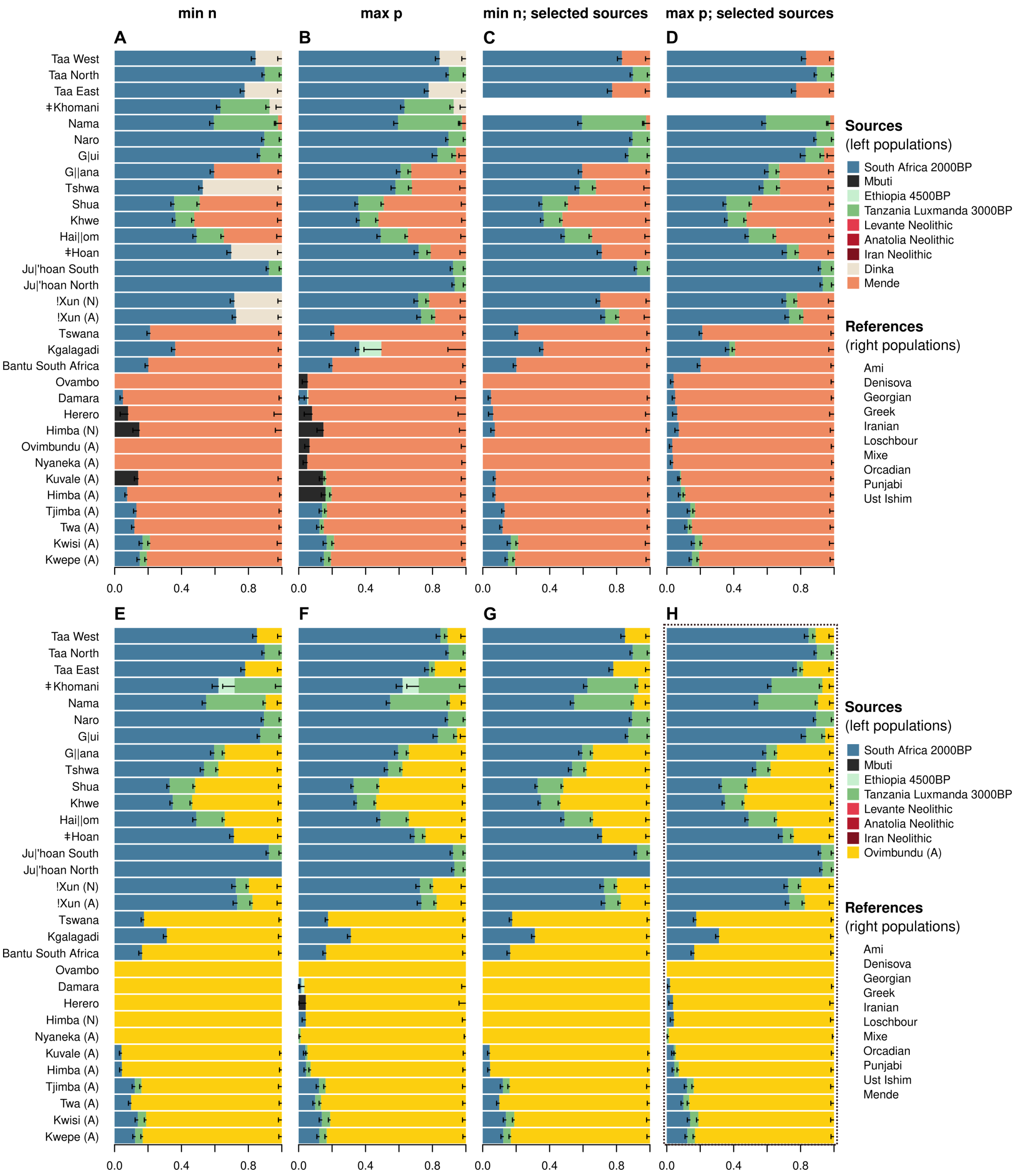


**Figure S13. Ancestry proportions estimated with *qpAdm* in southern African populations under different mixture models.** Models are tested using the reference and source populations as in Skoglund et al., (2017) (A-D) or using a modified set of populations (E-H). The best fit models are displayed according to different criteria: min n - minimum number of sources needed to obtain a model that fits (A, C, E, G); max p – model returning the highest P-value (B, D, F, H); selected sources – only considers models that include as sources the Mende/Ovimbundu, South Africa 2000BP, and Tanzania Luxmanda 3000 BP (C, D, G, H). H) Results displayed in Figure 4. Standard errors (black bars) were calculated with a weighted block jackknife. In the legend, *Sources* correspond to the full pool of sources; in a rotating strategy, each of them also served as references for some models.

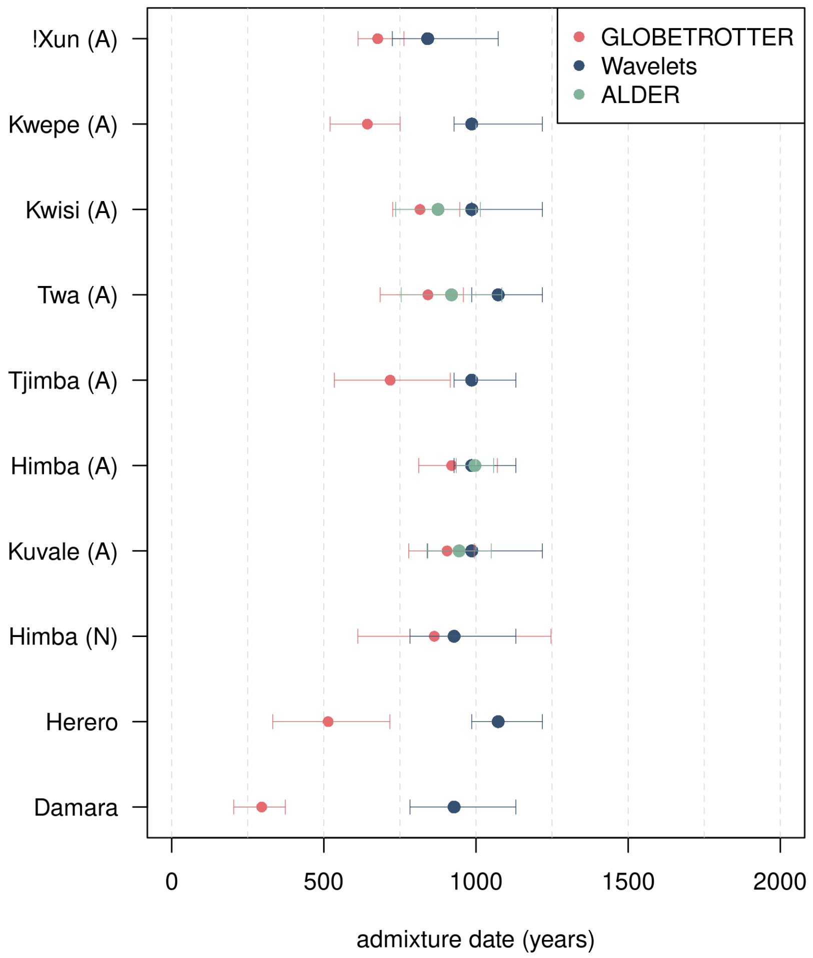


**Figure S14. Admixture time estimates between Bantu and pre-Bantu ancestries in admixed populations from southwestern Angola and northwestern Namibia**

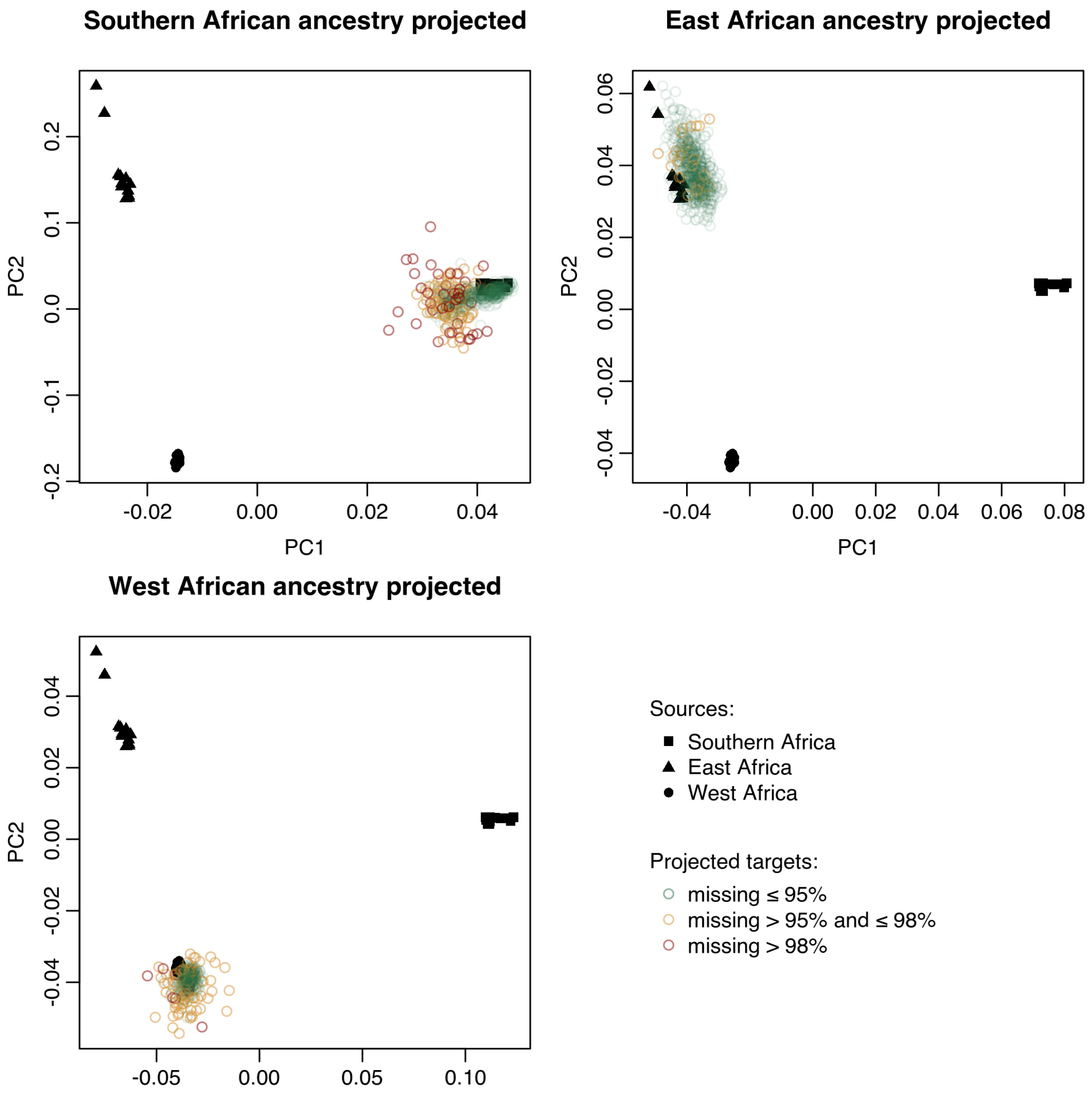


**Figure S15. Evaluation of ancestry assignment and masking procedures.** The PCA was built with the source populations used in the RFMix analysis (see Methods). In each panel, the individuals from southern African groups are projected after masking their non-targeted ancestries and are colored according to the amount of missing data.

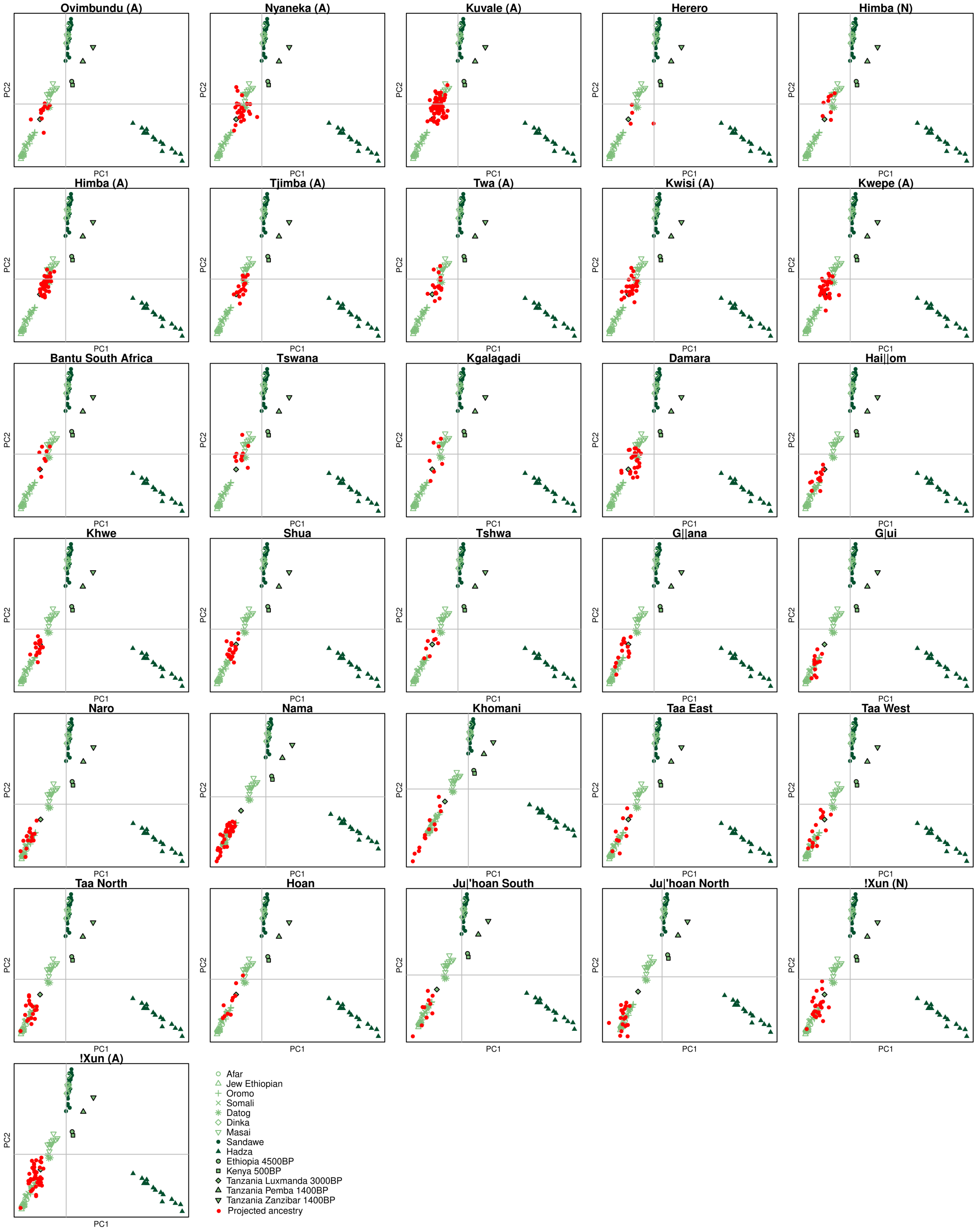


**Figure S16. PCA of East African-specific ancestry.** The PCA was built with present-day populations from East Africa. In each panel, the individuals from each southern African group are projected (red points), after masking their non-East African ancestries.

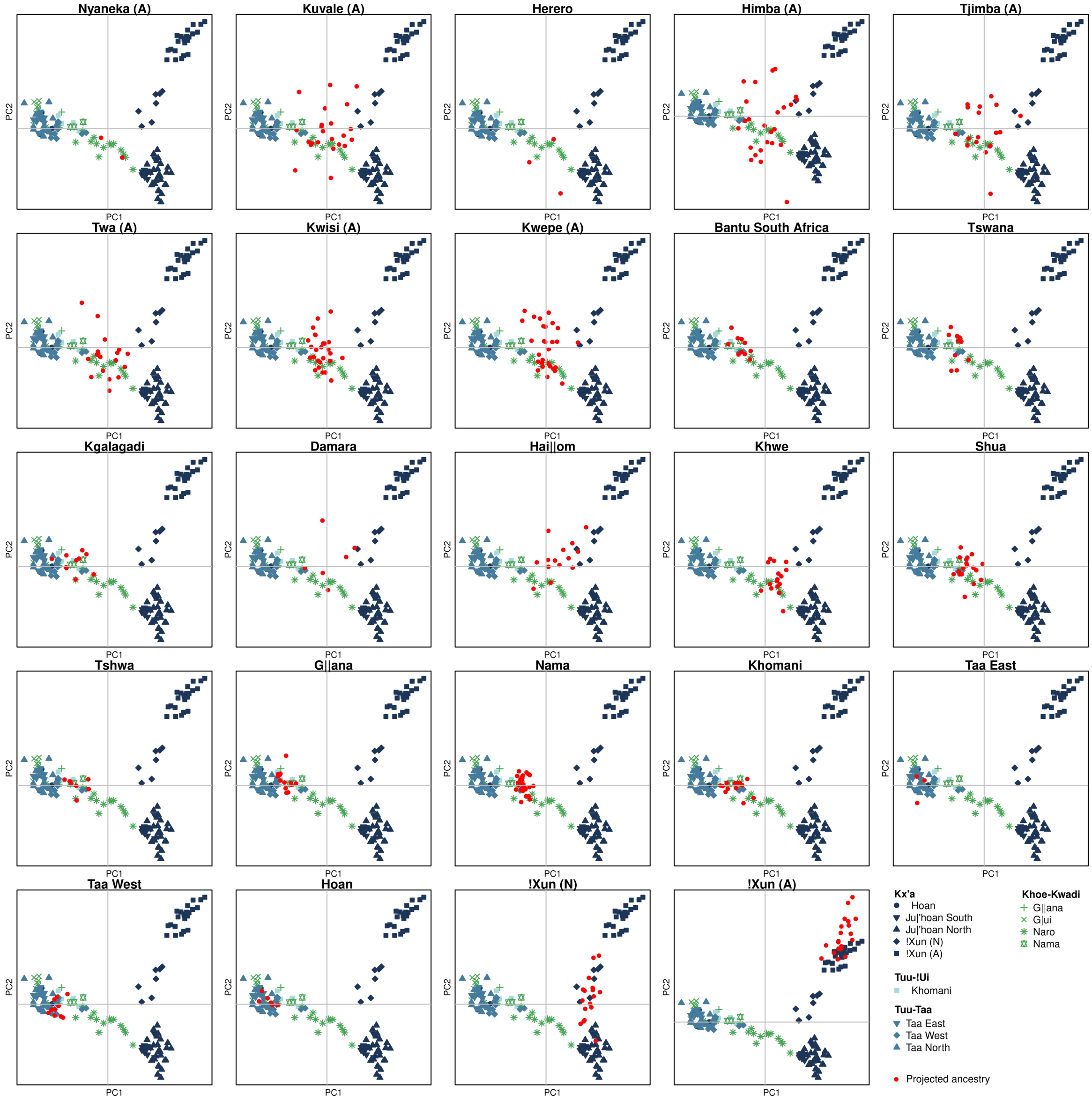


**Figure S17. PCA of autochthonous southern African ancestry.** The PCA was built with southern African individuals that have <25% of missing data after masking the non-southern African ancestries. In each panel, the individuals from each southern African group that have >25% of missing data after masking are projected (red points). Since individuals within a single southern African population can have different levels of missing data, some panels contain both projected and non-projected individuals from the same population.

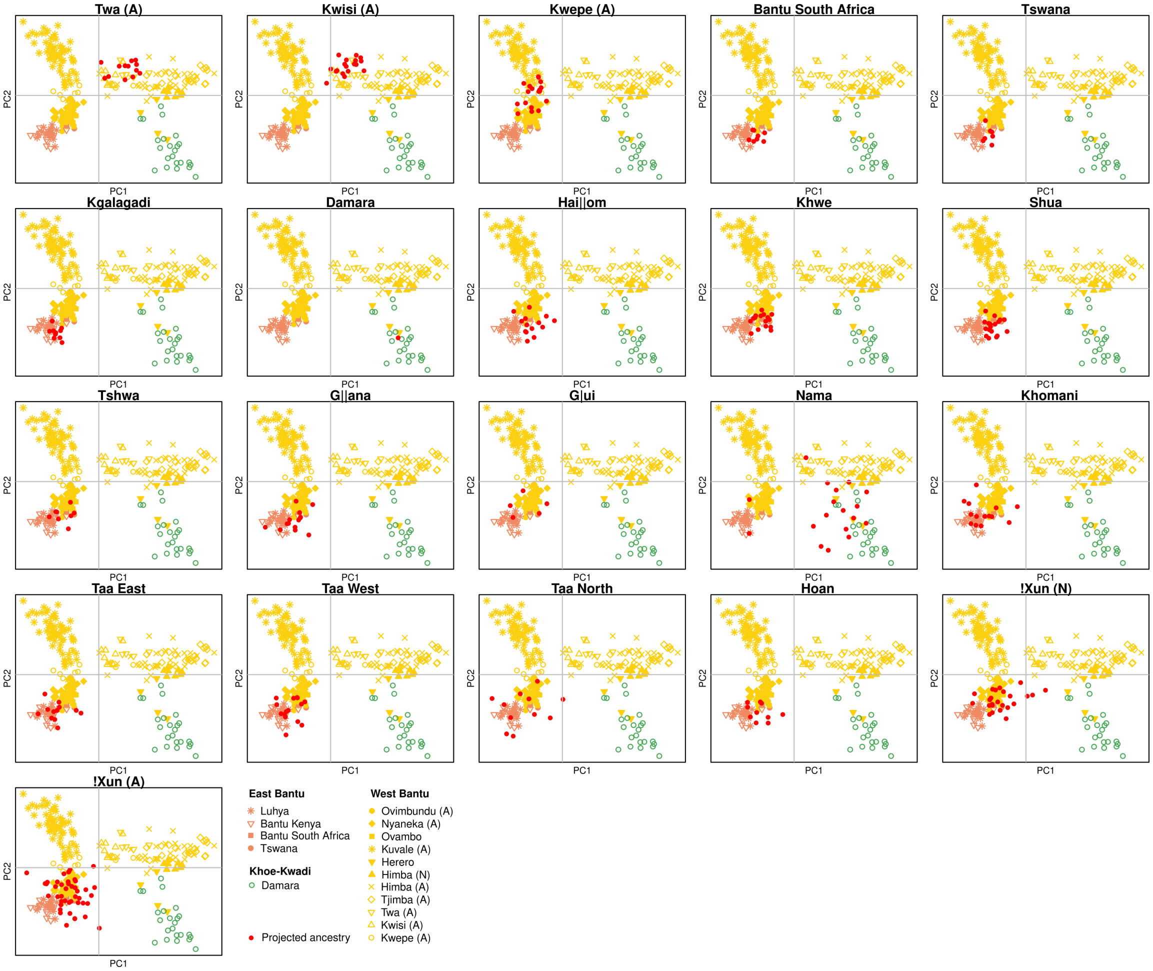


**Figure S18. PCA of Bantu-specific (West African) ancestry.** The PCA was built with southern African individuals and East African Bantu speakers that have <25% of missing data after masking the non-Bantu ancestries. In each panel, the individuals from each southern African group that have >25% of missing data after masking are projected (red points). Since individuals within a single southern African population can have different levels of missing data, some panels contain both projected and non-projected individuals from the same population.

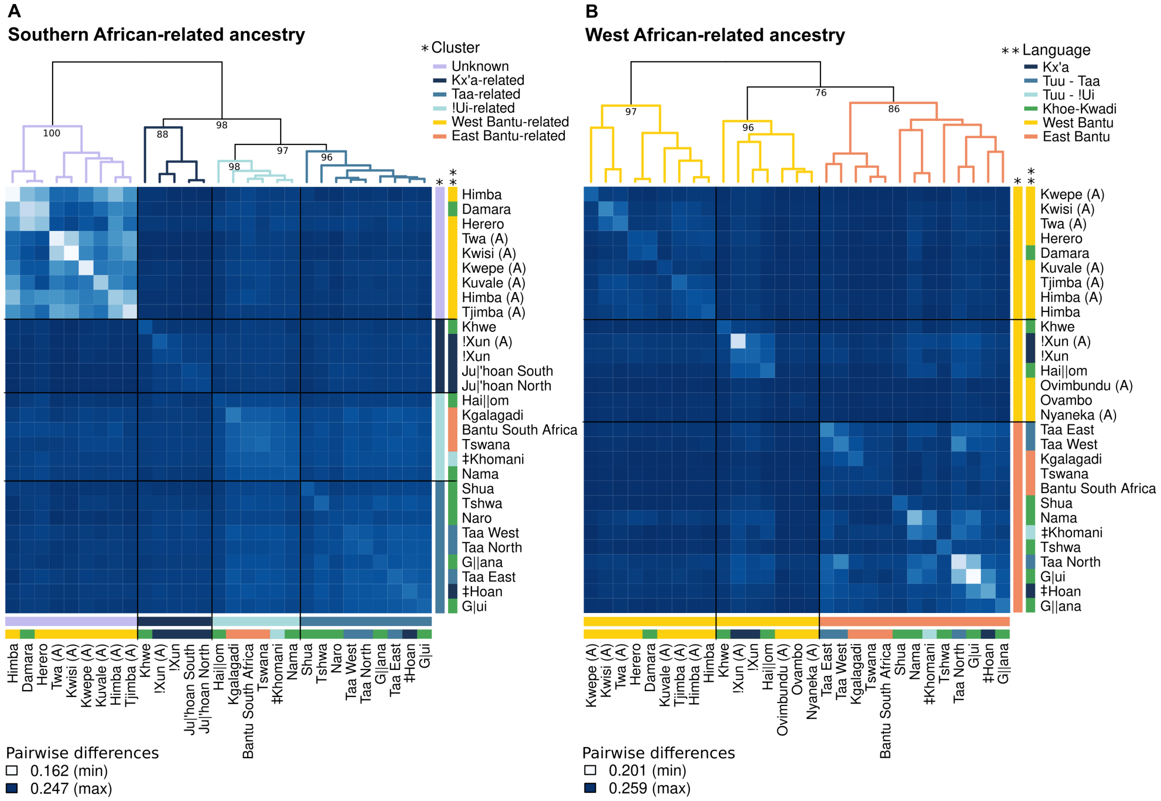


**Figure S19. Matrix of pairwise differences calculated on the southern African-related (A) and Bantu-related (B) ancestries.** The color gradient shows the average pairwise differences calculated using individuals with <10% of missing data after masking. The dendrogram shows the result of the clustering analysis, with Approximately Unbiased (AU) p-values computed by multiscale bootstrap resampling. Major cluster assignment (*) and language affiliation (**) for each southern African population are represented by different colors in the bars next to the population labels.

(plots shown in the next pages)

**Figure S20. Ancestry-specific IBD sharing.** The stacked barplots show the sum of IBD lengths (cM) from a given length category that are shared on average between individuals from a target population identified in the panel title and individuals from populations indicated in the y-axis. The colors along the y-axis represent the linguistic affiliation of each population, according to the legend. In the barplots, the colors orange, blue, and green correspond to West, southern, and East African-related ancestries, respectively. The amount of IBD fragments of ambiguous origin (see methods) are shown in grey.

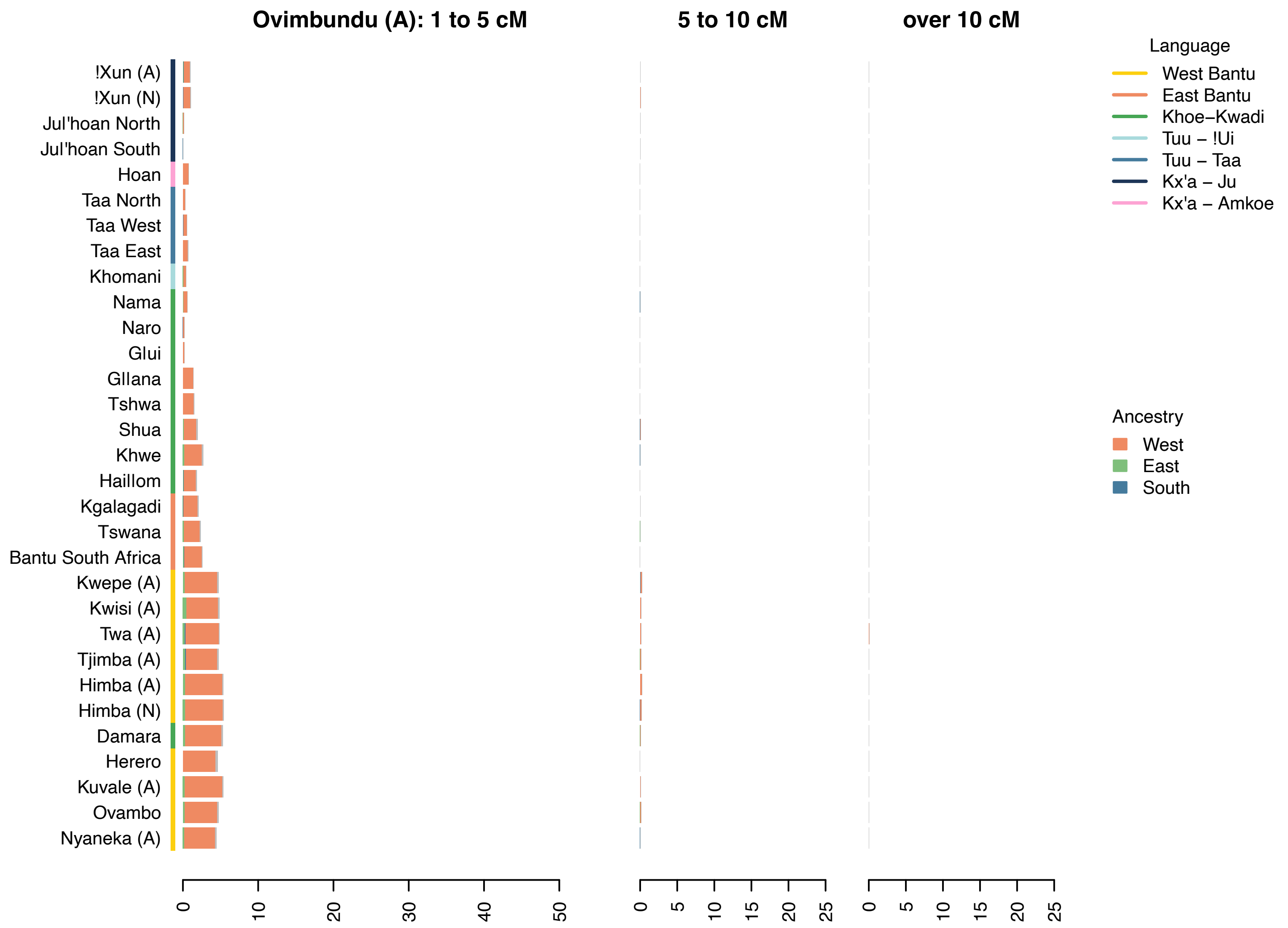

Ovimbundu (A): 1 to 5 cM
5 to 10 cM
over 10 cM
!Xun (A)
!Xun (N)
Jul'hoan North
Jul'hoan South
Hoan
Taa North
Taa West
Taa East
Khomani
Nama
Naro
Glui
Gllana
Tshwa
Shua
Khwe
Haillom
Kgalagadi
Tswana
Bantu South Africa
Kwepe (A)
Kwisi (A)
Twa (A)
Tjimba (A)
Himba (A)
Himba (N)
Damara
Herero
Kuvale (A)
Ovambo
Nyaneka (A)
0 10 20 30 40 50
0 5 10 15 20 25
0 5 10 15 20 25
Language
West Bantu
East Bantu
Khoe–Kwadi
Tuu – !Ui
Tuu – Taa
Kx'a – Ju
Kx'a – Amkoe
Ancestry
West
East
South


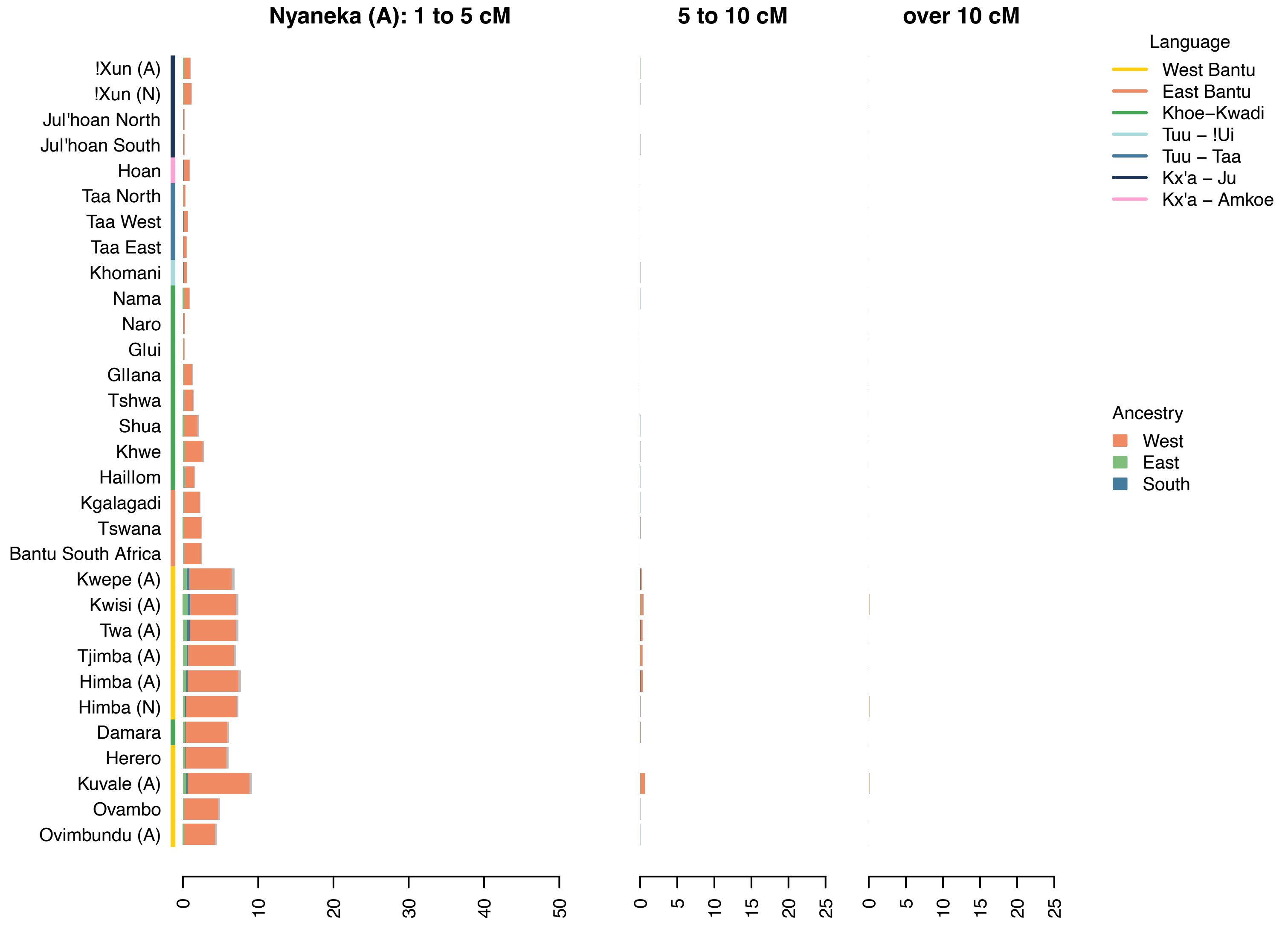

Nyaneka (A): 1 to 5 cM
5 to 10 cM
over 10 cM
!Xun (A)
!Xun (N)
Jul'hoan North
Jul'hoan South
Hoan
Taa North
Taa West
Taa East
Khomani
Nama
Naro
Glui
Gllana
Tshwa
Shua
Khwe
Haillom
Kgalagadi
Tswana
Bantu South Africa
Kwepe (A)
Kwisi (A)
Twa (A)
Tjimba (A)
Himba (A)
Himba (N)
Damara
Herero
Kuvale (A)
Ovambo
Ovimbundu (A)
0 10 20 30 40 50
0 5 10 15 20 25
0 5 10 15 20 25
Language
West Bantu
East Bantu
Khoe–Kwadi
Tuu – !Ui
Tuu – Taa
Kx'a – Ju
Kx'a – Amkoe
Ancestry
West
East
South

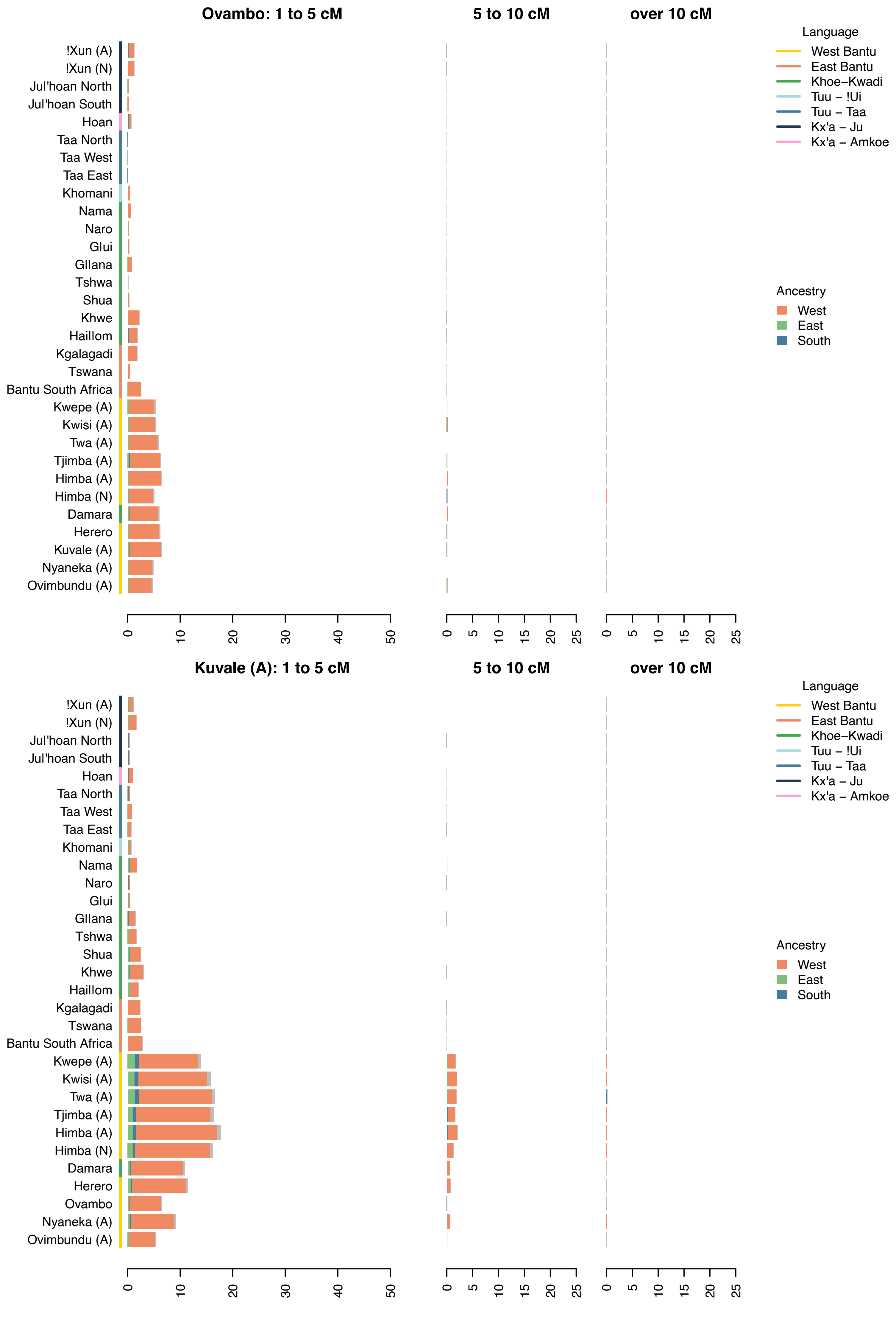

Ovambo: 1 to 5 cM
5 to 10 cM
over 10 cM
!Xun (A)
!Xun (N)
Jul'hoan North
Jul'hoan South
Hoan
Taa North
Taa West
Taa East
Khomani
Nama
Naro
Glui
Gllana
Tshwa
Shua
Khwe
Haillom
Kgalagadi
Tswana
Bantu South Africa
Kwepe (A)
Kwisi (A)
Twa (A)
Tjimba (A)
Himba (A)
Himba (N)
Damara
Herero
Kuvale (A)
Nyaneka (A)
Ovimbundu (A)
0 10 20 30 40 50
0 5 10 15 20 25
0 5 10 15 20 25
Language
West Bantu
East Bantu
Khoe–Kwadi
Tuu – !Ui
Tuu – Taa
Kx'a – Ju
Kx'a – Amkoe
Ancestry
West
East
South
Kuvale (A): 1 to 5 cM
5 to 10 cM
over 10 cM
!Xun (A)
!Xun (N)
Jul'hoan North
Jul'hoan South
Hoan
Taa North
Taa West
Taa East
Khomani
Nama
Naro
Glui
Gllana
Tshwa
Shua
Khwe
Haillom
Kgalagadi
Tswana
Bantu South Africa
Kwepe (A)
Kwisi (A)
Twa (A)
Tjimba (A)
Himba (A)
Himba (N)
Damara
Herero
Ovambo
Nyaneka (A)
Ovimbundu (A)
0 10 20 30 40 50
0 5 10 15 20 25
0 5 10 15 20 25
Language
West Bantu
East Bantu
Khoe–Kwadi
Tuu – !Ui
Tuu – Taa
Kx'a – Ju
Kx'a – Amkoe
Ancestry
West
East
South

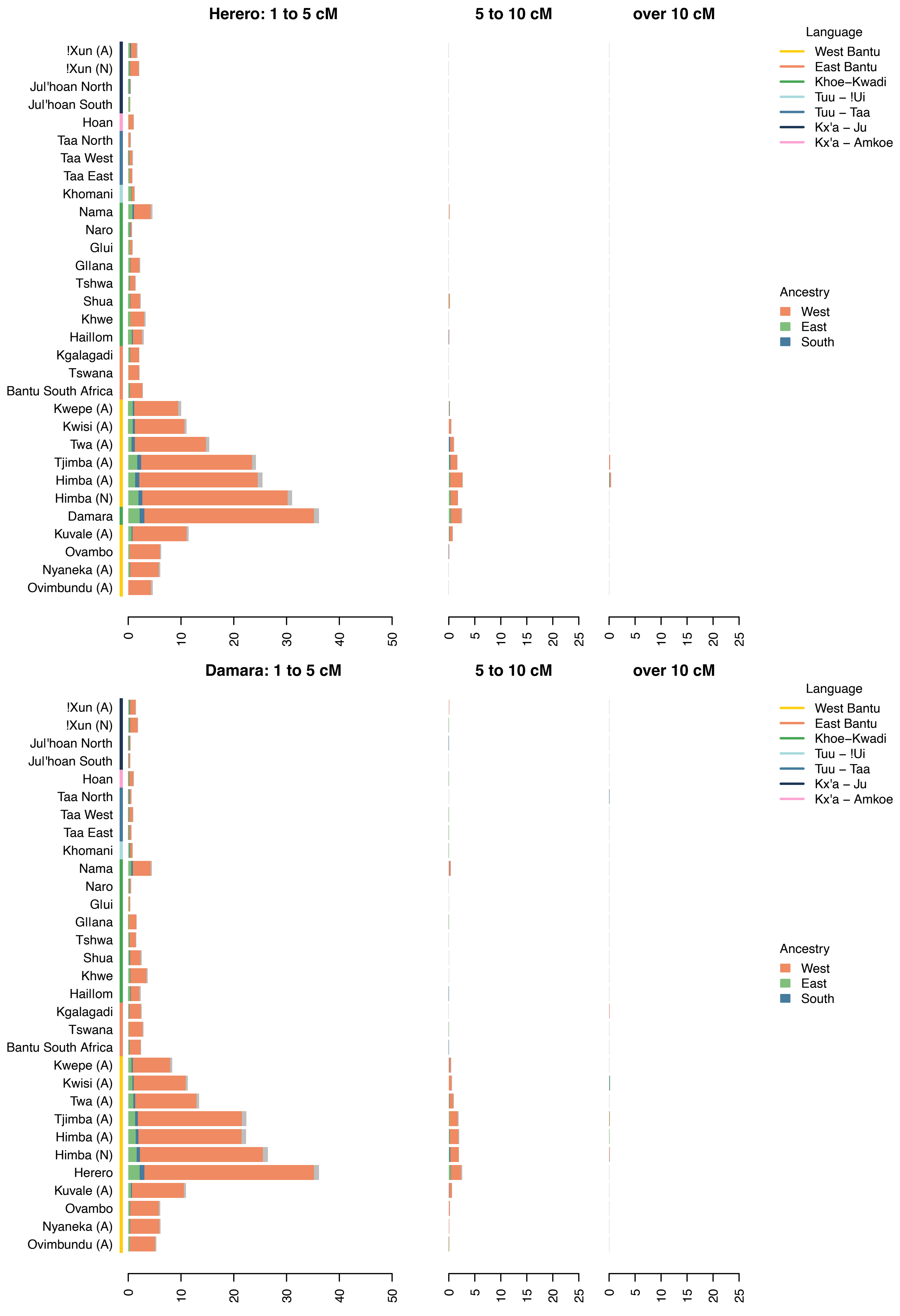
Herero: 1 to 5 cM
5 to 10 cM
over 10 cM
!Xun (A)
!Xun (N)
Jul'hoan North
Jul'hoan South
Hoan
Taa North
Taa West
Taa East
Khomani
Nama
Naro
Glui
Gllana
Tshwa
Shua
Khwe
Haillom
Kgalagadi
Tswana
Bantu South Africa
Kwepe (A)
Kwisi (A)
Twa (A)
Tjimba (A)
Himba (A)
Himba (N)
Damara
Kuvale (A)
Ovambo
Nyaneka (A)
Ovimbundu (A)
0 10 20 30 40 50
0 5 10 15 20 25
0 5 10 15 20 25
Language
West Bantu
East Bantu
Khoe–Kwadi
Tuu – !Ui
Tuu – Taa
Kx'a – Ju
Kx'a – Amkoe
Ancestry
West
East
South
Damara: 1 to 5 cM
5 to 10 cM
over 10 cM
!Xun (A)
!Xun (N)
Jul'hoan North
Jul'hoan South
Hoan
Taa North
Taa West
Taa East
Khomani
Nama
Naro
Glui
Gllana
Tshwa
Shua
Khwe
Haillom
Kgalagadi
Tswana
Bantu South Africa
Kwepe (A)
Kwisi (A)
Twa (A)
Tjimba (A)
Himba (A)
Himba (N)
Herero
Kuvale (A)
Ovambo
Nyaneka (A)
Ovimbundu (A)
0 10 20 30 40 50
0 5 10 15 20 25
0 5 10 15 20 25
Language
West Bantu
East Bantu
Khoe–Kwadi
Tuu – !Ui
Tuu – Taa
Kx'a – Ju
Kx'a – Amkoe
Ancestry
West
East
South

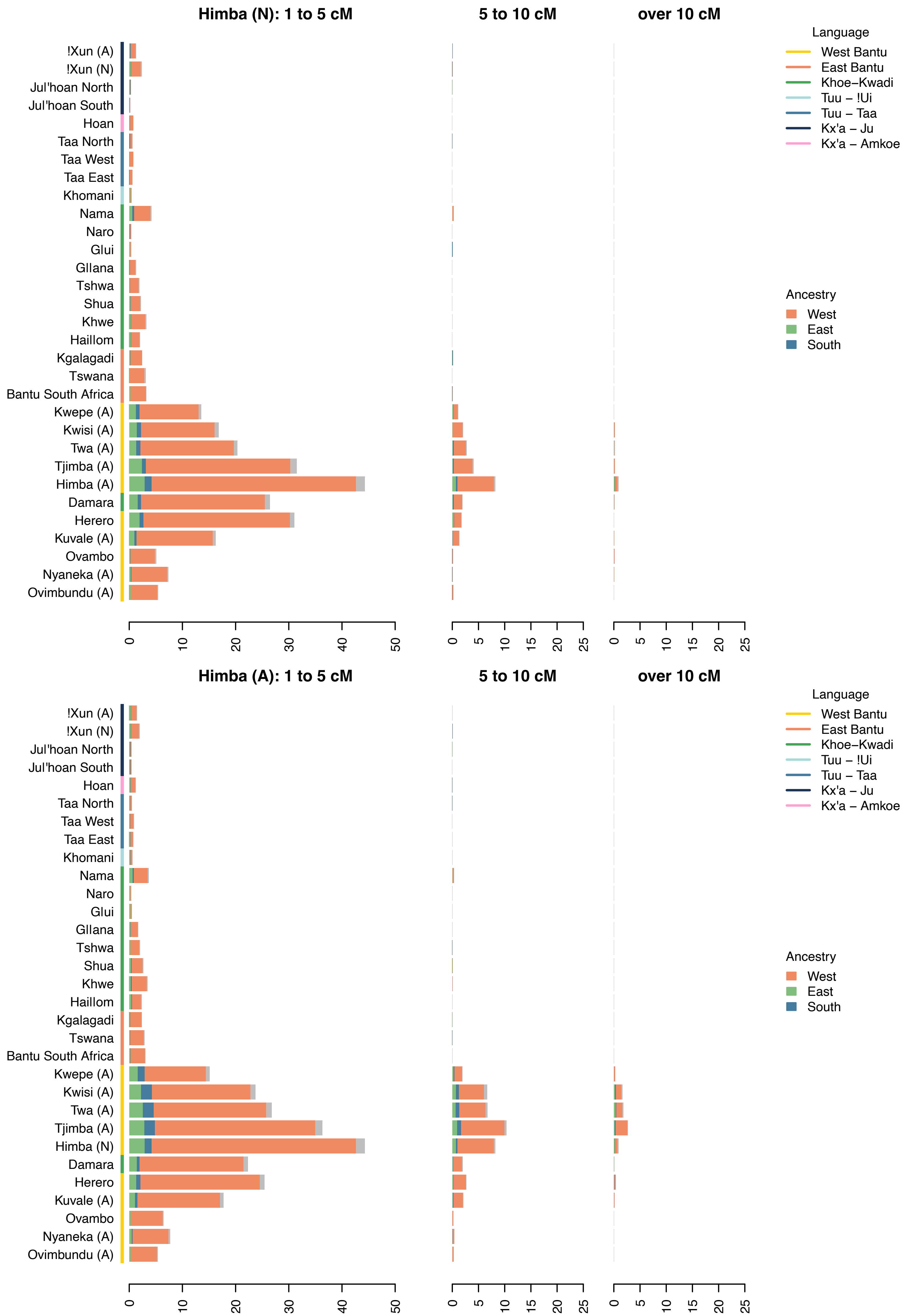

Himba (N): 1 to 5 cM
5 to 10 cM
over 10 cM
!Xun (A)
!Xun (N)
Jul'hoan North
Jul'hoan South
Hoan
Taa North
Taa West
Taa East
Khomani
Nama
Naro
Glui
Gllana
Tshwa
Shua
Khwe
Haillom
Kgalagadi
Tswana
Bantu South Africa
Kwepe (A)
Kwisi (A)
Twa (A)
Tjimba (A)
Himba (A)
Damara
Herero
Kuvale (A)
Ovambo
Nyaneka (A)
Ovimbundu (A)
0 10 20 30 40 50
0 5 10 15 20 25
0 5 10 15 20 25
Language
West Bantu
East Bantu
Khoe–Kwadi
Tuu – !Ui
Tuu – Taa
Kx'a – Ju
Kx'a – Amkoe
Ancestry
West
East
South
Himba (A): 1 to 5 cM
5 to 10 cM
over 10 cM
!Xun (A)
!Xun (N)
Jul'hoan North
Jul'hoan South
Hoan
Taa North
Taa West
Taa East
Khomani
Nama
Naro
Glui
Gllana
Tshwa
Shua
Khwe
Haillom
Kgalagadi
Tswana
Bantu South Africa
Kwepe (A)
Kwisi (A)
Twa (A)
Tjimba (A)
Himba (N)
Damara
Herero
Kuvale (A)
Ovambo
Nyaneka (A)
Ovimbundu (A)
0 10 20 30 40 50
0 5 10 15 20 25
0 5 10 15 20 25
Language
West Bantu
East Bantu
Khoe–Kwadi
Tuu – !Ui
Tuu – Taa
Kx'a – Ju
Kx'a – Amkoe
Ancestry
West
East
South

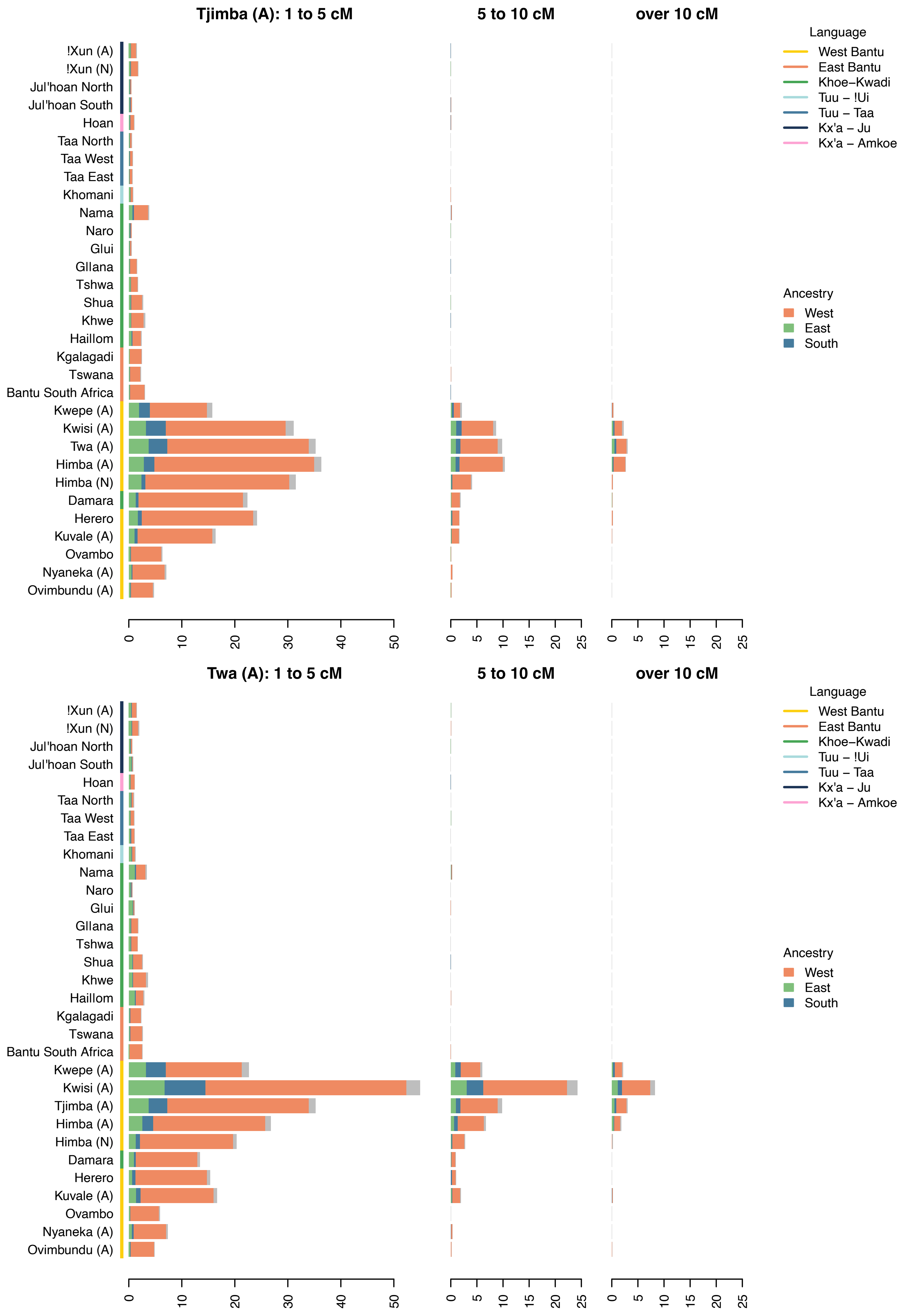

Tjimba (A): 1 to 5 cM
5 to 10 cM
over 10 cM
Language
West Bantu
East Bantu
Khoe–Kwadi
Tuu – !Ui
Tuu – Taa
Kx'a – Ju
Kx'a – Amkoe
Ancestry
West
East
South
!Xun (A)
!Xun (N)
Jul'hoan North
Jul'hoan South
Hoan
Taa North
Taa West
Taa East
Khomani
Nama
Naro
Glui
Gllana
Tshwa
Shua
Khwe
Haillom
Kgalagadi
Tswana
Bantu South Africa
Kwepe (A)
Kwisi (A)
Twa (A)
Himba (A)
Himba (N)
Damara
Herero
Kuvale (A)
Ovambo
Nyaneka (A)
Ovimbundu (A)
0
10
20
30
40
50
0
5
10
15
20
25
Twa (A): 1 to 5 cM
5 to 10 cM
over 10 cM
Kwepe (A)
Kwisi (A)
Tjimba (A)
Himba (A)
Himba (N)

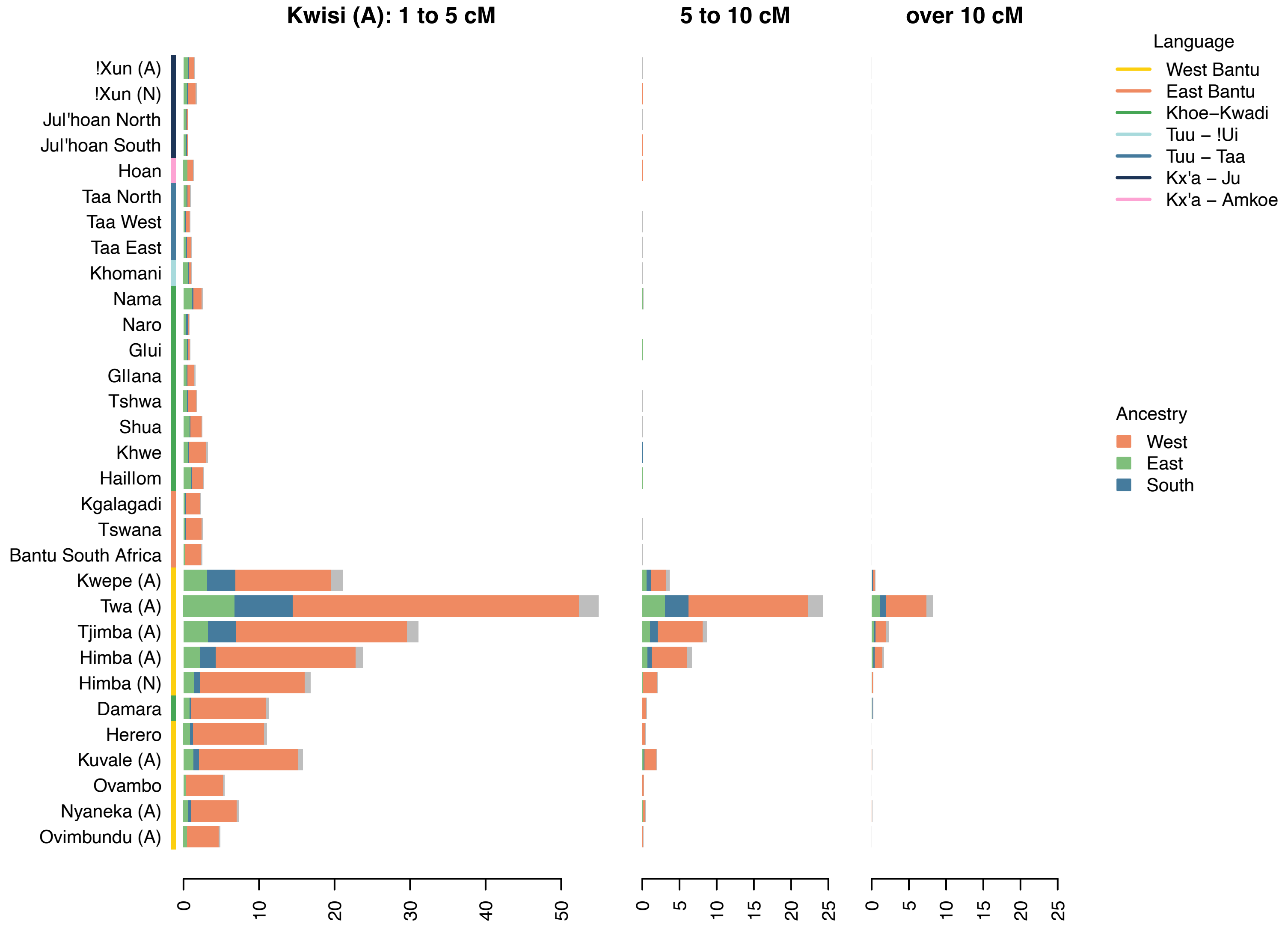
Kwisi (A): 1 to 5 cM
5 to 10 cM
over 10 cM
!Xun (A)
!Xun (N)
Jul'hoan North
Jul'hoan South
Hoan
Taa North
Taa West
Taa East
Khomani
Nama
Naro
Glui
Gllana
Tshwa
Shua
Khwe
Haillom
Kgalagadi
Tswana
Bantu South Africa
Kwepe (A)
Twa (A)
Tjimba (A)
Himba (A)
Himba (N)
Damara
Herero
Kuvale (A)
Ovambo
Nyaneka (A)
Ovimbundu (A)
0 10 20 30 40 50
0 5 10 15 20 25
0 5 10 15 20 25
Language
West Bantu
East Bantu
Khoe−Kwadi
Tuu − !Ui
Tuu − Taa
Kx'a − Ju
Kx'a − Amkoe
Ancestry
West
East
South

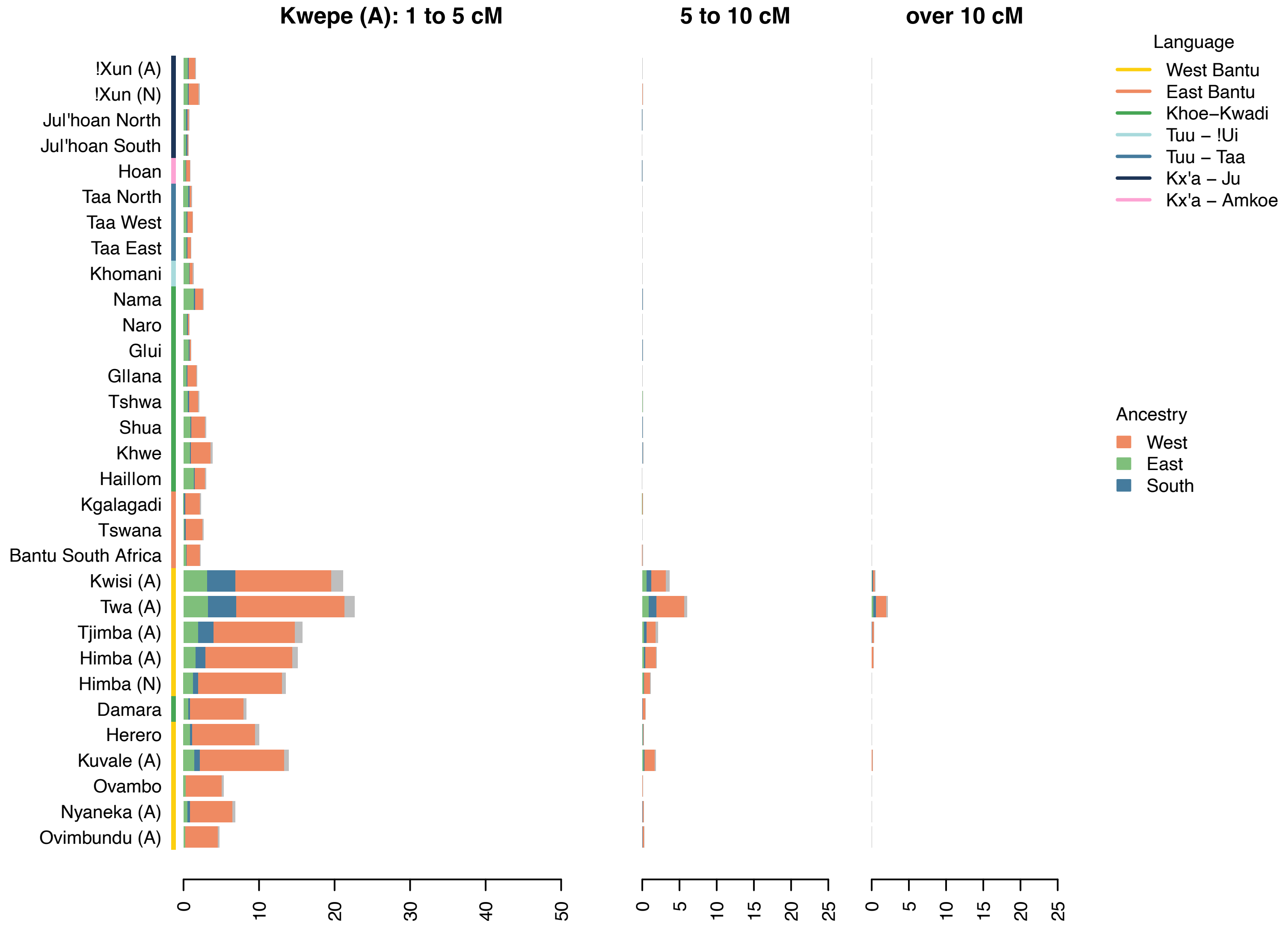
Kwepe (A): 1 to 5 cM
5 to 10 cM
over 10 cM
!Xun (A)
!Xun (N)
Jul'hoan North
Jul'hoan South
Hoan
Taa North
Taa West
Taa East
Khomani
Nama
Naro
Glui
Gllana
Tshwa
Shua
Khwe
Haillom
Kgalagadi
Tswana
Bantu South Africa
Kwisi (A)
Twa (A)
Tjimba (A)
Himba (A)
Himba (N)
Damara
Herero
Kuvale (A)
Ovambo
Nyaneka (A)
Ovimbundu (A)
0 10 20 30 40 50
0 5 10 15 20 25
0 5 10 15 20 25
Language
West Bantu
East Bantu
Khoe−Kwadi
Tuu − !Ui
Tuu − Taa
Kx'a − Ju
Kx'a − Amkoe
Ancestry
West
East
South

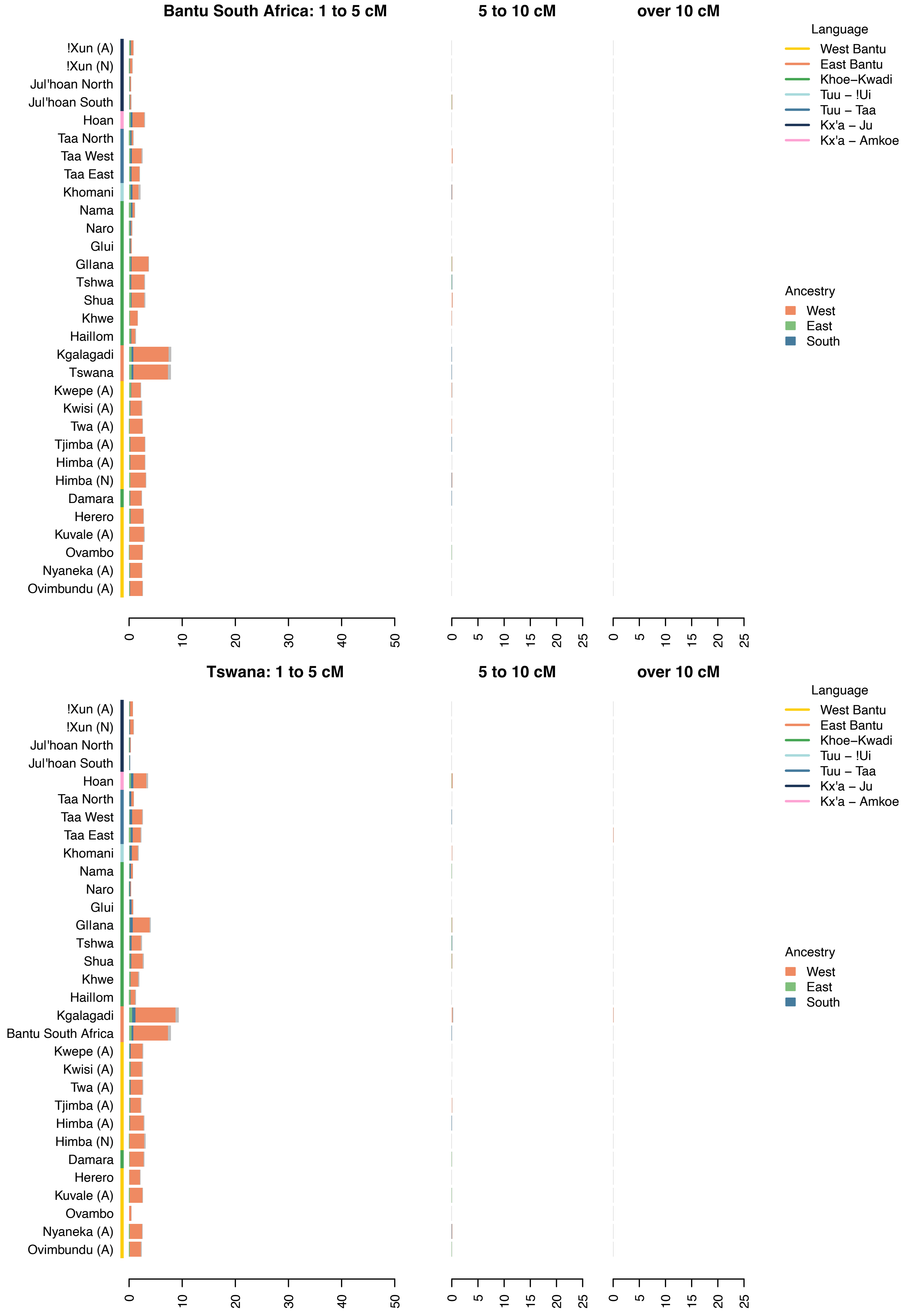
Bantu South Africa: 1 to 5 cM
5 to 10 cM
over 10 cM
Language
West Bantu
East Bantu
Khoe–Kwadi
Tuu – !Ui
Tuu – Taa
Kx'a – Ju
Kx'a – Amkoe
Ancestry
West
East
South
!Xun (A)
!Xun (N)
Jul'hoan North
Jul'hoan South
Hoan
Taa North
Taa West
Taa East
Khomani
Nama
Naro
Glui
Gllana
Tshwa
Shua
Khwe
Haillom
Kgalagadi
Tswana
Kwepe (A)
Kwisi (A)
Twa (A)
Tjimba (A)
Himba (A)
Himba (N)
Damara
Herero
Kuvale (A)
Ovambo
Nyaneka (A)
Ovimbundu (A)
0
10
20
30
40
50
0
5
10
15
20
25
Tswana: 1 to 5 cM
Bantu South Africa

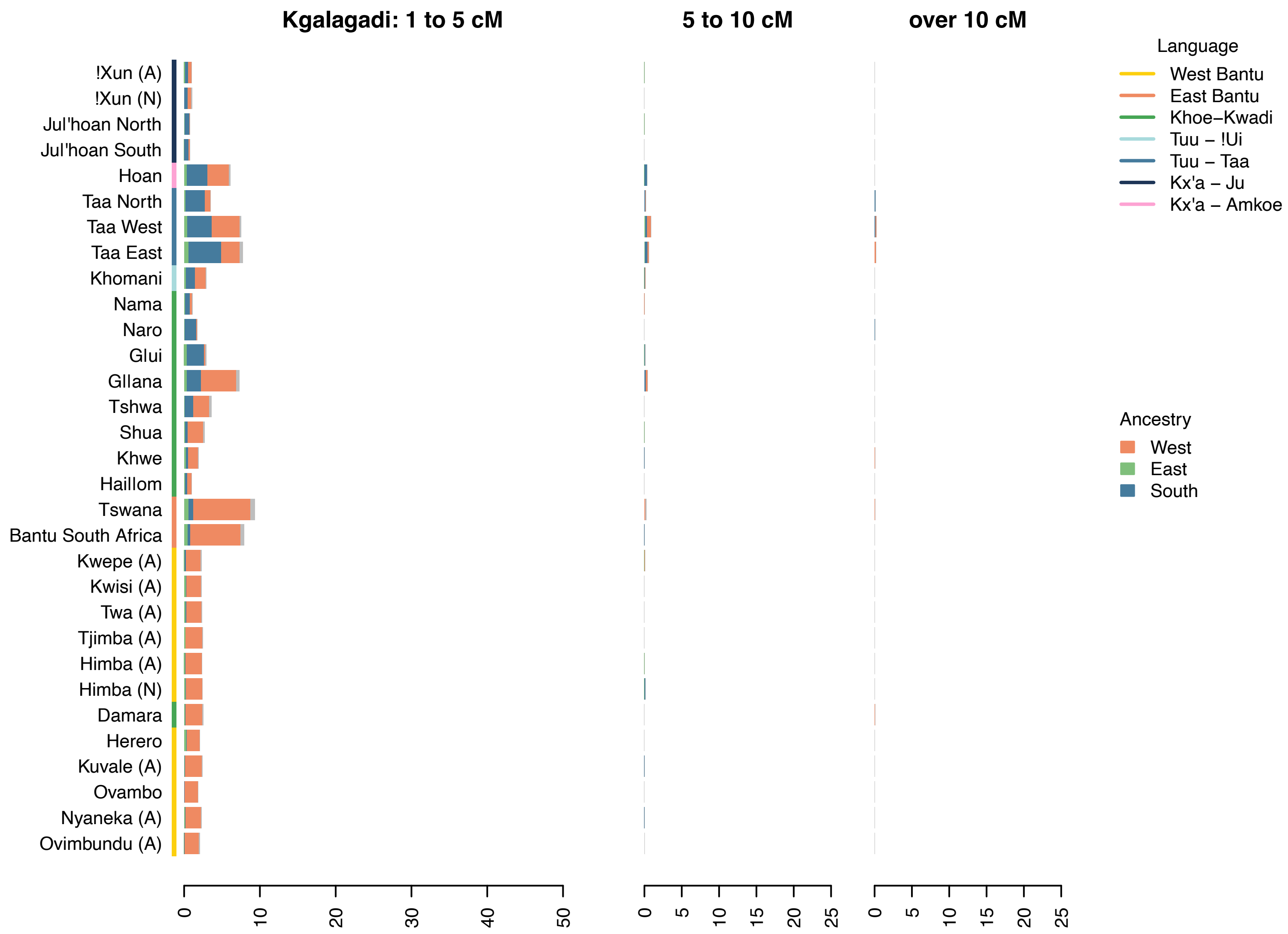

Kgalagadi: 1 to 5 cM
5 to 10 cM
over 10 cM
!Xun (A)
!Xun (N)
Jul'hoan North
Jul'hoan South
Hoan
Taa North
Taa West
Taa East
Khomani
Nama
Naro
Glui
Gllana
Tshwa
Shua
Khwe
Haillom
Tswana
Bantu South Africa
Kwepe (A)
Kwisi (A)
Twa (A)
Tjimba (A)
Himba (A)
Himba (N)
Damara
Herero
Kuvale (A)
Ovambo
Nyaneka (A)
Ovimbundu (A)
0 10 20 30 40 50
0 5 10 15 20 25
0 5 10 15 20 25
Language
West Bantu
East Bantu
Khoe–Kwadi
Tuu – !Ui
Tuu – Taa
Kx'a – Ju
Kx'a – Amkoe
Ancestry
West
East
South


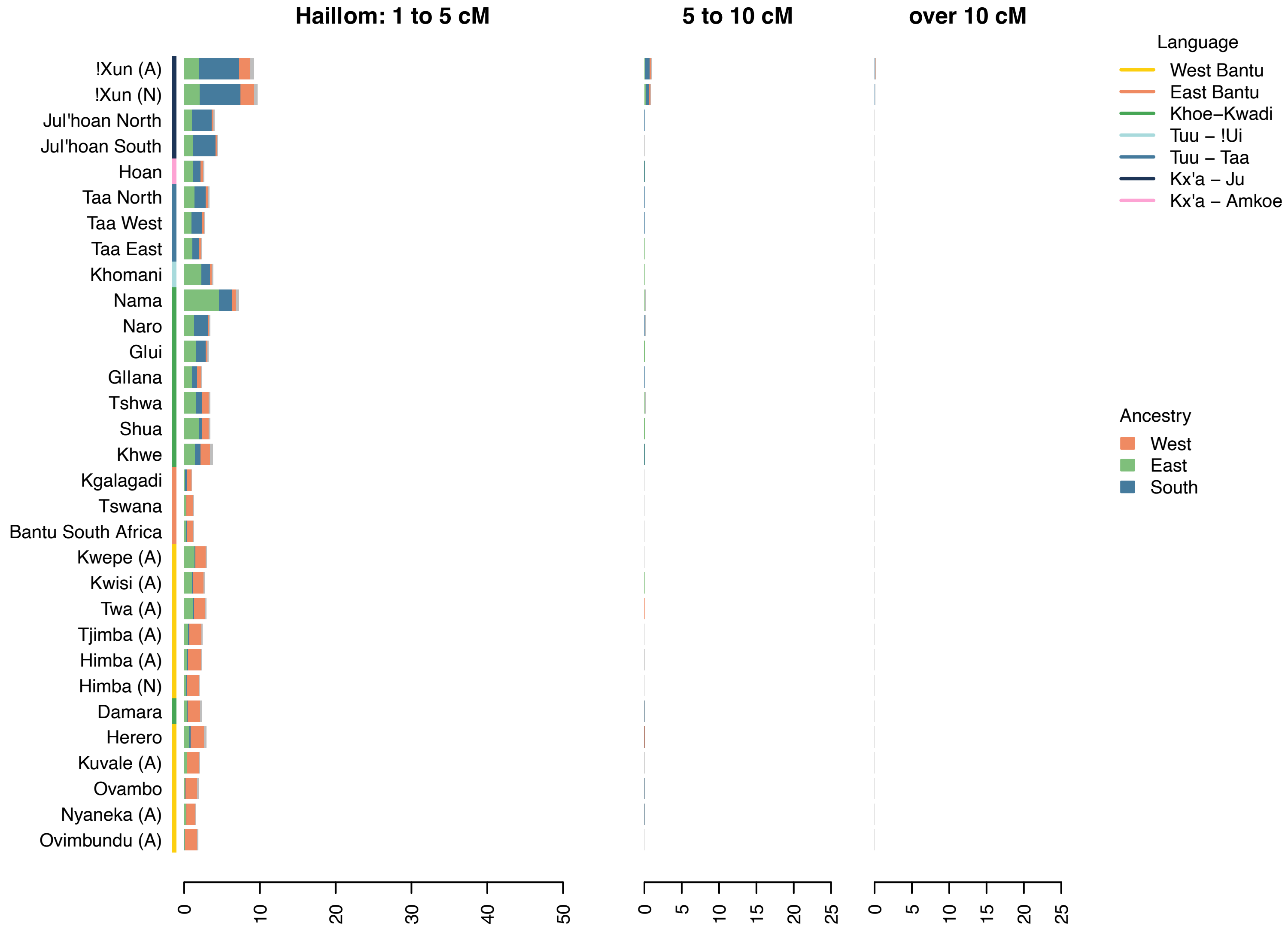

Haillom: 1 to 5 cM
5 to 10 cM
over 10 cM
!Xun (A)
!Xun (N)
Jul'hoan North
Jul'hoan South
Hoan
Taa North
Taa West
Taa East
Khomani
Nama
Naro
Glui
Gllana
Tshwa
Shua
Khwe
Kgalagadi
Tswana
Bantu South Africa
Kwepe (A)
Kwisi (A)
Twa (A)
Tjimba (A)
Himba (A)
Himba (N)
Damara
Herero
Kuvale (A)
Ovambo
Nyaneka (A)
Ovimbundu (A)
0 10 20 30 40 50
0 5 10 15 20 25
0 5 10 15 20 25
Language
West Bantu
East Bantu
Khoe–Kwadi
Tuu – !Ui
Tuu – Taa
Kx'a – Ju
Kx'a – Amkoe
Ancestry
West
East
South

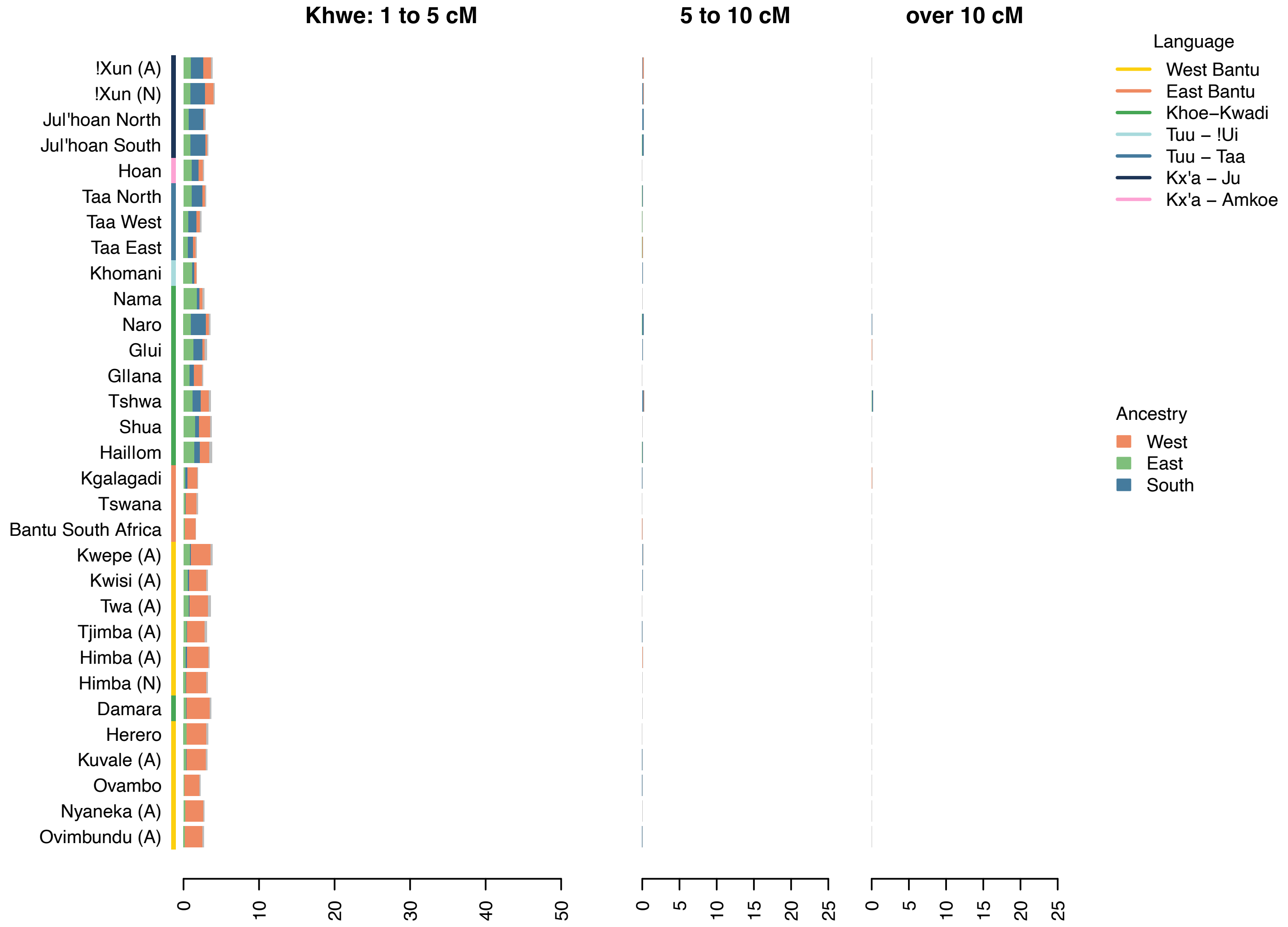
Khwe: 1 to 5 cM
5 to 10 cM
over 10 cM
!Xun (A)
!Xun (N)
Jul'hoan North
Jul'hoan South
Hoan
Taa North
Taa West
Taa East
Khomani
Nama
Naro
Glui
Gllana
Tshwa
Shua
Haillom
Kgalagadi
Tswana
Bantu South Africa
Kwepe (A)
Kwisi (A)
Twa (A)
Tjimba (A)
Himba (A)
Himba (N)
Damara
Herero
Kuvale (A)
Ovambo
Nyaneka (A)
Ovimbundu (A)
Language
West Bantu
East Bantu
Khoe−Kwadi
Tuu − !Ui
Tuu − Taa
Kx'a − Ju
Kx'a − Amkoe
Ancestry
West
East
South
0
10
20
30
40
50
0
5
10
15
20
25
0
5
10
15
20
25

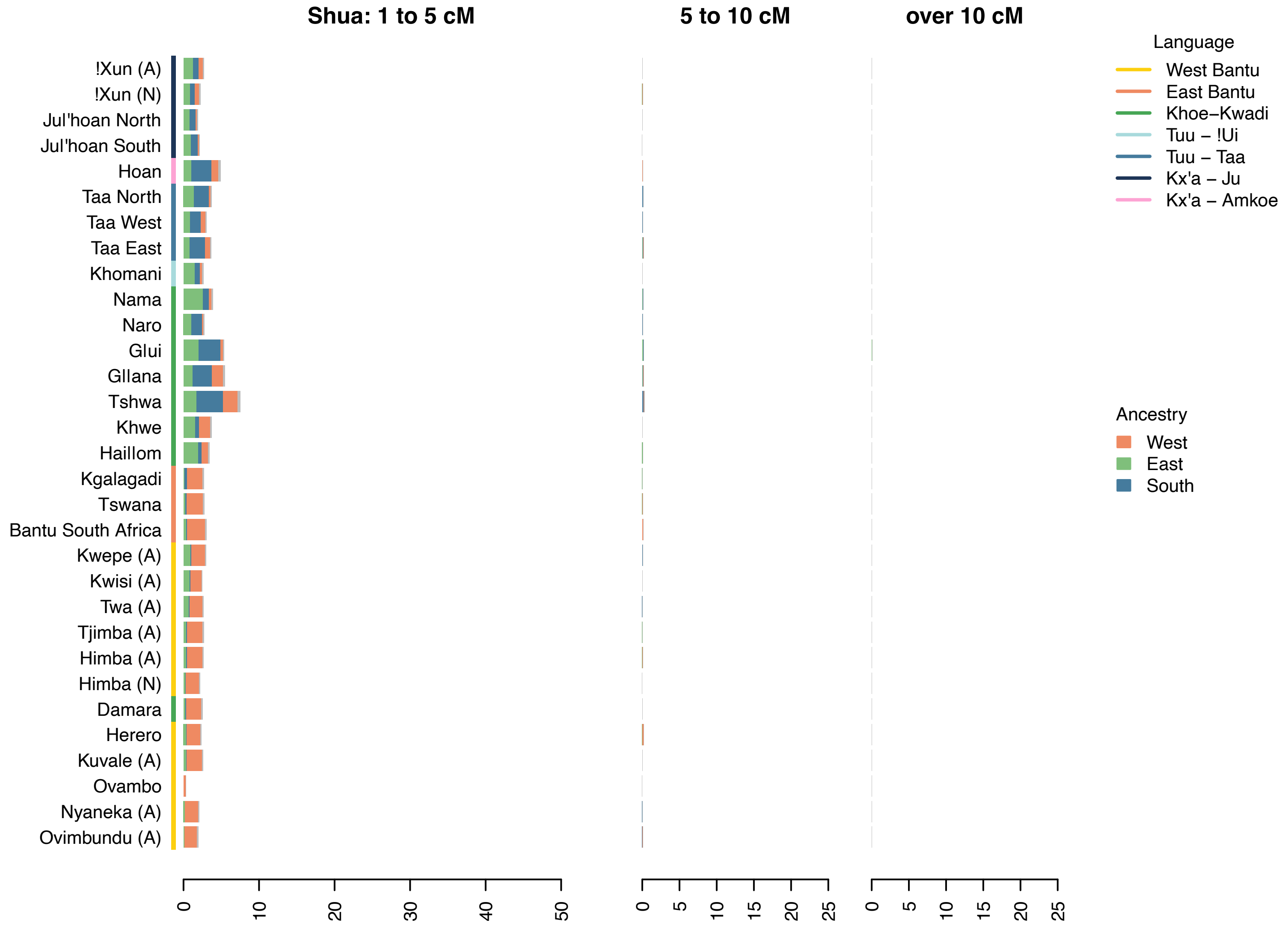
Shua: 1 to 5 cM
5 to 10 cM
over 10 cM
!Xun (A)
!Xun (N)
Jul'hoan North
Jul'hoan South
Hoan
Taa North
Taa West
Taa East
Khomani
Nama
Naro
Glui
Gllana
Tshwa
Khwe
Haillom
Kgalagadi
Tswana
Bantu South Africa
Kwepe (A)
Kwisi (A)
Twa (A)
Tjimba (A)
Himba (A)
Himba (N)
Damara
Herero
Kuvale (A)
Ovambo
Nyaneka (A)
Ovimbundu (A)
Language
West Bantu
East Bantu
Khoe−Kwadi
Tuu − !Ui
Tuu − Taa
Kx'a − Ju
Kx'a − Amkoe
Ancestry
West
East
South
0
10
20
30
40
50
0
5
10
15
20
25
0
5
10
15
20
25

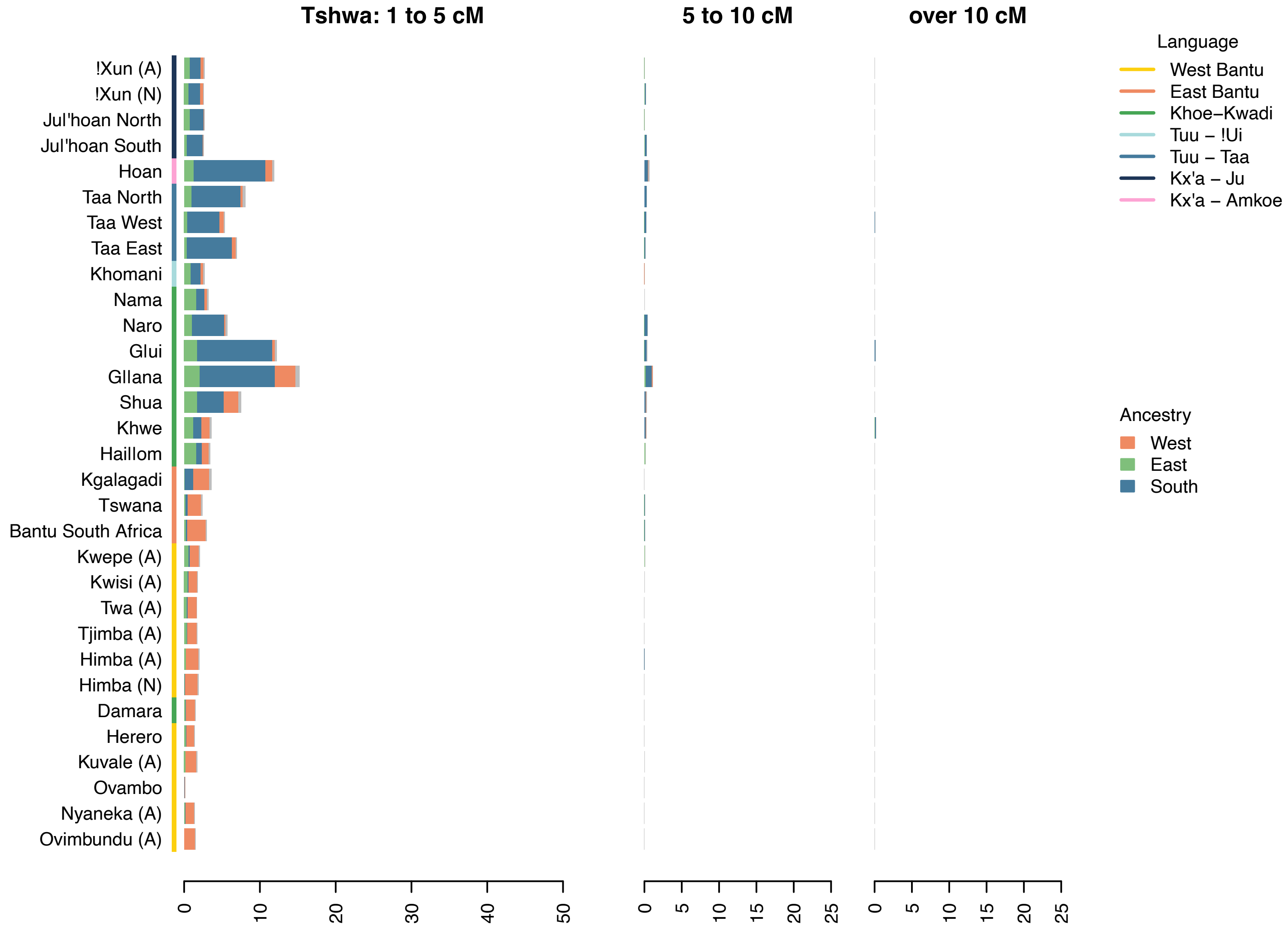

Tshwa: 1 to 5 cM
5 to 10 cM
over 10 cM
!Xun (A)
!Xun (N)
Jul'hoan North
Jul'hoan South
Hoan
Taa North
Taa West
Taa East
Khomani
Nama
Naro
Glui
Gllana
Shua
Khwe
Haillom
Kgalagadi
Tswana
Bantu South Africa
Kwepe (A)
Kwisi (A)
Twa (A)
Tjimba (A)
Himba (A)
Himba (N)
Damara
Herero
Kuvale (A)
Ovambo
Nyaneka (A)
Ovimbundu (A)
0 10 20 30 40 50
0 5 10 15 20 25
0 5 10 15 20 25
Language
West Bantu
East Bantu
Khoe–Kwadi
Tuu – !Ui
Tuu – Taa
Kx'a – Ju
Kx'a – Amkoe
Ancestry
West
East
South


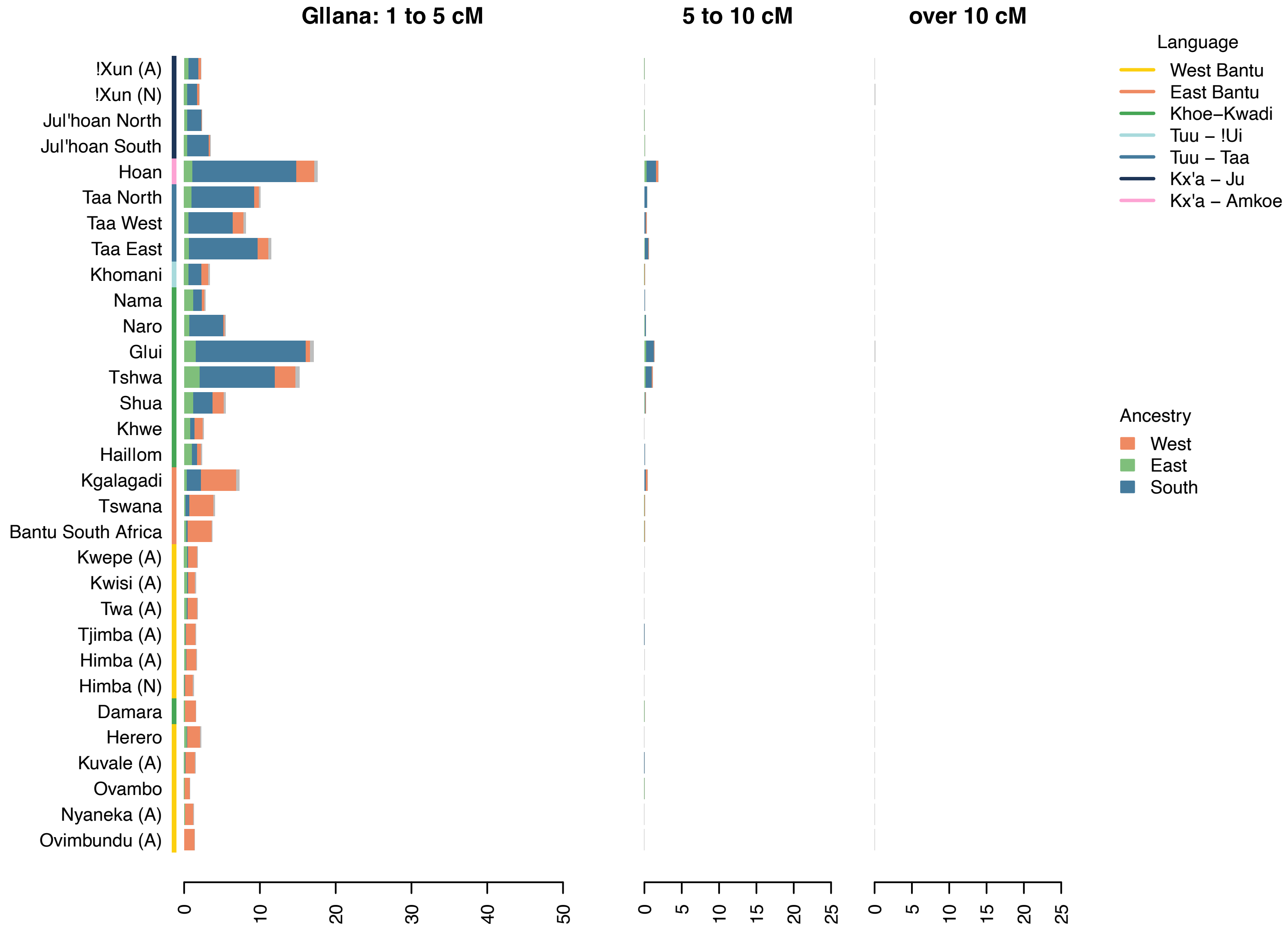

Gllana: 1 to 5 cM
5 to 10 cM
over 10 cM
!Xun (A)
!Xun (N)
Jul'hoan North
Jul'hoan South
Hoan
Taa North
Taa West
Taa East
Khomani
Nama
Naro
Glui
Tshwa
Shua
Khwe
Haillom
Kgalagadi
Tswana
Bantu South Africa
Kwepe (A)
Kwisi (A)
Twa (A)
Tjimba (A)
Himba (A)
Himba (N)
Damara
Herero
Kuvale (A)
Ovambo
Nyaneka (A)
Ovimbundu (A)
0 10 20 30 40 50
0 5 10 15 20 25
0 5 10 15 20 25
Language
West Bantu
East Bantu
Khoe–Kwadi
Tuu – !Ui
Tuu – Taa
Kx'a – Ju
Kx'a – Amkoe
Ancestry
West
East
South

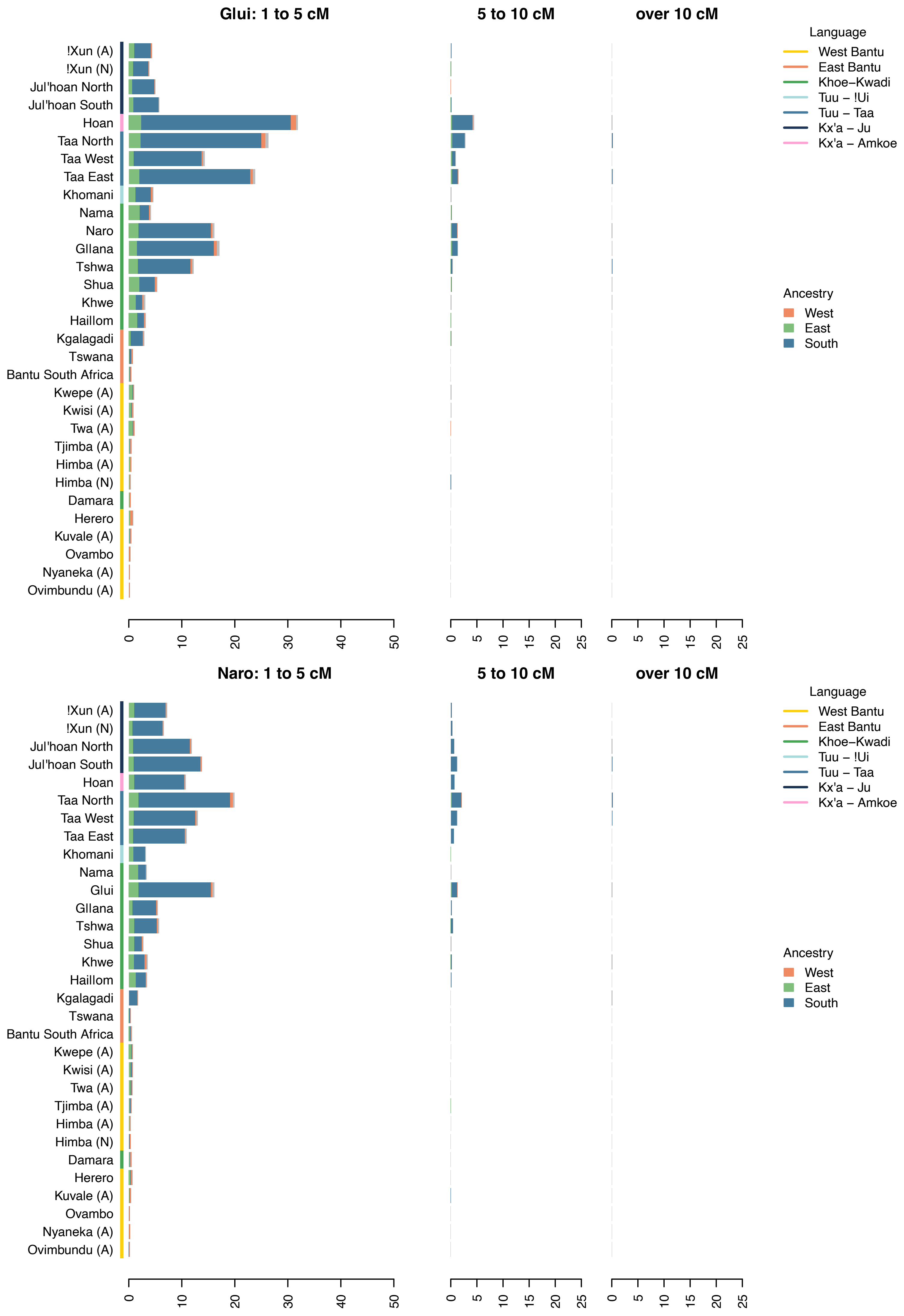
Glui: 1 to 5 cM
5 to 10 cM
over 10 cM
!Xun (A)
!Xun (N)
Jul'hoan North
Jul'hoan South
Hoan
Taa North
Taa West
Taa East
Khomani
Nama
Naro
Gllana
Tshwa
Shua
Khwe
Haillom
Kgalagadi
Tswana
Bantu South Africa
Kwepe (A)
Kwisi (A)
Twa (A)
Tjimba (A)
Himba (A)
Himba (N)
Damara
Herero
Kuvale (A)
Ovambo
Nyaneka (A)
Ovimbundu (A)
0
10
20
30
40
50
0
5
10
15
20
25
0
5
10
15
20
25
Language
West Bantu
East Bantu
Khoe–Kwadi
Tuu – !Ui
Tuu – Taa
Kx'a – Ju
Kx'a – Amkoe
Ancestry
West
East
South
Naro: 1 to 5 cM
5 to 10 cM
over 10 cM
!Xun (A)
!Xun (N)
Jul'hoan North
Jul'hoan South
Hoan
Taa North
Taa West
Taa East
Khomani
Nama
Glui
Gllana
Tshwa
Shua
Khwe
Haillom
Kgalagadi
Tswana
Bantu South Africa
Kwepe (A)
Kwisi (A)
Twa (A)
Tjimba (A)
Himba (A)
Himba (N)
Damara
Herero
Kuvale (A)
Ovambo
Nyaneka (A)
Ovimbundu (A)
0
10
20
30
40
50
0
5
10
15
20
25
0
5
10
15
20
25
Language
West Bantu
East Bantu
Khoe–Kwadi
Tuu – !Ui
Tuu – Taa
Kx'a – Ju
Kx'a – Amkoe
Ancestry
West
East
South

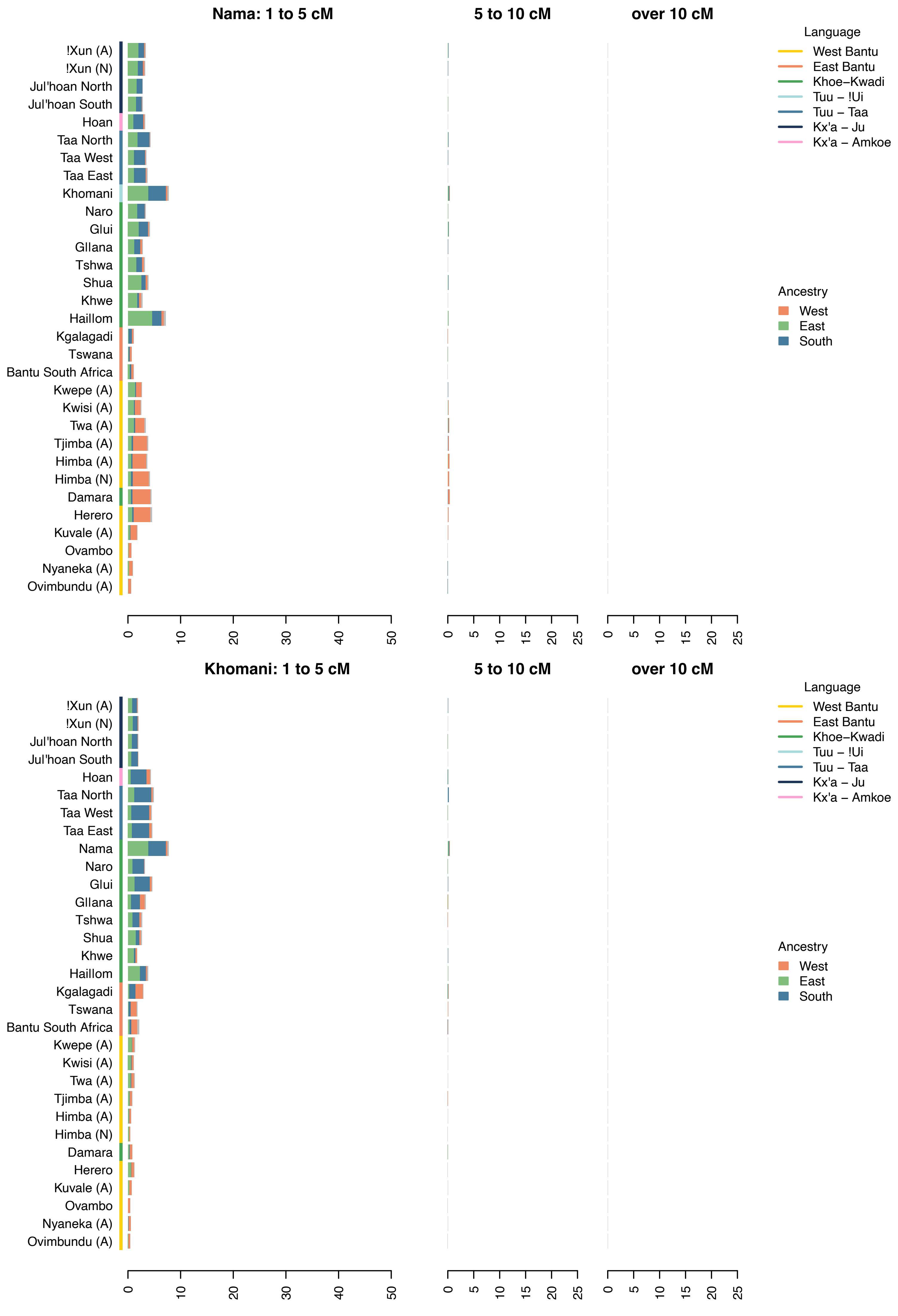
Nama: 1 to 5 cM
5 to 10 cM
over 10 cM
!Xun (A)
!Xun (N)
Jul'hoan North
Jul'hoan South
Hoan
Taa North
Taa West
Taa East
Khomani
Naro
Glui
Gllana
Tshwa
Shua
Khwe
Haillom
Kgalagadi
Tswana
Bantu South Africa
Kwepe (A)
Kwisi (A)
Twa (A)
Tjimba (A)
Himba (A)
Himba (N)
Damara
Herero
Kuvale (A)
Ovambo
Nyaneka (A)
Ovimbundu (A)
0
10
20
30
40
50
0
5
10
15
20
25
0
5
10
15
20
25
Language
West Bantu
East Bantu
Khoe–Kwadi
Tuu – !Ui
Tuu – Taa
Kx'a – Ju
Kx'a – Amkoe
Ancestry
West
East
South
Khomani: 1 to 5 cM
5 to 10 cM
over 10 cM
!Xun (A)
!Xun (N)
Jul'hoan North
Jul'hoan South
Hoan
Taa North
Taa West
Taa East
Nama
Naro
Glui
Gllana
Tshwa
Shua
Khwe
Haillom
Kgalagadi
Tswana
Bantu South Africa
Kwepe (A)
Kwisi (A)
Twa (A)
Tjimba (A)
Himba (A)
Himba (N)
Damara
Herero
Kuvale (A)
Ovambo
Nyaneka (A)
Ovimbundu (A)
0
10
20
30
40
50
0
5
10
15
20
25
0
5
10
15
20
25
Language
West Bantu
East Bantu
Khoe–Kwadi
Tuu – !Ui
Tuu – Taa
Kx'a – Ju
Kx'a – Amkoe
Ancestry
West
East
South

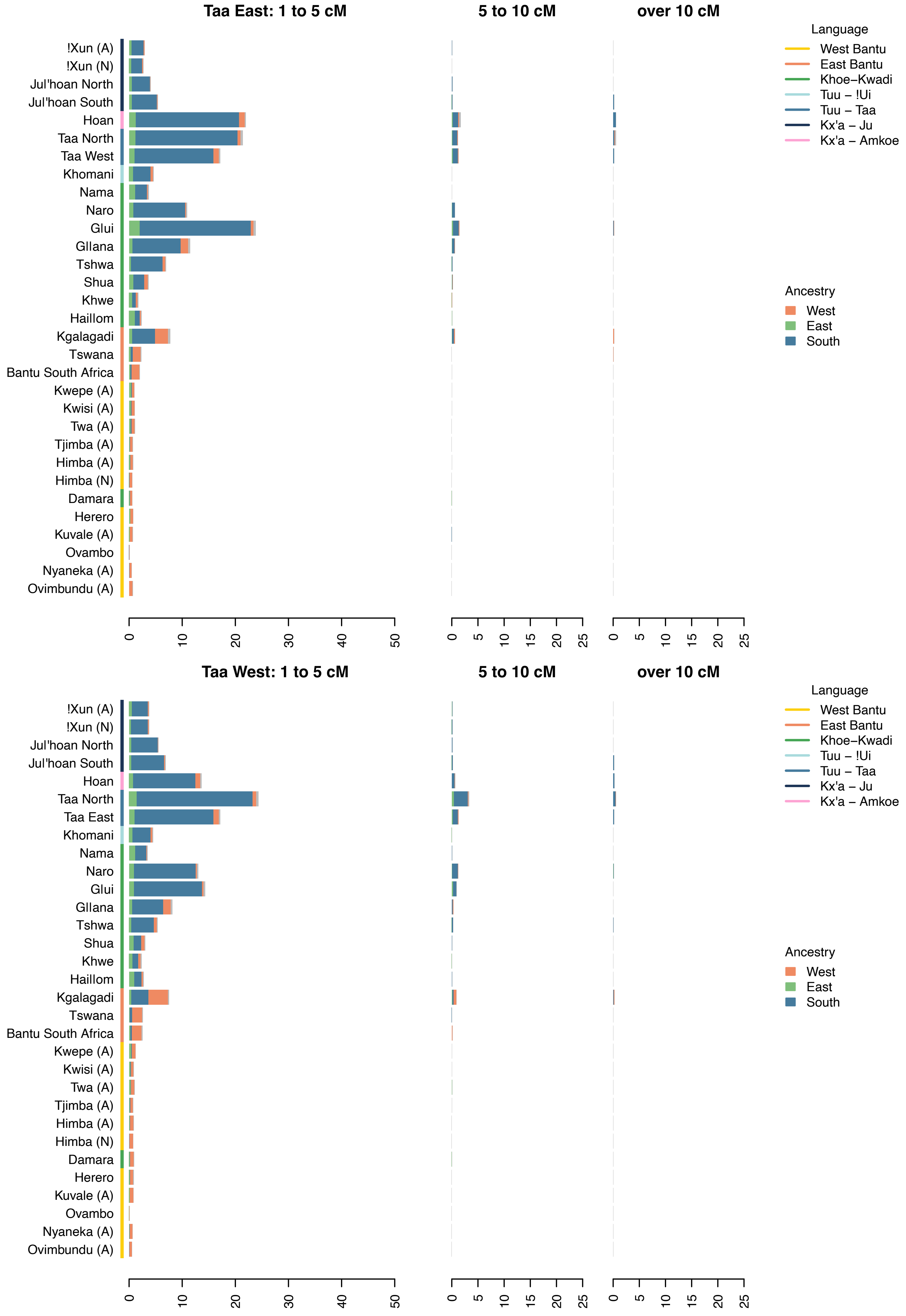

Taa East: 1 to 5 cM
5 to 10 cM
over 10 cM
!Xun (A)
!Xun (N)
Jul'hoan North
Jul'hoan South
Hoan
Taa North
Taa West
Khomani
Nama
Naro
Glui
GIIana
Tshwa
Shua
Khwe
Haillom
Kgalagadi
Tswana
Bantu South Africa
Kwepe (A)
Kwisi (A)
Twa (A)
Tjimba (A)
Himba (A)
Himba (N)
Damara
Herero
Kuvale (A)
Ovambo
Nyaneka (A)
Ovimbundu (A)
0 10 20 30 40 50
0 5 10 15 20 25
0 5 10 15 20 25
Language
West Bantu
East Bantu
Khoe−Kwadi
Tuu − !Ui
Tuu − Taa
Kx'a − Ju
Kx'a − Amkoe
Ancestry
West
East
South
Taa West: 1 to 5 cM
5 to 10 cM
over 10 cM
!Xun (A)
!Xun (N)
Jul'hoan North
Jul'hoan South
Hoan
Taa North
Taa East
Khomani
Nama
Naro
Glui
GIIana
Tshwa
Shua
Khwe
Haillom
Kgalagadi
Tswana
Bantu South Africa
Kwepe (A)
Kwisi (A)
Twa (A)
Tjimba (A)
Himba (A)
Himba (N)
Damara
Herero
Kuvale (A)
Ovambo
Nyaneka (A)
Ovimbundu (A)
0 10 20 30 40 50
0 5 10 15 20 25
0 5 10 15 20 25
Language
West Bantu
East Bantu
Khoe−Kwadi
Tuu − !Ui
Tuu − Taa
Kx'a − Ju
Kx'a − Amkoe
Ancestry
West
East
South

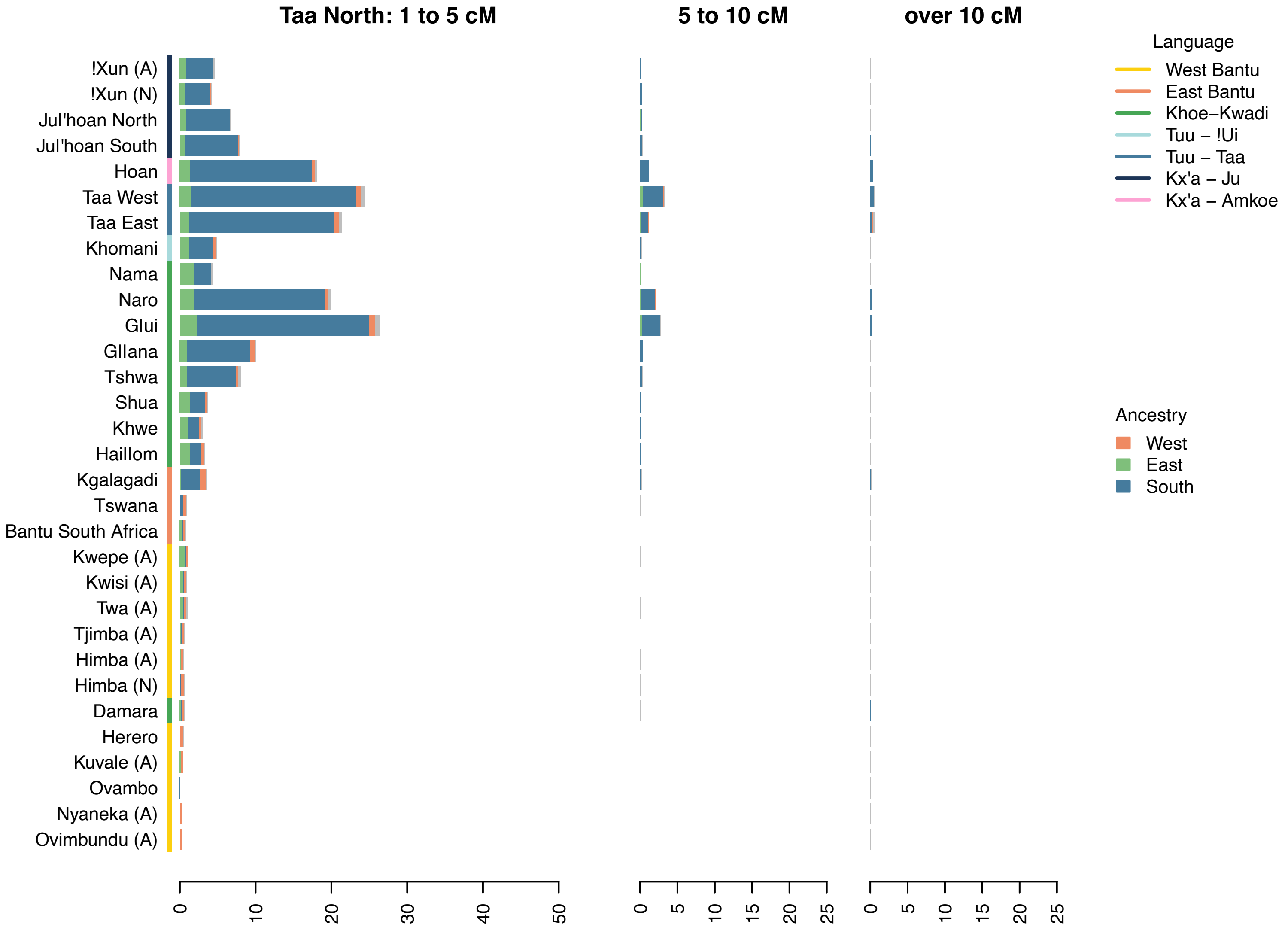
Taa North: 1 to 5 cM
5 to 10 cM
over 10 cM
!Xun (A)
!Xun (N)
Jul'hoan North
Jul'hoan South
Hoan
Taa West
Taa East
Khomani
Nama
Naro
Glui
Gllana
Tshwa
Shua
Khwe
Haillom
Kgalagadi
Tswana
Bantu South Africa
Kwepe (A)
Kwisi (A)
Twa (A)
Tjimba (A)
Himba (A)
Himba (N)
Damara
Herero
Kuvale (A)
Ovambo
Nyaneka (A)
Ovimbundu (A)
0 10 20 30 40 50
0 5 10 15 20 25
0 5 10 15 20 25
Language
West Bantu
East Bantu
Khoe–Kwadi
Tuu – !Ui
Tuu – Taa
Kx'a – Ju
Kx'a – Amkoe
Ancestry
West
East
South

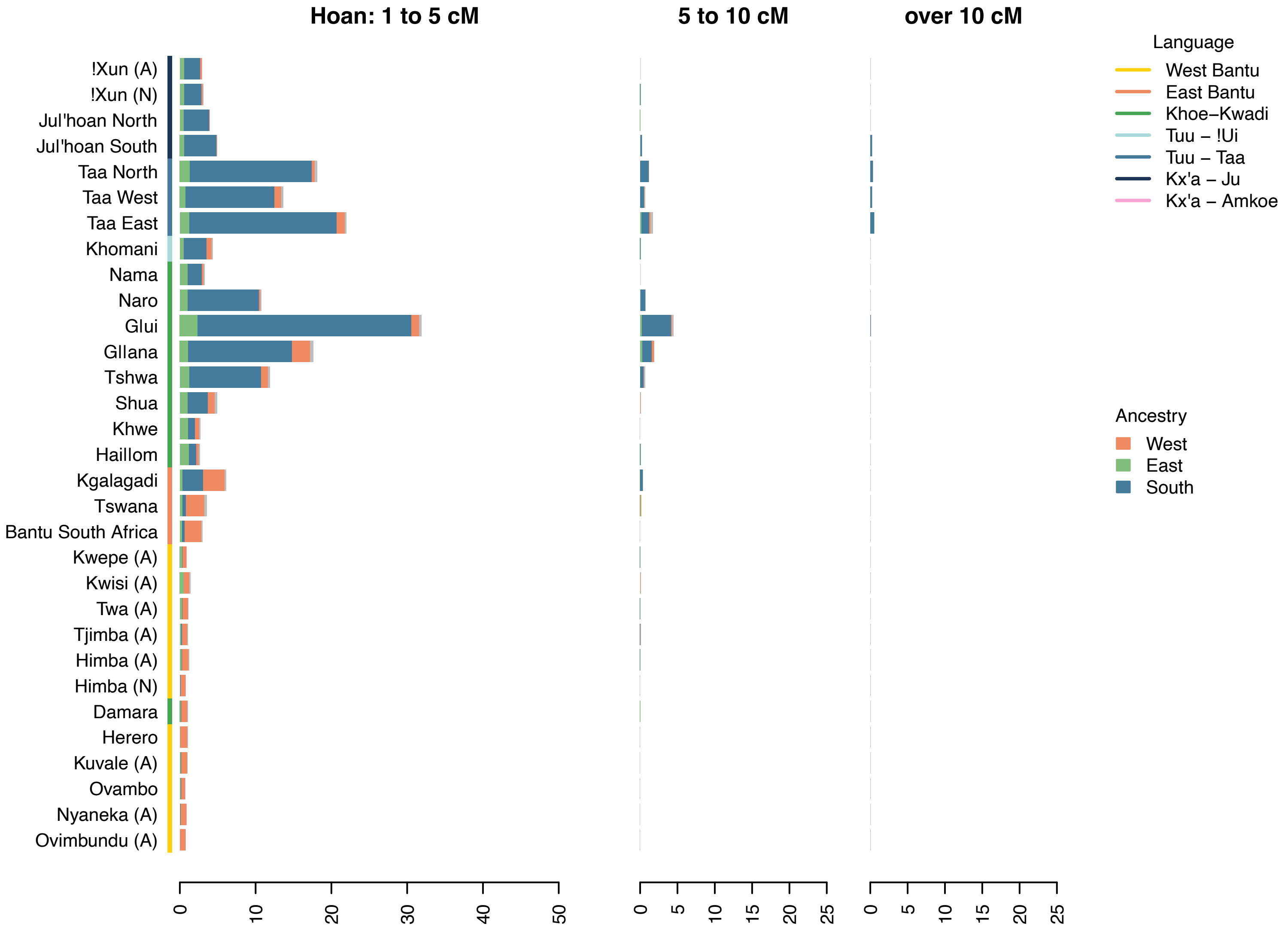
Hoan: 1 to 5 cM
5 to 10 cM
over 10 cM
!Xun (A)
!Xun (N)
Jul'hoan North
Jul'hoan South
Taa North
Taa West
Taa East
Khomani
Nama
Naro
Glui
Gllana
Tshwa
Shua
Khwe
Haillom
Kgalagadi
Tswana
Bantu South Africa
Kwepe (A)
Kwisi (A)
Twa (A)
Tjimba (A)
Himba (A)
Himba (N)
Damara
Herero
Kuvale (A)
Ovambo
Nyaneka (A)
Ovimbundu (A)
0 10 20 30 40 50
0 5 10 15 20 25
0 5 10 15 20 25
Language
West Bantu
East Bantu
Khoe–Kwadi
Tuu – !Ui
Tuu – Taa
Kx'a – Ju
Kx'a – Amkoe
Ancestry
West
East
South

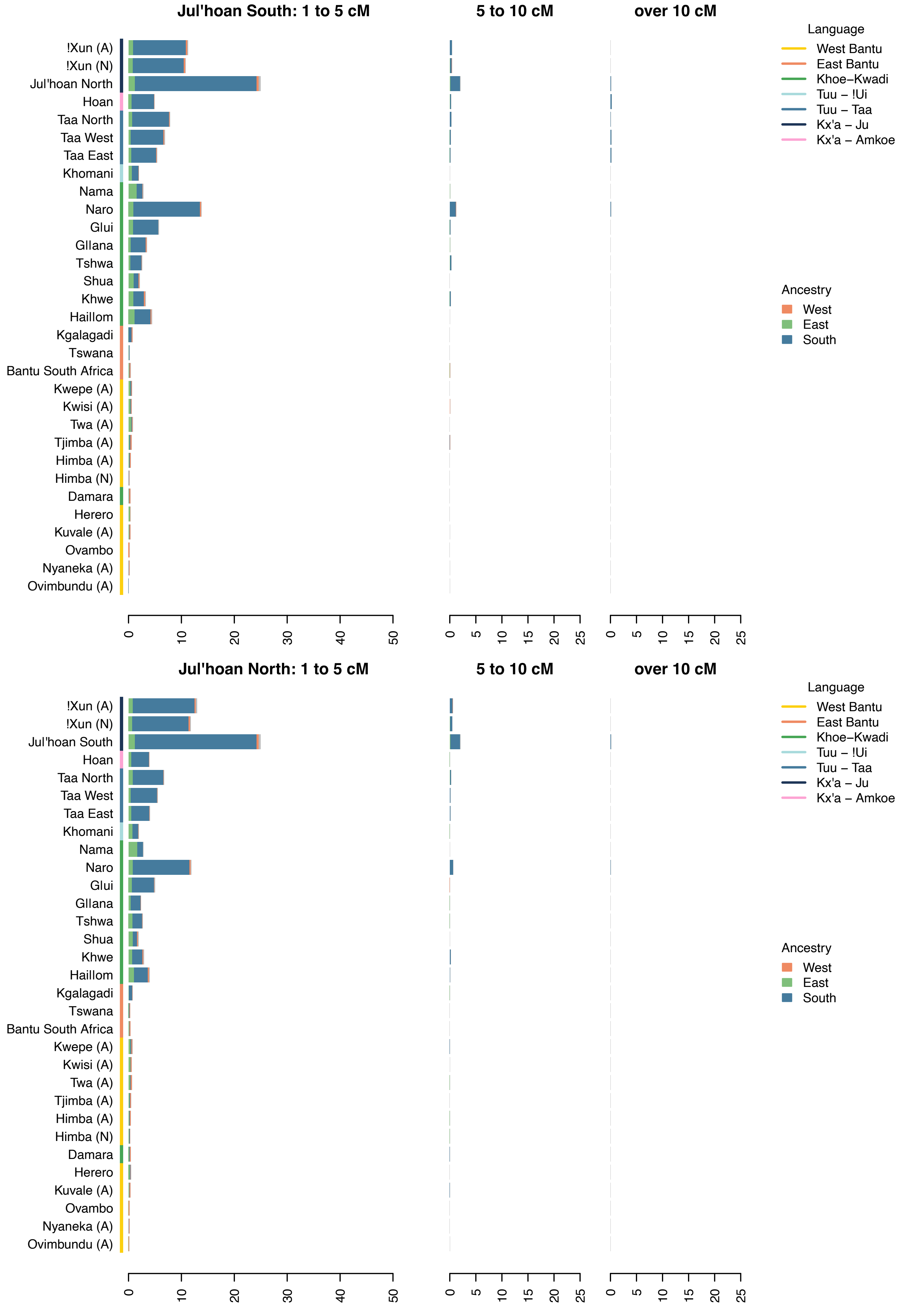
Jul'hoan South: 1 to 5 cM
5 to 10 cM
over 10 cM
Language
West Bantu
East Bantu
Khoe–Kwadi
Tuu – !Ui
Tuu – Taa
Kx'a – Ju
Kx'a – Amkoe
Ancestry
West
East
South
!Xun (A)
!Xun (N)
Jul'hoan North
Hoan
Taa North
Taa West
Taa East
Khomani
Nama
Naro
Glui
Gllana
Tshwa
Shua
Khwe
Haillom
Kgalagadi
Tswana
Bantu South Africa
Kwepe (A)
Kwisi (A)
Twa (A)
Tjimba (A)
Himba (A)
Himba (N)
Damara
Herero
Kuvale (A)
Ovambo
Nyaneka (A)
Ovimbundu (A)
0
10
20
30
40
50
0
5
10
15
20
25
Jul'hoan North: 1 to 5 cM
5 to 10 cM
over 10 cM
Jul'hoan South

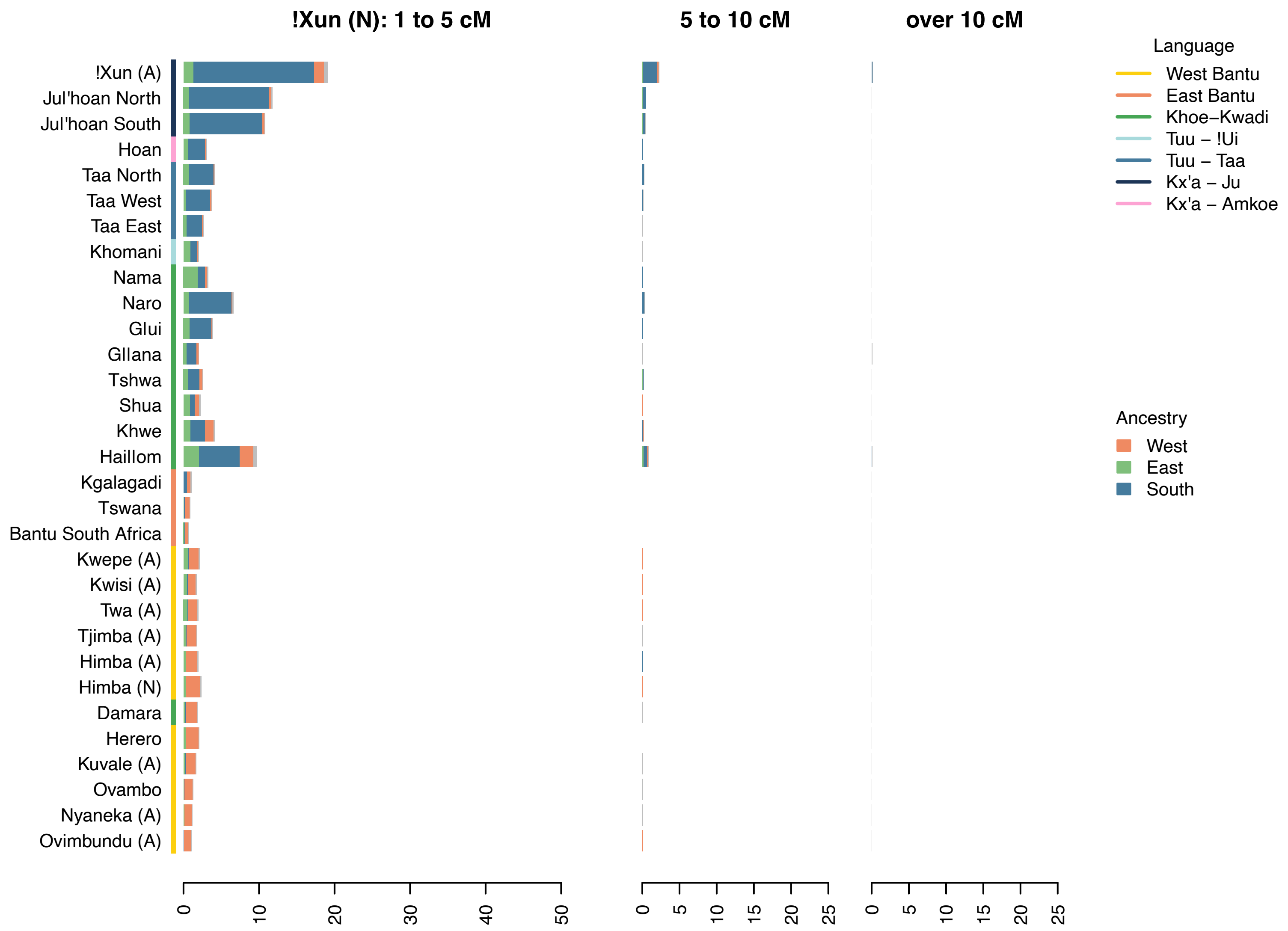
!Xun (N): 1 to 5 cM
5 to 10 cM
over 10 cM
!Xun (A)
Jul'hoan North
Jul'hoan South
Hoan
Taa North
Taa West
Taa East
Khomani
Nama
Naro
Glui
Gllana
Tshwa
Shua
Khwe
Haillom
Kgalagadi
Tswana
Bantu South Africa
Kwepe (A)
Kwisi (A)
Twa (A)
Tjimba (A)
Himba (A)
Himba (N)
Damara
Herero
Kuvale (A)
Ovambo
Nyaneka (A)
Ovimbundu (A)
Language
West Bantu
East Bantu
Khoe–Kwadi
Tuu – !Ui
Tuu – Taa
Kx'a – Ju
Kx'a – Amkoe
Ancestry
West
East
South
0 10 20 30 40 50
0 5 10 15 20 25
0 5 10 15 20 25

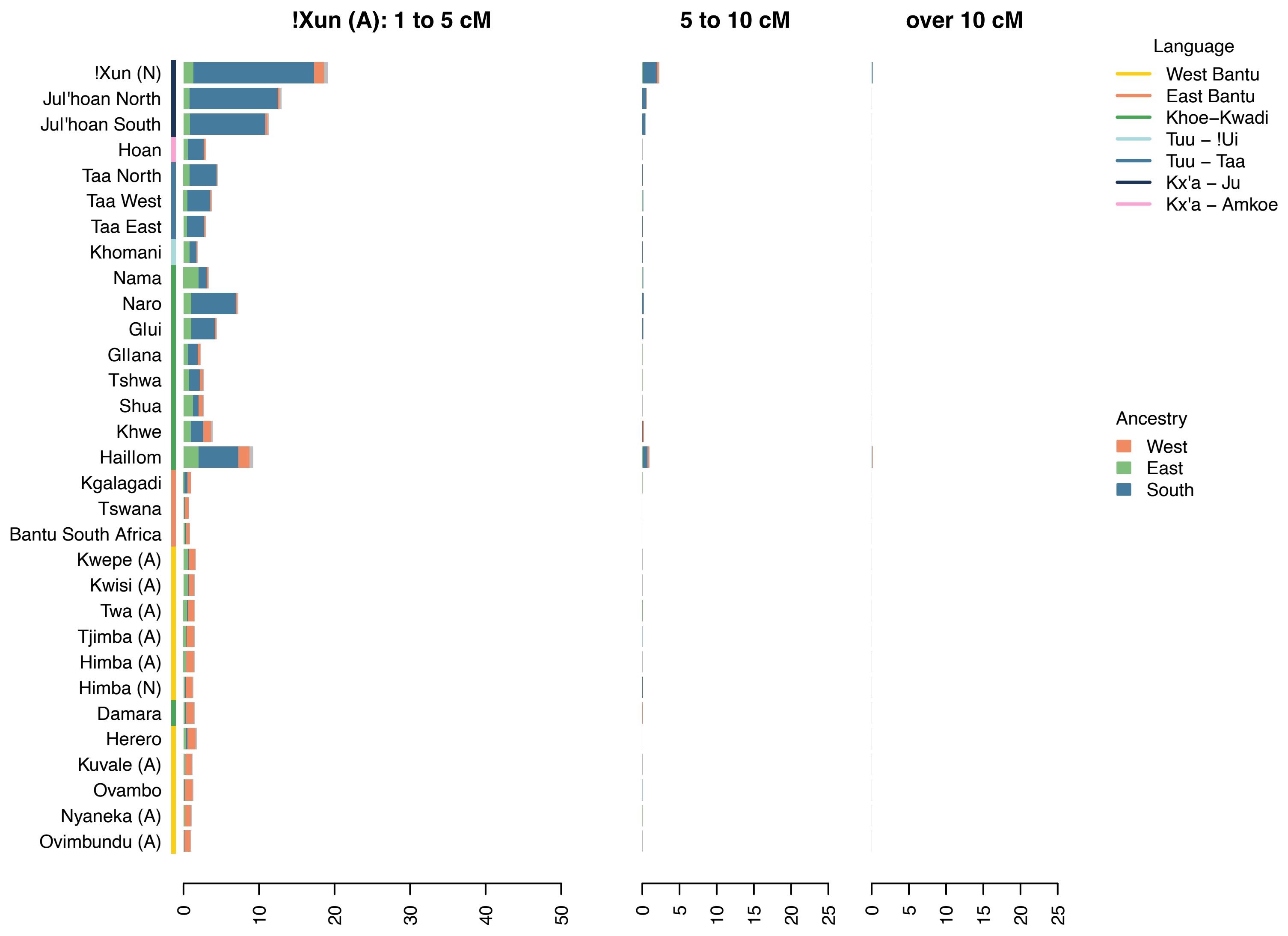
!Xun (A): 1 to 5 cM
5 to 10 cM
over 10 cM
!Xun (N)
Jul'hoan North
Jul'hoan South
Hoan
Taa North
Taa West
Taa East
Khomani
Nama
Naro
Glui
Gllana
Tshwa
Shua
Khwe
Haillom
Kgalagadi
Tswana
Bantu South Africa
Kwepe (A)
Kwisi (A)
Twa (A)
Tjimba (A)
Himba (A)
Himba (N)
Damara
Herero
Kuvale (A)
Ovambo
Nyaneka (A)
Ovimbundu (A)
Language
West Bantu
East Bantu
Khoe–Kwadi
Tuu – !Ui
Tuu – Taa
Kx'a – Ju
Kx'a – Amkoe
Ancestry
West
East
South
0 10 20 30 40 50
0 5 10 15 20 25
0 5 10 15 20 25
